# Synthetic analogue of adrenocorticotropic hormone, ACTH_(4-7)_PGP delays neurological manifestations in diseases of mucopolysaccharidosis III spectrum by reducing neuroinflammation and rescuing neurotransmission, synaptogenesis, and axonal demyelination

**DOI:** 10.64898/2026.02.20.707013

**Authors:** Travis Moore, Patricia Dubot, Gustavo Viana, Poulomee Bose, Erjun Zhang, Behzad Nasseri, Xuefang Pan, Derek Norman Robertson, Lara Michele Feulner, Mahsa Taherzadeh, Patrick Piet van Vliet, Éric Bonneil, Shaukat Khan, Longguo Zhang, Francesco Attanasio, Srikanth Singamsetty, Thomas M. Durcan, Shunji Tomatsu, Pierre Thibault, Carlos R. Morales, Graziella Di Cristo, Gregor Andelfinger, Graciela Piñeyro, Jannic Boehm, Gregory A. Lodygensky, Jill Wood, Alexey V. Pshezhetsky

**Affiliations:** CHU Sainte-Justine Research Center, Université de Montréal, Montreal, H3T 1C5, QC, Canada; Department of Anatomy and Cell Biology, McGill University, Montreal, H3A 0C7, QC, Canada; Proteomic Platform, Institute for Research in Immunology and Cancer, Université de Montréal, Montreal, H3T 1J4, QC, Canada; Nemours/Alfred I. duPont Hospital for Children, Wilmington, DE, USA; Institute of Crystallography, National Council of Research, Catania, Italy; Phoenix Nest Inc., Brooklyn NY, USA; The Neuro’s Early Drug Discovery Unit (EDDU), McGill University, Montreal, QC, Canada; Department of Neurosciences, Université de Montréal, H3C 3J7, QC, Canada

## Abstract

Mucopolysaccharidosis III (MPS III or Sanfilippo disease) is a spectrum of 4 genetic disorders (MPS IIIA-D), caused by defects in the genes *SGSH, NAGLU, HGSNAT* and *GNS* encoding enzymes involved in degradation of heparan sulfate (HS). HS accumulates in brain tissues and causes neuronal dysfunction and neurodegeneration leading to neuropsychiatric problems, developmental delays, childhood dementia, blindness and death during the second decade of life.

Previously, we demonstrated that pathophysiological mechanisms, underlying MPS IIIC in mouse models, involves functional pathological changes, affecting synaptogenesis and synaptic transmission and leading to learning and memory deficits. These results suggested that a treatment for MPS III could be developed by using compounds inducing synaptogenesis. In the current study, we tested the efficacy of a synthetic peptide ACTH_(4-7)_PGP, an analog of adrenocorticotropic hormone fragment, previously used as a neuroprotective and anti-inflammatory medication for treatment of acute neurological conditions, including stroke. We show that intranasal administration of ACTH_(4-7)_PGP restores defective synaptic transmission in CA1 pyramidal neurons of MPS IIIA and MPS IIIC mouse models and rescues the decrease in synaptic proteins in cultured MPS IIIC mouse hippocampal neurons and iPSC-derived neurons of human MPS IIIA, MPS IIIB and MPS IIIC patients. Furthermore, daily intranasal administration of ACTH_(4-7)_PGP to MPS IIIC and MPS IIIA mice reduces hyperactivity and rescues defects in working and spatial memory, delays progression of CNS pathology including neuroinflammation and axonal demyelination, and increases the lifespan. Together with the absence of any adverse reactions to ACTH_(4-7)_PGP in the MPS III and WT mice, our results justify testing the drug’s efficacy in clinical settings.

## Introduction

Mucopolysaccharidosis III or Sanfilippo disease is a spectrum of neurological progressive childhood diseases caused by genetic mutations in four enzymes involved in lysosomal degradation of a glycosaminoglycan, heparan sulfate (HS). Accumulation of HS oligomers in the CNS leads to progressive neurological deterioration manifesting with behavioral abnormalities and cognitive decline, followed by childhood dementia and death in the second decade of life [1], (reviewed in [2–4]). Peripheral manifestations are generally mild and include joint stiffness and limited range of joint mobility, recurrent rhinitis, hearing loss, coarse facial features and hepatosplenomegaly reviewed in [2–4]. Currently, no specific treatments are approved for Sanfilippo disease. In the brains of MPS III patients and mouse models, HS is primarily accumulated in phagocytic microglia and astrocytes, triggering their activation and release of pro-inflammatory cytokines and chemokines [5,6], which can further stimulate the transendothelial migration of monocytes from the blood into the parenchyma resulting in further damage (reviewed in [3]).

In the neurons, storage of HS leads to impaired vesicular trafficking, including that of synaptic vesicles, resulting in reduction of their density in synaptic terminals. Together with a drastic decrease in the density of mature dendritic spines, which are the preferential sites of glutamatergic synapse localization, and smaller areas of post-synaptic densities, this leads to defects in synaptic transmission. In particular, drastically reduced frequencies/amplitudes of excitatory miniature synaptic currents in pyramidal CA1 hippocampal neurons of MPS IIIC mice were observed as early as at P15-P20, 2-3 months before the development of other neuronal pathologies [7–9]. At later stages of the disease progression, blocked autophagy/mitophagy contributes to secondary accumulation of toxic amyloid materials, changes in the neuronal membrane lipid composition and reduction in the number of healthy mitochondria in the neurons, overall, leading to pronounced neurodegeneration[10–12]. Similar processes in oligodendrocytes cause a reduction in the number of mature functionally active cells, thereby contributing to axonal demyelination and white matter defects [13]. Finally, our recent data suggested that lysosomal HS storage in the neurons of Sanfilippo patients and mouse models causes the secondary deficiency of lysosomal neuraminidase 1, which in turn, alters the sialylation of proteins at the neuronal surface thought to be essential for synaptogenesis, memory consolidation, and behaviour processes [14].

Several treatments for MPS III aimed at correcting the root cause of the disease, the genetic deficiency of the HS-degrading enzymes, through an intrathecal delivery of the replacement enzyme, AAV-mediated in vivo gene therapy, or LV-mediated HSPC gene therapy have shown good efficacy in mice and dogs [15–18]. However, clinical trials for these strategies failed to demonstrate clinical benefits in trial participants treated after the onset of neurological symptoms, reviewed in [19,20]. In most cases, the levels of HS in the cerebrospinal fluid of the patients were drastically reduced or even normalized suggesting that although the disease progression was stabilized, the damage to the CNS could not be reversed.

While further studies are necessary to define the exact mechanism underlying appearance of early and drastic synaptic defects in Sanfilippo patients and mouse models, the results discussed above provide a strong rationale for testing if drugs known to enhance excitatory synaptic transmission and reduce neuroinflammation can rescue behavioral and cognitive defects in animal models of MPS III. One of such compounds is the synthetic analogue of adrenocorticotropic hormone 4-10 heptapeptide, consisting of an ACTH fragment, ACTH_(4–7)_ and the C-terminal tripeptide Pro-Gly-Pro (ACTH_(4-7)_PGP; M-E-H-F-P-G-P). This peptide has been previously shown to promote the survival of neurons during hypoxia[21] and glutamate neurotoxicity[22]. In contrast to ACTH_(4-10)_, ACTH_(4-7)_PGP does not have hormonal activity[23], but shows neuroprotective properties, inhibits nitric oxide synthesis[24], induces production of nerve growth factor (NGF) and brain-derived neurotrophic factor (BDNF)[25,26]. After intranasal administration at a dose of 50 µg/kg BW in rodents [23,27–29] and in humans [30], ACTH_(4-7)_PGP was shown to be readily targeted to the brain and to enhance memory and learning within 24 h after treatment. Besides, clinical trials demonstrated that after intranasal administration, ACTH_(4-7)_PGP improved recovery in human patients with ischemic stroke and diseases of the optic nerve [31,32], which led to its approval for clinical use in some countries under the commercial name of Semax [31,32].

In the current study, we demonstrate that the N-terminally acetylated form of ACTH_(4-7)_PGP restores miniature events and evoked excitatory postsynaptic currents in pyramidal CA1 neurons in brain slices from MPS IIIA (*Sgsh^mps3a^*) and IIIC (*Hgsnat^P304L^*) mouse models. The drug also reverses the decrease in synaptic protein levels in cultured MPS IIIC mouse hippocampal neurons and in iPSC-derived cortical neurons of human MPS IIIA, MPS IIIB and MPS IIIC patients. Furthermore, daily intranasal administration of this peptide rescues behavioural defects in MPS IIIA and MPS IIIC mice and delays progression of brain pathology.

## Results

### 1. ACTH_(4-7)_PGP increases reduced levels of synaptic protein markers in cultured neurons from MPS IIIC mice and in iPSC-derived cultured cortical neurons of human MPS IIIA, MPS IIIB and MPS IIIC patients

Previously, we demonstrated that the density of glutamatergic synapses in cultured hippocampal neurons of two MPS IIIC mouse models, *Hgsnat-Geo* constitutive knockout strain and *Hgsnat^P304L^* strain with debilitating missense HGSNAT mutation, was reduced compared to those from control mice, by quantifying the colocalization between the presynaptic marker VGLUT1 and the postsynaptic marker, PSD-95 [8,33]. In order to test whether ACTH_(4-7)_PGP is capable of preventing these deficits, we treated cultures of hippocampal neurons from the *Hgsnat^P304L^* mice with ACTH_(4-7)_PGP at a concentration of 10 µM previously shown to be physiologically active in the cultured neurons [21][22]. We opted for using the N-terminally acetylated form of the peptide which could potentially reduce its degradation by aminopeptidases and increase stability in tissues. The peptide was added to the culture media when the neurons were plated and, further, when 50% of the media was changed every 3 days. At 21 days *in vitro* (DIV21), neurons were fixed and analysed by immunohistochemistry (ICC) to assess levels of markers of dendrites (MAP2), axons (medium chain of neurofilament protein, NF-M), and glutamatergic synapses (SYN1, VGLUT1, and PSD-95). We, also, analysed levels of BDNF using antibodies previously validated in *Bdnf* KO mice [34]. SYN1+ and BDNF+ puncta, as well as VGLUT1+ puncta in juxtaposition with PSD-95+ puncta were quantified over 20 μm-long segments of neurites randomly selected at >30 μm distance from the soma. Our results indicate that while untreated hippocampal neurons of *Hgsnat^P304L^* mice show drastic reduction of BDNF+, SYN1+, and VGLUT1+ puncta in juxtaposition with PSD-95+ puncta compared to WT cells, the *Hgsnat^P304L^* neurons treated with ACTH_(4-7)_PGP do not display these deficits (**Figure S1**).

To test whether reduced levels of synaptic markers and the effect of the drug were recapitulated in neurons of human patients with different subtypes of MPS III, we have generated iPSC lines from cultured primary skin fibroblasts of six (2 MPS IIIA, 2 MPS IIIB and 2 MPS IIIC) patients and two healthy controls (CNTs) received from cell depositories or obtained with the consent of the families. All newly generated iPSCs lines were positive for pluripotency markers TRA-1-60, SOX2, SSEA4, and OCT4 (**Figure S2A**), and demonstrated the ability to differentiate *in vitro* into the three germ layer cells (**Figure S2B**). After confirmation of the primary enzymatic defects (**Figure S2C**), the iPSCs were differentiated into forebrain committed neural precursor cells (NPC) by dual SMAD inhibition. NPCs induced in cortical neuronal induction media (DMEM/F12) for 3 weeks in a monolayer [35] expressed neuronal markers, axonal β-tubulin III (clone TUJ1) and SYN1 (**Figure S3A**), along with NPC-specific markers, Nestin and PAX6 (**Figure S3B**). SGSH, NAGLU, or HGSNAT deficiencies in generated NPC lines were confirmed by measuring the corresponding enzyme activities in cell homogenates (**Figure S4A**). Consistent with the lysosomal storage phenotype, increased size/abundance of LAMP2+ puncta was detected in both MPS IIIB NPC lines compared to controls, while MPS IIIA and MPS IIIC cells showed a trend towards an increase (**Figure S4B-C**). Subsequently, the cortical NPCs were differentiated to cortical neurons by replating and culturing in a neurobasal media (NB) as described [36]. Neurons were cultured for 4 weeks (DIV28), in the absence or presence of 10 µM ACTH_(4-7)_PGP added during the media changes, before being fixed and labelled for VGLUT1, PSD-95, VGAT, gephyrin, NeuN and the lysosomal marker, LAMP2. The cells were also labeled with previously validated monoclonal antibodies against BDNF [34]. All iPSC-derived neurons of MPS III patients revealed primary enzyme deficiencies (**Figure S5A**). They also showed a significant increase in LAMP2 staining consistent with the lysosomal storage phenotype, which was not affected by the presence of ACTH_(4-7)_PGP in the culture medium (**Figure S5B-C**).

Notably, all iPSC-derived cortical MPS III neurons showed significantly reduced levels of BDNF+, VGLUT1+, PSD-95+, VGAT+ and gephyrin+ puncta (**Figure 1**). This result suggested that the deficiencies of protein markers of excitatory glutamatergic and inhibitory GABAergic synapses, previously observed in cultured neurons from MPS IIIC *Hgsnat^P304L^* mice [33], were recapitulated in human MPS IIIA, MPS IIIB and MPS IIIC cells (**Figure 1**). In contrast, all MPS III neurons treated with ACTH_(4-7)_PGP showed increased density of both excitatory and inhibitory synaptic proteins compared to untreated cells (**Figure 1A-D**). In most cases, the levels of synaptic proteins in treated MPS III neurons were similar to those observed in control cells (**Figure 1A-D**). The changes in the density of synaptic puncta in the ACTH_(4-7)_PGP-treated MPS III cells coincided with the increase in the density of BDNF+ puncta which was restored to the levels of the healthy control cells (**Figure 1C-D**).

**Figure 1.**
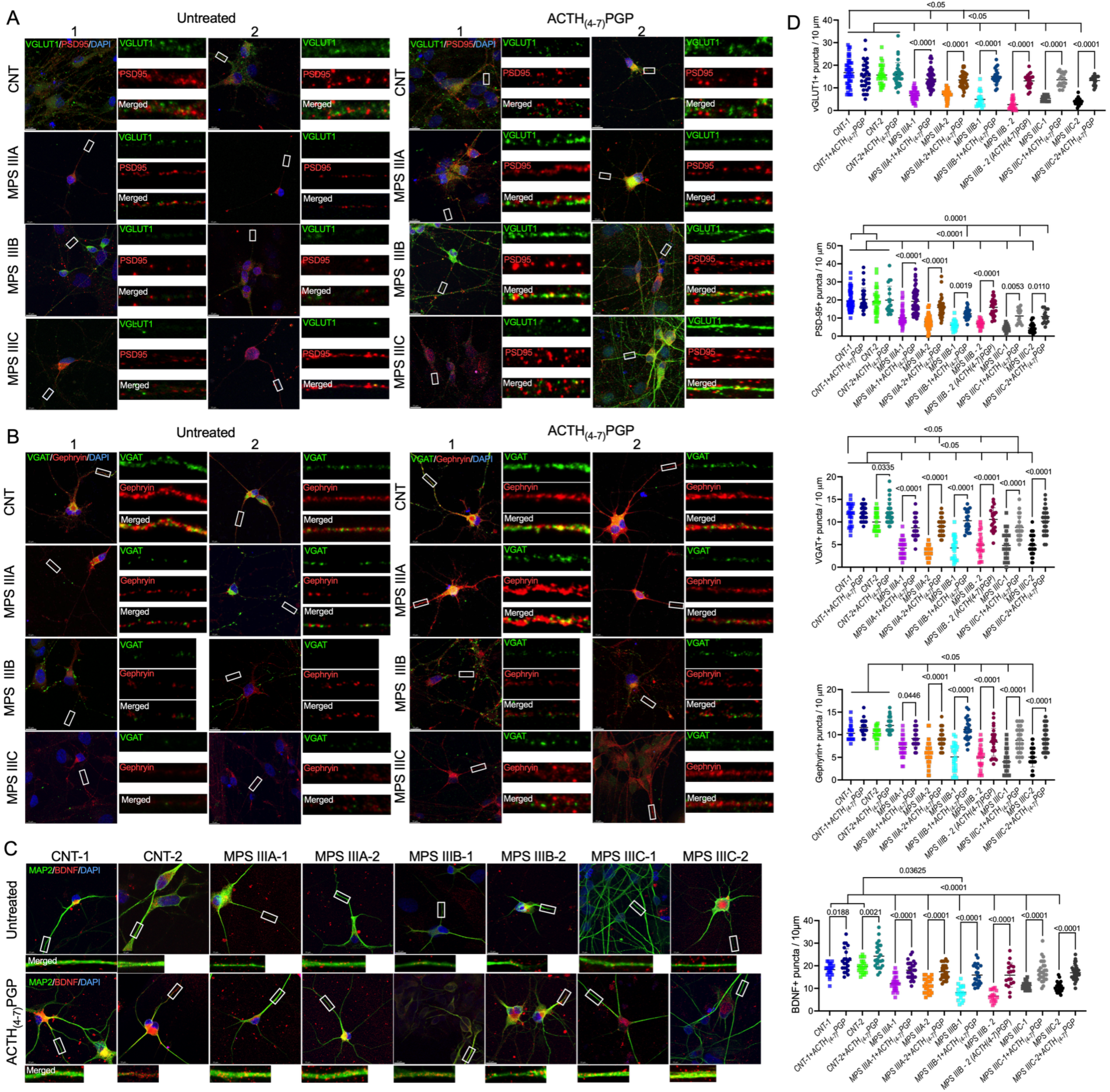
ACTH_(4-7)_PGP increases reduced levels of synaptic protein markers and BDNF in DIV28 iPSC-derived cultured cortical neurons from human MPS IIIA, MPS IIIB and MPS IIIC patients. **(A)** Density of markers of the excitatory synapse, VGLUT1+ and PSD95+ puncta, are reduced in neurons of MPS III patients and increased by ACTH_(4-7)_PGP treatment. Panels show representative images of MPS IIIA, IIIB, and IIIC neurons and those of healthy controls (CNT-1, and CNT-2) treated or not with ACTH_(4-7)_PGP and labeled for VGLUT1 (green) and PSD95 (red). DAPI (blue) was used to label nuclei. The scale bars equal 10 µm. Inserts show enlarged images of neurites selected at ≥10 µm from the soma (white rectangles). **(B)** Density of markers of the inhibitory synapse, VGAT+ and Gephyrin+ puncta, are reduced in neurons of MPS III patients and increased by ACTH_(4-7)_PGP treatment. Panels show representative images of MPS IIIA, IIIB, IIIC neurons and those of healthy controls (CNT-1, and CNT-2) treated or not with ACTH_(4-7)_PGP and labeled for VGAT (green) and Gephyrin (red). DAPI (blue) was used to label nuclei. The scale bars equal 10 µm. Inserts show enlarged images of neurites selected at ≥10 µm from the soma (white rectangles). **(C)** Levels of BDNF+ puncta are reduced in neurons of MPS III patients and increased by ACTH_(4-7)_PGP treatment. Panels show representative images of MPS IIIA, IIIB, IIIC neurons and those of healthy controls (CNT-1, and CNT-2) treated or not with ACTH_(4-7)_PGP and labeled for BDNF (red) and neuronal dendrite marker MAP2 (green). DAPI (blue) was used to label nuclei. The scale bars equal 10 µm. Inserts show enlarged images of neurites selected at ≥10 µm from the soma (white rectangles). **(D)** The graphs show quantification of BDNF+, VGLUT1+, PSD95+, VGAT+, and Gephyrin+ puncta along 10-µm segments of neuronal projections by ImageJ software. Individual values, means and SD from 3 biological replicates (>15 cells in each experiment) are shown. P-values were calculated using one-way ANOVA and Tukey post hoc test.

### 2. Intranasal administration of ACTH_(4-7)_PGP rescues synaptic transmission defects and ameliorates behaviour deficits

We hypothesised that defects in synaptic neurotransmission present in MPS III mouse models [7,8,37,38] could be rescued by ACTH_(4-7)_PGP previously shown to induce synaptogenesis [25,26]. We have selected intranasal administration as the primary delivery route in the MPS III mouse models, because it is non-invasive and used in clinical trials for short synthetic peptide drugs.

Peptide biodistribution and stability was evaluated in tissues of P45 MPS IIIC (*Hgsnat^P304L^*) mice by measuring ACTH_(4-7)_PGP levels in brain parenchyma, blood and visceral organs (liver, kidney, spleen) at different time points (10 min - 24 h) after a single intranasal dose of 50 µg/kg BW. The entire mouse brain was cut in four sections, rostral to caudal and assessed for biodistribution of ACTH_(4-7)_PGP by targeted LC-MS/MS using parallel reaction monitoring. For quantification, samples were spiked with isotopically labeled (Phe U-^13^C_9_; U-^15^N) ACTH_(4-7)_PGP peptide as an internal standard. These experiments demonstrated that 1 h after intranasal administration the concentration of the peptide in the brain (1.8-4.2 μg/kg) was higher than in blood plasma or visceral organs and remained above the previously estimated neuroprotective concentration [23,27–29] for at least 24 h after administration (**Figure 2A-B**).

**Figure 2.**
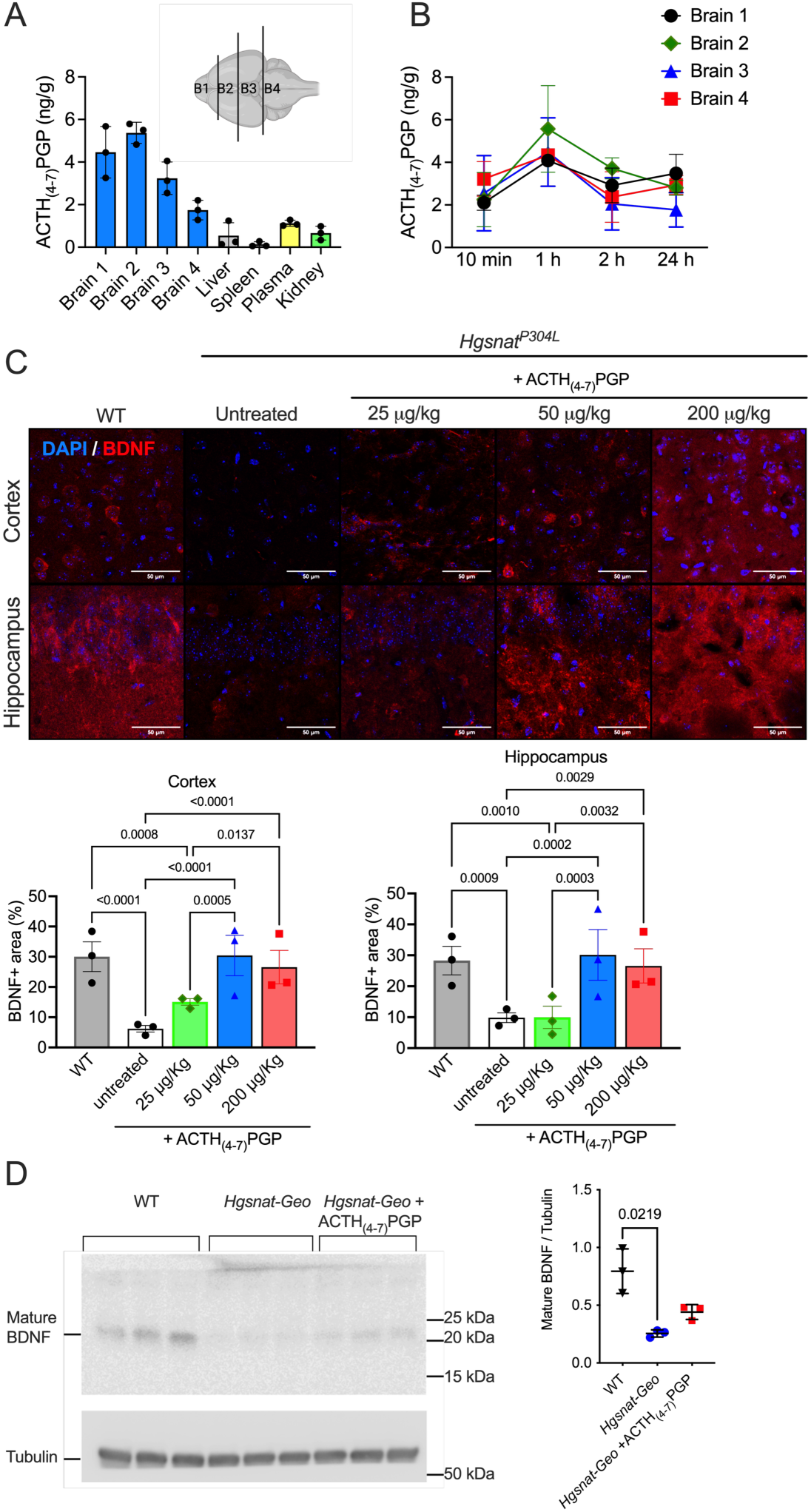
Pharmacokinetics and pharmacodynamics of ACTH_(4-7)_PGP after intranasal administration in MPS IIIC mice. **(A-B)** Active concentration of ACTH_(4-7)_PGP in the brain is achieved 1 h after intranasal administration and is sustained for 24 h. **(A)** LC-MS/MS MRM analysis shows effective delivery of ACTH_(4-7)_PGP to the brain 1 h after intranasal administration in MPS IIIC mice. Higher concentrations were detected in the rostral half (B1, B2) of the brain and tapering to the caudal regions. The drug via the systemic circulation (plasma) reaches visceral organs. Graphs show means and SD (n=3). **(B)** ACTH_(4-7)_PGP concentration in the four regions of the brain of MPS IIIC mice is sustained for 24 h after intranasal dosing. Graphs show individual results (ng of peptide/g of tissue weight), means and SD (n=3). **(C)** ACTH_(4-7)_PGP induces BDNF expression in a dose-dependent manner. Panels show typical images of brain cortices and CA1 hippocampi regions immunolabeled for BDNF (red). Nuclei are stained with DAPI (blue). Size bars are equal to 50 μm. Graphs show quantification of BDNF-positive areas (%) by ImageJ software. Individual data, means and SD (*n*=3-4) are shown. **(D)** Mature BDNF levels are reduced in the hippocampi of saline-treated 4-month-old MPS IIIC *Hgsnat*-*Geo* mice as compared to WT mice, and show a trend towards increase after a 10-day treatment with ACTH_(4-7)_PGP at the dose of 50 µg/kg BW/day. *P* values are calculated by ANOVA with Tukey post hoc test **(C)** or Kruskal-Wallis test with Dunn’s post hoc test **(D)**.

We further conducted pharmacodynamics (PD) studies, where we analyzed BDNF levels using immunofluorescent microscopy in the hippocampus and somatosensory cortex of P45 MPS IIIC mice after intranasal treatment with 0, 25, 50, and 200 µg/kg BW/day of ACTH_(4-7)_PGP for 15 consecutive days (**Figure 2C**). Untreated WT mice of the same age were used as controls. We found that while BDNF levels in *Hgsnat^P304L^* mice were reduced compared to WT controls, ACTH_(4-7)_PGP treatment at 50-200 µg/kg BW rescued BDNF deficits. The results of the immunohistochemistry experiments were confirmed by immunoblotting experiments in 4-month-old MPS IIIC *Hgsnat-Geo* mice treated with ACTH_(4-7)_PGP for 15 consecutive days. While mature BDNF levels in hippocampal tissues of *Hgsnat-Geo* mice treated with saline were reduced compared to WT animals, BDNF levels in the animals treated with ACTH_(4-7)_PGP showed a trend towards an increase (**Figure 2D**).

Since both pharmacokinetics and pharmacodynamics experiments have confirmed that the optimal ACTH_(4-7)_PGP concentration and regimen for intranasal administration in mice was 50 µg/kg BW/day, we used these conditions to test the ACTH_(4-7)_PGP effect on synaptic transmission in hippocampal pyramidal neurons of MPS IIIC *Hgsnat^P304L^* mice and previously described [39] MPS IIIA (*Sgsh^mps3a^*) mice. We treated mice with the peptide for 30 consecutive days between P30 and P60 followed by the preparation of acute brain slices and whole-cell patch-clamp recordings of miniature excitatory postsynaptic currents (mEPSC) from hippocampal CA1 neurons. Acute brain slices of untreated WT mice were used as controls.

Compared to WT mice, untreated *Sgsh^mps3a^* and *Hgsnat^P304L^* mice showed reduced excitatory miniature amplitude, and *Sgsh^mps3a^* mice showed an additional reduction in miniature frequency (**Figure 3A and C**). This phenotype, previously reported by us as early as at P14-20 [7,8] and confirmed by other studies [37] was rescued in both MPS IIIA and MPS IIIC mice, treated with ACTH_(4-7)_PGP for 30 days (**Figure 3A and C**).

**Figure 3.**
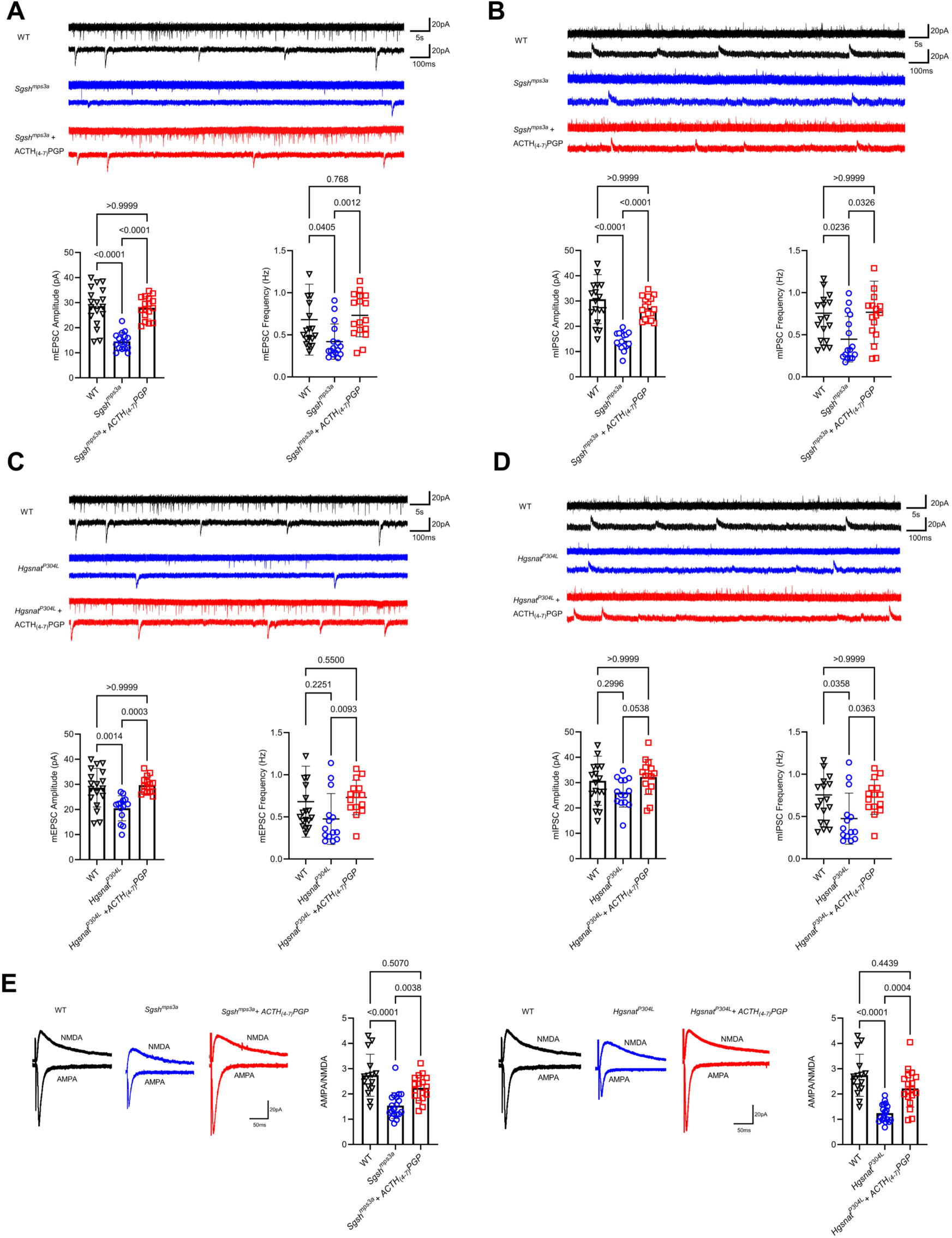
ACTH_(4-7)_PGP treatment rescues the synaptic phenotype in MPS IIIA and MPS IIIC mouse models. Whole-cell patch clamp recording of excitatory and inhibitory miniature events **(A-C)** and evoked AMPA-R vs NMDA-R responses **(E)** in hippocampal CA1 neurons in acute brain slices. **(A)** Excitatory miniature amplitude and frequency for WT, *Sgsh^mps3a^* and treated *Sgsh^mps3a^* mice. **(B)** Inhibitory miniature amplitude and frequency for WT, *Sgsh^mps3a^* and treated *Sgsh^mps3a^* mice. **(C)** Excitatory miniature amplitude and frequency for WT, *Hgsnat^P304L^* and treated *Hgsnat^P304L^* mice. **(D)** Inhibitory miniature amplitude and frequency for WT, *Hgsnat^P304L^* and treated *Hgsnat^P304L^* mice. **(E)** Ratio of evoked AMPA-R and NMDA-R responses for WT, *Sgsh^mps3a^* and treated *Sgsh^mps3a^* mice (left panel), and for WT, *Hgsnat^P304L^* and treated *Hgsnat^P304L^* mice (right panel). *N*=3-4 mice (15-18 cells) per group for miniature events and 3 mice (15-20 cells) per group for AMPA/NMDA recordings. *P* values are calculated by Kruskal-Wallis test with Dunn’s post hoc test.

Likewise, we observed a decrease in mIPSC amplitudes and frequences in *Sgsh^mps3a^* mice, which was rescued by treating mice with ACTH_(4-7)_PGP (**Figure 3B**). In untreated *Hgsnat^P304L^* mice, we observed reduced inhibitory miniature frequency (but not amplitude) compared to their WT counterparts which was also rescued by ACTH_(4-7)_PGP (**Figure 3D**). Notably, we also did not observe a significant difference between treated MPS III and WT mice.

Having obtained a strong indication that ACTH_(4-7)_PGP rescues reduced synaptic inputs in both *Sgsh^mps3a^* and *Hgsnat^P304L^* mice, we, next, analysed whether the synaptic glutamate receptor composition in these animals was altered as compared to WT mice. Two glutamate receptors play a critical role in excitatory synaptic transmission and synaptic plasticity: AMPA-receptors and NMDA-receptors. Hence, we patch-clamped CA1 neurons in acute slices and recorded evoked glutamatergic responses at −60 mV holding potential to analyze AMPA-R responses, and at +40 mV holding potential to assess the NMDA-R component. AMPA/NMDA-ratios were significantly reduced in both *Sgsh^mps3a^* and *Hgsnat^P304L^* mice, indicating altered evoked excitatory transmission (**Figure 3E**). Treatment of mice with ACTH_(4-7)_PGP for 30 days rescued this phenotype in both mouse models of Sanfilippo syndrome. Together with the ability of ACTH_(4-7)_PGP to induce levels of synapses in the cultured neurons, these results suggested that the drug rescues synaptic deficits in MPS III neurons *in vitro* and *in vivo*.

We further tested if ACTH_(4-7)_PGP can rescue neurobehavioral deficits associated with synaptic dysfunction in MPS IIIC mice. The ability of the peptide to improve short-term memory deficits in the symptomatic 4-month-old MPS IIIC *Hgsnat^P304L^* mice was tested using the Novel Object Recognition (NOR) test. ACTH_(4-7)_PGP was administered intranasally at 50 µg/kg BW either in a single dose 17 h prior to the test or for ten consecutive days before the experiment, the time sufficient for restoration of BDNF deficit (**Figure 2**). The saline treated *Hgsnat^P304L^* and, both, saline-treated and ACTH_(4-7)_PGP-treated WT groups were used as controls. We found that saline-treated MPS IIIC mice showed reduced discrimination index suggesting an impaired ability to recognize the familiar object (**Figure S6A**). The discrimination index for *Hgsnat^P304L^* mice, receiving the peptide for 10 consecutive days before the experiment was similar to those for the WT controls suggesting a rescue of the short-term memory deficit. However, a single dose of ACTH_(4-7)_PGP given 17 h before the test, did not improve behavior in the NOR test (**Figure S6A**) or in the YM test (**Figure S6B**). The single dose of the peptide, however, improved hyperactivity and increased impulsivity that *Hgsnat^P304L^* mice display during the early disease phase (4-5 months of age) [7]. Four-month-old *Hgsnat^P304L^* mice demonstrated ∼2-fold increase in the total distance traveled and the distance traveled in the center of the arena in the Open Field (OF) test compared to the saline-treated WT mice (**Figure S6C-D**). In *Hgsnat^P304L^*mice, treated with a single dose of 50 µg/kg BW of ACTH_(4-7)_PGP 17 hours before the experiment, the total distance traveled and the distance traveled in the center were reduced compared to untreated *Hgsnat^P304L^* mice indicating that the treatment partially rescued these behavioral traits. Ability of the peptide to rescue increased impulsivity was also tested in the Elevated Plus Maze test (EPM) that measures a natural fear of heights in rodents. Saline-treated *Hgsnat^P304L^* mice at 4 months showed increased time spent in the open arms and increased numbers of entries into the open arm compared to the WT animals, while *Hgsnat^P304L^* mice animals treated with a single dose of ACTH_(4-7)_PGP showed a behavior similar to their WT counterparts (**Figure S6E-F**).

### 3. Long-term treatment with ACTH_(4-7)_PGP delays neurobehavioral manifestations

We further conducted a preclinical efficacy study to test whether long-term administration of the peptide can delay clinical and pathological manifestations of the disease in MPS IIIA (*Sgsh^mps3a^*) and MPS IIIC (*Hgsnat^P304L^*) mouse models.

The study design is shown in **Figure S7**. In both models, treatment was started at P30. WT, *Hgsnat^P304L^* and *Sgsh^mps3a^* mice were randomly assigned to the treatment and control groups. The cohort size (∼18 mice/sex/treatment) was calculated based on mean variability of replicates in previous behavioral tests in *Hgsnat^P304L^* mice to detect a ∼40% difference between means (power=0.8). To test if long-term administration of the peptide results in major metabolic changes, the mouse body weight was measured weekly and no difference in body weight and body weight gain was detected between the treated and untreated *Sgsh^mps3a^ Hgsnat^P304L^* or WT mice (**Figure S8)**.

MPS IIIC (*Hgsnat^P304L^*) mice were treated between P30 and 5 months, when phenotypic differences between WT and *Hgsnat^P304L^* mice are fully pronounced according to our previous data [7]. Then, their behavior was assessed by OF (anxiety, impulsivity, fear and hyperactivity), EPM (anxiety, fear), Y-Maze (YM) and NOR (short-term memory) tests. In the OF test, both male and female *Hgsnat^P304L^* mice treated with saline showed significantly increased hyperactivity (increase in the total distance traveled) and impulsivity (increased time spent in the center of the arena) compared to the WT animals **(Figure 4A,B).** In contrast, *Hgsnat^P304L^* mice, treated with ACTH_(4-7)_PGP, showed an absence of these phenotypes **(Figure 4A,B)**. When studied by EPM test, 5-month-old female and male *Hgsnat^P304L^* mice, treated with saline showed increased impulsivity (increase in the number of entries into open arms) compared to WT animals **(Figure 4C).** In contrast, ACTH_(4-7)_PGP*-*treated *Hgsnat^P304L^* mice showed a reduction in the number of open arm entries compared to untreated *Hgsnat^P304L^* mice and no difference with their WT counterparts indicating normal levels of fear and impulsivity. In the YM test, saline-treated *Hgsnat^P304L^* mice but not ACTH_(4-7)_PGP-treated or WT mice showed a reduction of alternation index suggestive of short-term memory deficits (**Figure 4D**). In NOR test, *Hgsnat^P304L^* mice treated with saline showed a significant reduction in discrimination index suggesting a short memory deficit, while *Hgsnat^P304L^* mice treated with ACTH_(4-7)_PGP revealed an absence of this phenotype (**Figure 4E**). Together, all data demonstrated that daily intranasal treatment with ACTH_(4-7)_PGP prevented development of neurobehavioral deficits in the *Hgsnat^P304L^* mice.

**Figure 4.**
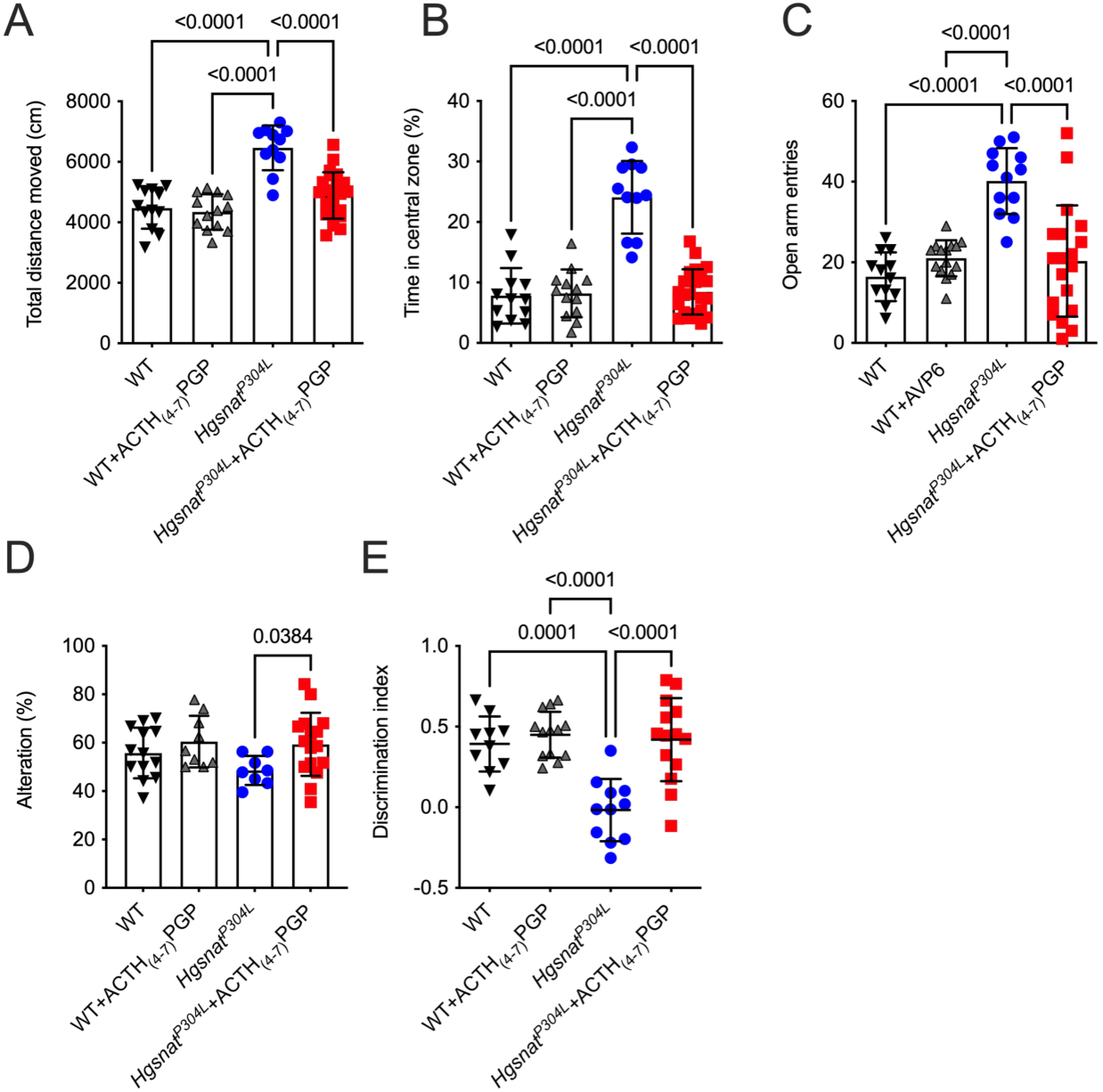
*Hgsnat^P304L^* mice treated with ACTH_(4-7)_PGP reveal delay in development of neurobehavioral abnormalities at the age of 5 months. Saline-treated *Hgsnat^P304L^* mice at the age of 5 months show increased total distance traveled in the OF test **(A)**, increased time spend in the central zone in the OF test **(B)**, and increased number of open arm entries in the EPM test, **(C)** suggestive of hyperactivity and increased impulsivity. They also show reduced alterations between arms in the Y-Maze test **(D)** and decrease in the discrimination index in the NOR test **(E)**, suggestive of spatial and short-term memory deficits. *Hgsnat^P304L^*mice treated daily, starting from P30, with ACTH_(4-7)_PGP show a rescue of all above deficits. Individual results, means and SD from experiments performed with 6-11 mice per genotype, per sex, per treatment are shown. P values were calculated using one-way ANOVA with Tukey post-hoc test.

While majority of *Hgsnat^P304L^* and WT mice were sacrificed at the age of 5 month after completion of behavioural tests for histopathological examination, the treatment was extended for a smaller group (3-5 mice sex/treatment/genotype) to test whether the peptide ameliorated behavioural abnormalities for a longer duration until the age of 8-9 months (previously determined to be the terminal stage of the disease for this strain [7]) and increased survival. Since no difference in behaviour was observed between male and female mice in the same group, the results for both sexes were pooled. The OF test, conducted at the age of 8 months, revealed reduced thigmotaxis (increased time spent in the center of the arena) and hyperactivity (increased total distance traveled) in untreated *Hgsnat^P304L^* mice. These traits were partially reversed in *Hgsnat^P304L^* mice treated with ACTH_(4-7)_PGP compared to *Hgsnat^P304L^* mice treated with saline (**Figure S9A-B)**. No difference in behaviour was observed between saline-treated and ACTH_(4-7)_PGP-treated WT mice. In the YM test, saline-treated *Hgsnat^P304L^* 8-month-old mice showed significantly reduced percent of alternation between arms compared to WT animals. The ACTH_(4-7)_PGP-treated *Hgsnat^P304L^* mice demonstrated an increase in alternation compared to saline-treated animals (**Figure S9C**).

For MPS IIIA *Sgsh^mps3a^* mice, behavior abnormalities were observed starting from 6 months. At this age, mice showed short memory deficits in the NOR test and reduced ability to associate an environment with a fear-inducing stimulus in the Contextual Fear Conditioning (CFC) test. They also demonstrated a reduced general sociability in Three-Chamber Sociability (TCS) test. On the other hand, OF tests conducted for *Sgsh^mps3a^* mice at the ages of 4 and 7 months did not reveal signs of hyperactivity or increased impulsivity (**Figure S10**) and, thus, this test has not been used for their evaluation. Based on these observations, MPS IIIA mice were treated with ACTH_(4-7)_PGP until the age of 7 months, and assessed with TCS, NOR, and CFC tests. They were then sacrificed for the analysis of tissue pathology.

Social behavior of ACTH_(4-7)_PGP-treated and vehicle-treated *Sgsh^mps3a^*and WT mice was evaluated using Crawley’s TCS protocol that assesses cognition in the form of general sociability. Both ACTH_(4-7)_PGP-treated and vehicle-treated WT mice spent significantly more time in the chamber with access to a never-before-met intruder mouse than in the identical empty chamber showing a normal sociability typical for rodents (**Figure 5A**). No preference for the chamber with an intruder over the empty chamber was detected in vehicle-treated *Sgsh^mps3a^* mice consistent with deficits in social behavior. In contrast, ACTH_(4-7)_PGP-treated *Sgsh^mps3a^* mice spent more time in the chamber with a stranger mouse compared to the empty chamber, indicating a rescue of impaired social motivation and affiliation (**Figure 5A**). Besides, unlike saline-treated or ACTH_(4-7)_PGP-treated WT mice, saline-treated *Sgsh^mps3a^* mice showed no difference in the interaction time with a familiar mouse and a stranger mouse, while ACTH_(4-7)_PGP-treated *Sgsh^mps3a^*mice spent more time with a never-before-met mouse and were similar in this respect to the WT mice (**Figure 5B**).

**Figure 5.**
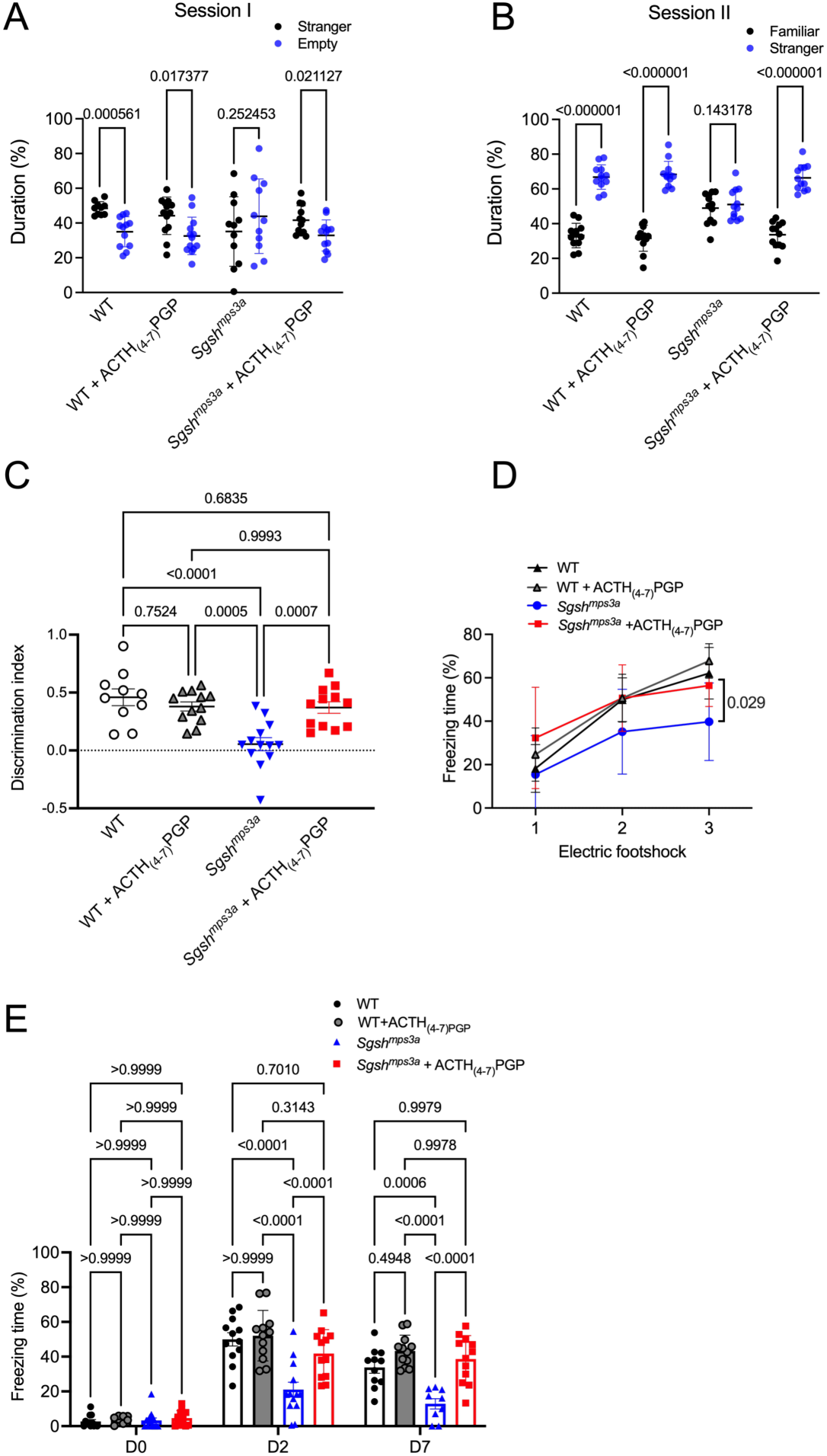
ACTH_(4-7)_PGP treatment rescues behavior abnormalities in *Sgsh^mps3a^* mice. Untreated and ACTH_(4-7)_PGP treated WT and *Sgsh^mps3a^* mice were analysed at 7 months of age for a reduced sociability (absence of an interest in a stranger mouse) in the TCS test **(A-B)**, short-term memory deficit (reduced discrimination index) in NOR test (**C**), and defects in associative learning (reduced freezing time) in the CFC test (**D**-**E)**. Individual results, means and SD of experiments performed with 10-12 male and female mice per genotype, per treatment are shown. *P* values were calculated using one-way **(C)** or two-way ANOVA **(A,B,D,E)** with Tukey post hoc test.

As in *Hgsnat^P304L^* mice, NOR revealed reduced values of a discrimination index for saline-treated *Sgsh^mps3a^* compared to WT mice consistent with deficits in short-term memory (**Figure 5C**). This deficit was absent in the *Sgsh^mps3a^* mice receiving ACTH_(4-7)_PGP treatment. In the CFC test, during the training day, saline-treated but not ACTH_(4-7)_PGP-treated *Sgsh^mps3a^* mice showed a decrease in freezing time after the third foot shock compared to WT mice (**Figure 5D**). Two and seven days after the end of the fear-training period, saline-treated *Sgsh^mps3a^* mice showed significantly reduced freezing time compared to WT mice revealing a problem associating the environment with the fear-inducing stimulus (**Figure 5E**). ACTH_(4-7)_PGP-treated *Sgsh^mps3a^* mice showed an increased freezing time compared to saline-treated *Sgsh^mps3a^* mice and similar to that of WT mice on the days 2 and 7 of the experiment consistent with a complete recovery of associative learning ability (**Figure 5E**).

### 4. Long-term treatment with ACTH_(4-7)_PGP prolongs survival and improves pathology

Untreated *Hgsnat^P304L^* mice develop severe urinary retention between the age of 8 and 9 months requiring euthanasia at the mean age of 42 weeks [7]. The mechanism underlying this phenotype, present also in other neurological MPS murine models including those of MPS IIIA and MPS IIIB, is not completely clear, but it was suggested to be caused by GAG storage and infiltration of immune cells in the epithelium of the urinary tract and the bladder wall [40]. To test whether the treatment delayed development of this phenotype, mice were examined for the signs of urinary retention daily, starting from the age of 7 months, and sacrificed, when abdominal distension was detected. We found that ACTH_(4-7)_PGP-treated *Hgsnat^P304L^* mice had a longer survival with the average life span of 49 weeks, 8 weeks longer than that of the saline-treated group (**Figure S11A**). When the wet weights of mouse spleens were measured at sacrifice, we found that the ACTH_(4-7)_PGP-treated *Hgsnat^P304L^* mice had drastically lower spleen weight than the saline-treated *Hgsnat^P304L^* mice despite being, on average, 8 weeks older (**Figure S11B**). Survival of MPS IIIA *Sgsh^mps3a^* mice was not evaluated, but ACTH_(4-7)_PGP-treated *Sgsh^mps3a^* mice, sacrificed at the age of 7 months, also showed wet weights of spleens (**Figure S11C**), significantly lower than that of saline-treated *Sgsh^mps3a^* mice and similar to that of WT mice, suggesting that the drug potentially ameliorated macrophage infiltration in some peripheral tissues.

After completing behavioural studies, brain tissues of ACTH_(4-7)_PGP-treated and saline-treated MPS III mice and of their matching WT controls were analyzed by immunofluorescence to assess the levels of previously described MPS III CNS biomarkers associated with synaptic defects, neuroinflammation and secondary storage/autophagy block. We found that the levels of presynaptic (VGLUT1) and postsynaptic (PSD-95) markers of glutamatergic synapse as well as the levels of BDNF puncta were reduced in the CA1 hippocampus area and the layers 4-5 of the somatosensory cortex of saline-treated *Hgsnat^P304L^*compared to WT mice (**Figure 6A-B**). In the brains of the ACTH_(4-7)_PGP-treated *Hgsnat^P304L^* mice, the levels of all 3 markers were increased compared to saline-treated animals, although they did not reach WT levels (**Figure 6A-B**).

**Figure 6.**
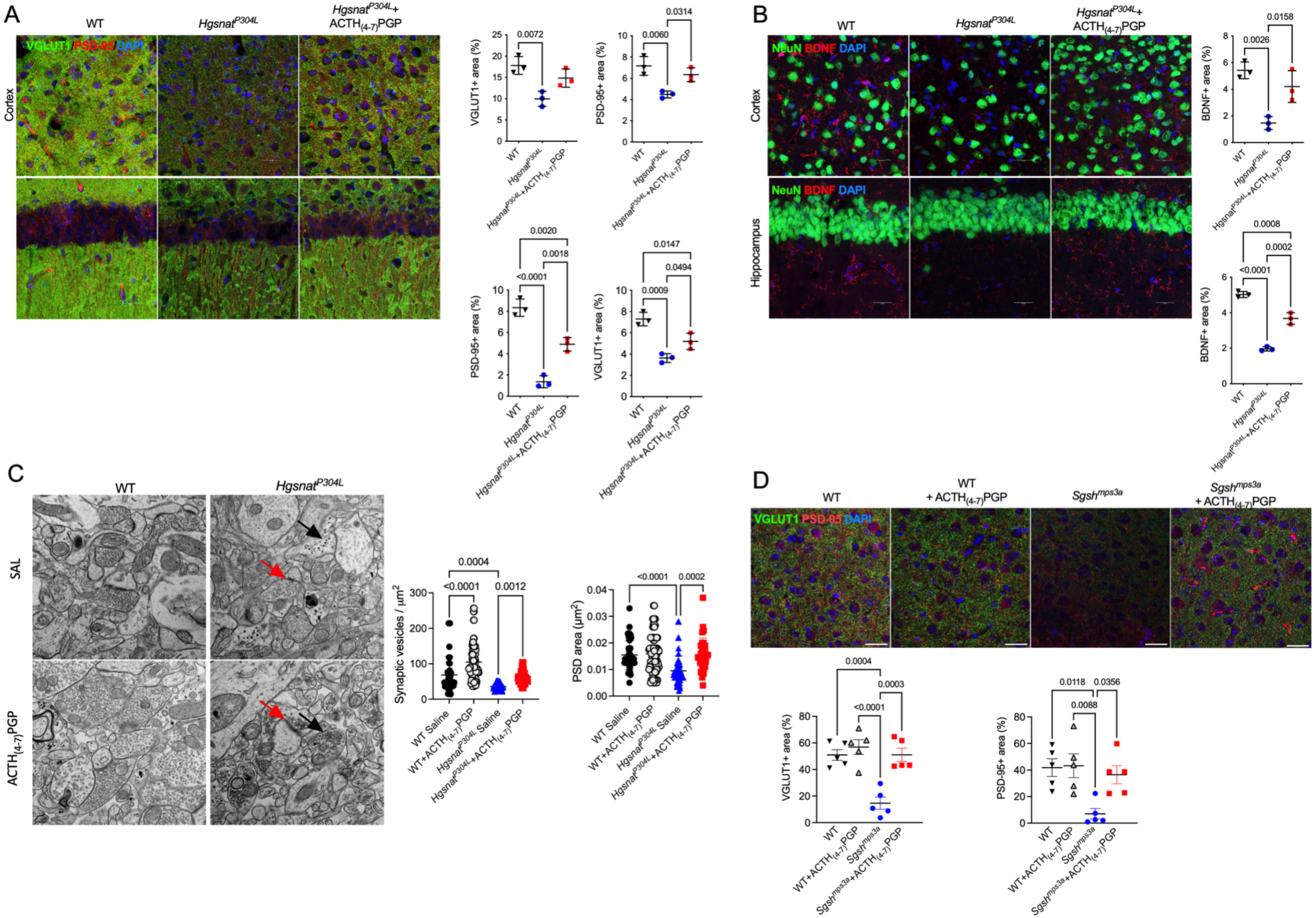
*Hgsnat^P304L^* mice treated with ACTH_(4-7)_PGP reveal partial rescue of synaptic protein markers and neuroinflammation at the age of 5 months. **(A-B)** Deficient levels of protein markers of glutamatergic synaptic neurotransmission, VGUT1 and PSD-95 **(A)** and BDNF **(B)** are rescued in the somatosensory cortex and hippocampus of *Hgsnat^P304L^* mice treated with ACTH_(4-7)_PGP. Panels show representative images of brain cortex (layers 4-5) and CA1 area of the hippocampus of 5-month-old WT and *Hgsnat^P304L^* mice, treated or not with ACTH_(4-7)_PGP. The tissues are stained with antibodies against PSD-95 (red) and VGLUT1 (green). **(A)**, BDNF (red) and NeuN (green) **(B)**. **(C)** TEM analysis confirms that synaptic defects in *Hgsnat^P304L^* mice are restored by ACTH_(4-7)_PGP. Panels show representative TEM images of excitatory synapses in CA1 hippocampal neurons of 6-month-old WT mice and *Hgsnat^P304L^* mice treated with saline (SAL) or ACTH_(4-7)_PGP, taken at 13000X magnification. Graphs show area (µm^2^) of PSDs and density of synaptic vesicles measured in asymmetric synapses of neurons from the CA1 region of the hippocampus. Synaptic terminals on the TEM images are marked with black and PSDs, with red arrows. Data show values, means, and SD of the results obtained with two mice per genotype (14 neurons per animal). *P* values were calculated by nested one-way ANOVA test with Tukey post hoc test. Scale bars equal 400 nm. **(D)** Deficient levels of VGUT1 and PSD-95 are rescued in the somatosensory cortex of *Sgsh^mps3a^* mice treated with ACTH_(4-7)_PGP. Panels show representative images of brain cortex (layers 4-5) of 7-month-old WT, and *Sgsh^mps3a^* mice, treated or not with ACTH_(4-7)_PGP. The tissues are stained with antibodies against PSD-95 (red) and VGLUT1 (green) In all confocal microscopy panels DAPI (blue) was used as a nuclear counterstain. Scale bars equal 25 µm. The graphs show quantification of fluorescence with ImageJ software. Individual results, means and SD from experiments performed with 3-5 mice per genotype (3 areas/mouse), per treatment are shown. *P* values were calculated using nested ANOVA with Tukey post hoc test.

Transmission Electron Microscopy (TEM) was further used to evaluate histopathological changes in the synapses of treated and untreated 6-month-old *Hgsnat^P304L^* mice at the ultrastructural level. TEM images of the CA1 region of the hippocampus taken at 13000x magnification (≥14 neurons/sample) were analyzed to measure the lengths and areas of PSD of excitatory asymmetric synapses, and the number of synaptic vesicles in synaptic terminals (**Figure 6C**). While the average PSD length was similar in all groups, the PSD areas were reduced by ∼50% in saline-treated *Hgsnat^P304L^* compared to WT mice and rescued in the ACTH_(4-7)_PGP-treated *Hgsnat^P304L^* mice (**Figure 6C**). The treatment also restored the reduced densities of synaptic vesicles in the terminals of excitatory synapses of *Hgsnat^P304L^* mice to the levels observed in WT mice (**Figure 6C**). Interestingly, synaptic vesicle densities were also increased in the ACTH_(4-7)_PGP-treated WT mice compared to saline-treated WT mice. Together, the results of TEM experiments confirmed at the ultrastructural level that the *Hgsnat^P304L^* mice have significant reductions of PSD areas and densities of synaptic vesicles in the terminals of excitatory synapses, and that ACTH_(4-7)_PGP treatment restores them to the normal levels. In similar fashion, in the cortices of saline-treated 7-month-old *Sgsh^mps3a^* mice, we observed a reduction in the levels of BDNF and synaptic markers, PSD-95 and VGLUT1 compared to WT controls. In the ACTH_(4-7)_PGP-treated *Sgsh^mps3a^* mice the levels of both proteins were increased compared to saline treated *Sgsh^mps3a^* mice and similar to those of WT mice (**Figure 6D**).

The numbers of GFAP+ astrocytes and CD68+ activated microglia, the markers of astrocytosis and microgliosis, respectively, were reduced in both cortices and hippocampi of ACTH_(4-7)_PGP-treated 6-month-old *Hgsnat^P304L^* mice compared to saline-treated *Hgsnat^P304L^* mice, suggesting that the drug decreased the neuroimmune response (**Figure 7A-B**). Numbers of GFAP^+^ and CD68^+^ cells were also significantly reduced in ACTH_(4-7)_PGP-treated mice compared to saline-treated *Hgsnat^P304L^* at the age of 10-11 months (**Figure S12A-B)**. The amelioration of neuroimmune response by ACTH_(4-7)_PGP treatment was further corroborated by the analyses performed in the brain tissues of 7-month-old MPS IIIA mice. Cortices of saline-treated 7-month-old MPS IIIA mice showed increase in GFAP+ activated astrocytes and CD68+ activated microglia compared to treated or untreated WT mice or *Sgsh^mps3a^*mice treated with ACTH_(4-7)_PGP (**Figure 7C**). Quantitative RT-PCR analysis also demonstrated reduction in the expression of proinflammatory cytokines, IL-1β, and TNF-α in the brain tissues of ACTH_(4-7)_PGP-treated treated *Hgsnat^P304L^*(**Figure 7D**) and *Sgsh^mps3a^* mice (**Figure 7E**) compared to untreated groups. In *Sgsh^mps3a^*mice ACTH_(4-7)_PGP also reduced expression of proinflammatory cytokine, Mip1α in the brain and circulating levels of Mip1α in the blood (**Figure 7E**).

**Figure 7.**
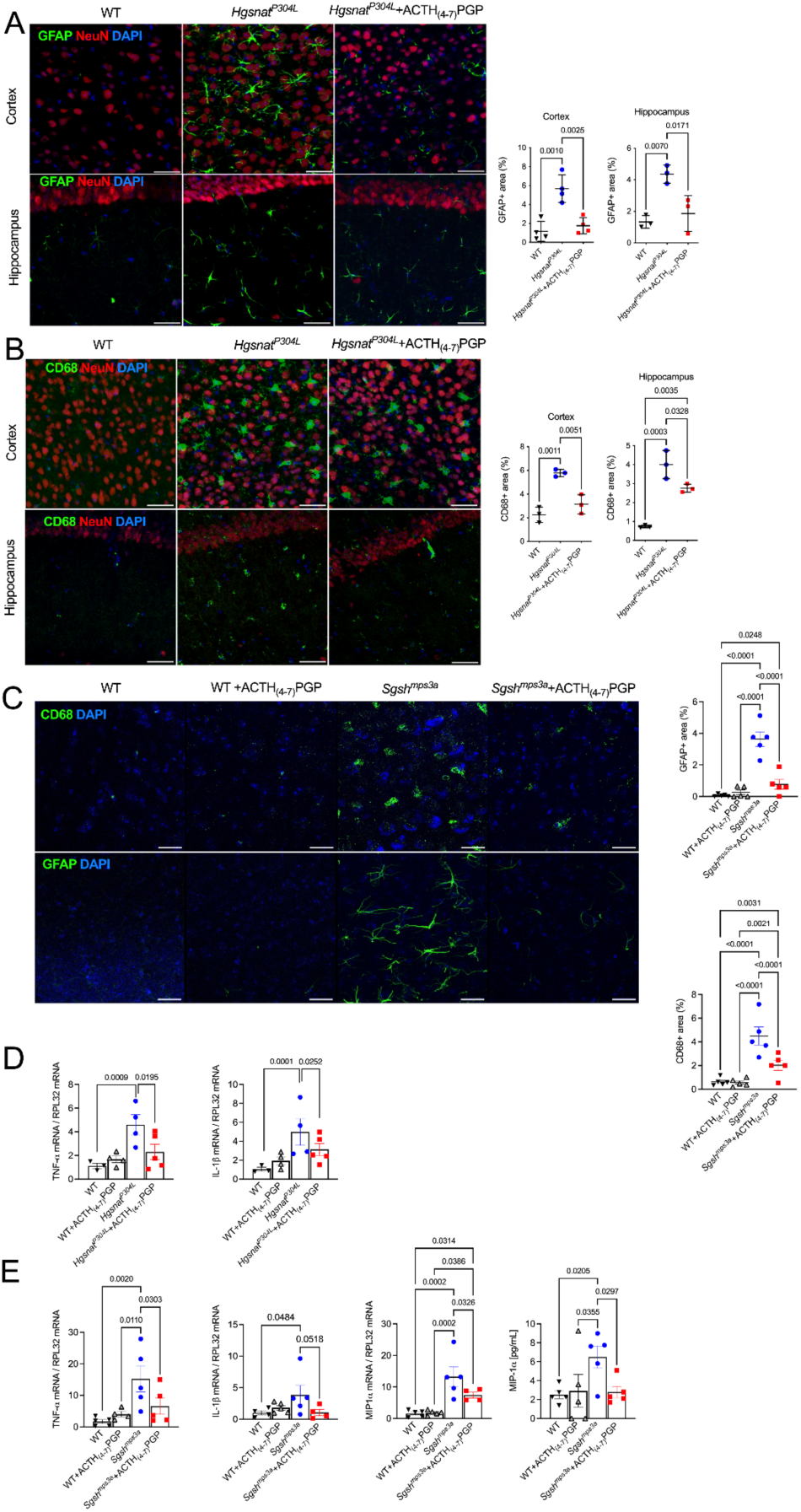
*Hgsnat^P304L^* and *Sgsh^mps3a^* mice treated with ACTH_(4-7)_PGP reveal partial rescue of neuroinflammation. **(A-C)** Levels of activated CD68+ microglia and GFAP+ astrocytes are reduced in the somatosensory cortex and hippocampus of *Hgsnat^P304L^* mice and in cortices of *Sgsh^mps3a^* mice treated with ACTH_(4-7)_PGP. Panels show representative images of brain cortex (layers 4-5) and CA1 area of the hippocampus of 5-month-old WT, and *Hgsnat^P304L^* mice, and cortex of 7-month-old WT and *Sgsh^mps3a^*mice treated or not with ACTH_(4-7)_PGP. The tissues are labeled with antibodies against GFAP (green) and NeuN (red) **(A),** or CD68 (green) and NeuN (red) **(B)** and CD68 (green) and GFAP (green) **(C)**. In all panels DAPI (blue) was used as a nuclear counterstain. Scale bars equal 25 µm. The graphs show quantification of fluorescence with ImageJ software. Individual results, means and SD from experiments performed with 3-5 mice per genotype (3 areas/mouse), per treatment are shown. P values were calculated using nested ANOVA with Tukey post hoc test. The mRNA expression levels of proinflammatory cytokines TNF-α and IL-1β measured by RT-PCR are increased in the brain tissues of untreated *Hgsnat^P304L^* **(D)** mice and *Sgsh^mps3a^* **(E)** mice compared to WT and ACTH_(4-7)_PGP-treated mice. **(E)** In *Sgsh^mps3a^* mice ACTH_(4-7)_PGP also reduces expression of Mip1α in the brain and circulating levels of Mip1α in the blood.

Previous research identified accumulation of unfolded proteins (including SCMAS, subunit C of the mitochondrial ATP synthase) as a hallmark of neuronal pathology in mouse MPS III models and brains of human patients [10]. These aggregates are drastically pronounced in the pyramidal layer IV-V neurons of somatosensory cortex and thought to be directly related to neurodegeneration and behavioural abnormalities [7]. To detect if this biomarker is altered by ACTH_(4-7)_PGP treatment, brain tissues were labeled with Thioflavin-S that stains misfolded proteins and with antibodies against SCMAS. We also assessed neuronal levels of GM2 ganglioside, and LC3+ puncta, a marker of impaired autophagy.

In the *Hgsnat^P304L^*mice, the ACTH_(4-7)_PGP treatment reduced secondary accumulation of GM2 ganglioside, SCMAS and Thioflavin-S+ protein aggregates in the cortical pyramidal neurons at 6 (**Figure 8A)** and 10-11 (**Figure S12C-E**) months of age. Similar result was observed for *Sgsh^mps3a^* mice at 7 months (**Figure 8B**). At the same time, the treatment did not change the brain levels of HS measured by LC-MS/MS (**Figure S13A**) or the levels of lysosomal storage marker, LAMP2-positive puncta **(Figure S13B**) suggesting that positive effects of ACTH_(4-7)_PGP on mouse behaviour and CNS pathology are not caused by a decrease in the lysosomal storage of HS.

**Figure 8.**
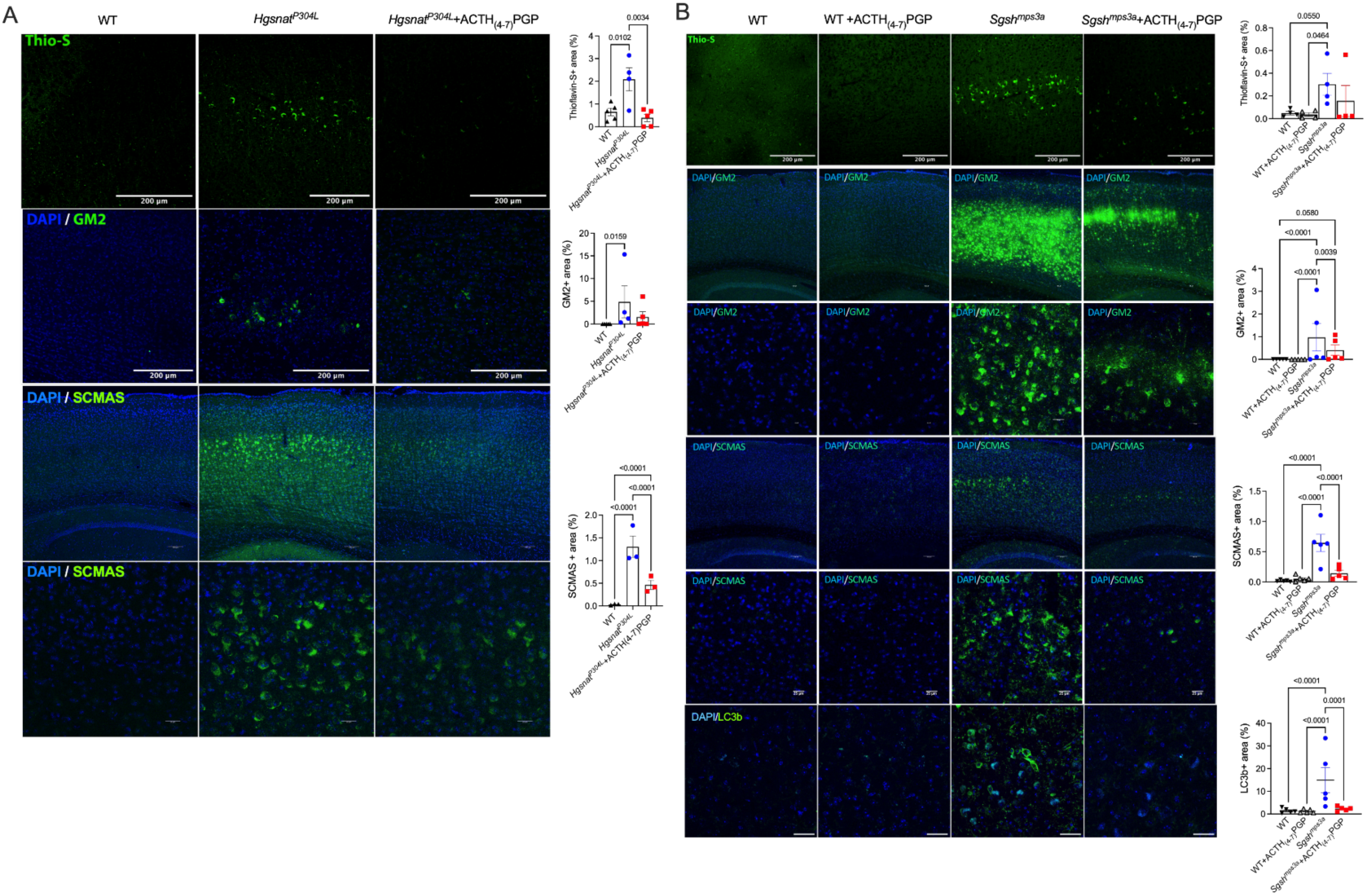
Secondary accumulation of GM2 ganglioside and misfolded proteins are reduced in brain cortices from *Hgsnat^P304L^* and *Sgsh^mps3a^*mice treated with ACTH_(4-7)_PGP. Panels show representative images of confocal laser scanning microscopy of brain somatosensory cortex (layers 4-5) from 5-month-old WT and *Hgsnat^P304L^* mice **(A)** and 7-month-old WT and *Sgsh^mps3a^* mice **(B)**, treated or not with ACTH_(4-7)_PGP, starting from P30. Brain sections were stained with Thioflavin-S or labeled with antibodies against G_M2_ ganglioside, unfolded subunit C of mitochondrial ATP synthase (SCMAS) or LC3b. DAPI was used as a nuclear counterstain. Graphs show quantification of fluorescence with ImageJ software. Individual results, means and SEM from experiments performed with 4-5 mice per genotype per treatment are shown. P values were calculated using nested one-way ANOVA with Tukey post hoc test. Scale bars: 200 μm, 100 μm or 25 μm.

Together, our results suggest that intranasal treatment with ACTH_(4-7)_PGP rescues pathological changes in the CNS of MPS IIIA and MPS IIIC mice.

### 5. Long-term treatment with ACTH_(4-7)_PGP halts axonal demyelination and pathological changes in the white matter

Previously, *ex vivo* Diffusion Tensor Imaging (DTI) revealed compelling signs of demyelination (26% increase in radial diffusivity) in corpus collosum of *Hgsnat^P304L^* mice [13]. Thus, DTI biomarkers were assessed in the brains of ACTH_(4-7)_PGP-treated and untreated WT, *Hgsnat^P304L^* and *Sgsh^mps3a^* mice to study the drug effect on the grey and white matter injury (**Figure 9A-C**). In the cortex (CX), DTI analysis identified a drastic increase in Mean diffusivity in both MPS IIIA (*Sgsh^mps3a^*) and MPS IIIC (*Hgsnat^P304L^*) mice, with an increase in Radial diffusivity in MPS IIIC (**Figure 9B**) suggesting reduced white matter integrity, or edema. Treatment restored Radial and Mean diffusivity in MPS IIIA and MPS IIIC as well as Axial diffusivity in MPS IIIA mice. In the hippocampus (HP), DTI analysis identified significant increase in Radial diffusivity and Mean diffusivity in both MPS IIIA and IIIC mice compared to WT. Therapy restored Radial and Mean diffusivity in both MPS IIIA and MPS IIIC mice and Axial diffusivity in type A only. In the corpus callosum (CC), DTI analysis identified an increase in Axial diffusivity in MPS IIIA *Sgsh^mps3a^* mice, and a normalization of Radial, Axial and Mean diffusivity values in treated compared to untreated *Sgsh^mps3a^* mice (**Figure 9C**). In the cerebral peduncle, we observed an increase in Mean diffusivity and Axial diffusivity in *Sgsh^mps3a^* mice and an increase in Radial diffusivity in *Hgsnat^P304L^* mice. Therapy resulted in a restoration of Axial diffusivity and Mean diffusivity in *Sgsh^mps3a^* model (**Figure 9C**). Together, our results suggest that the treatment restores white and grey matter integrity in both mouse models.

**Figure 9.**
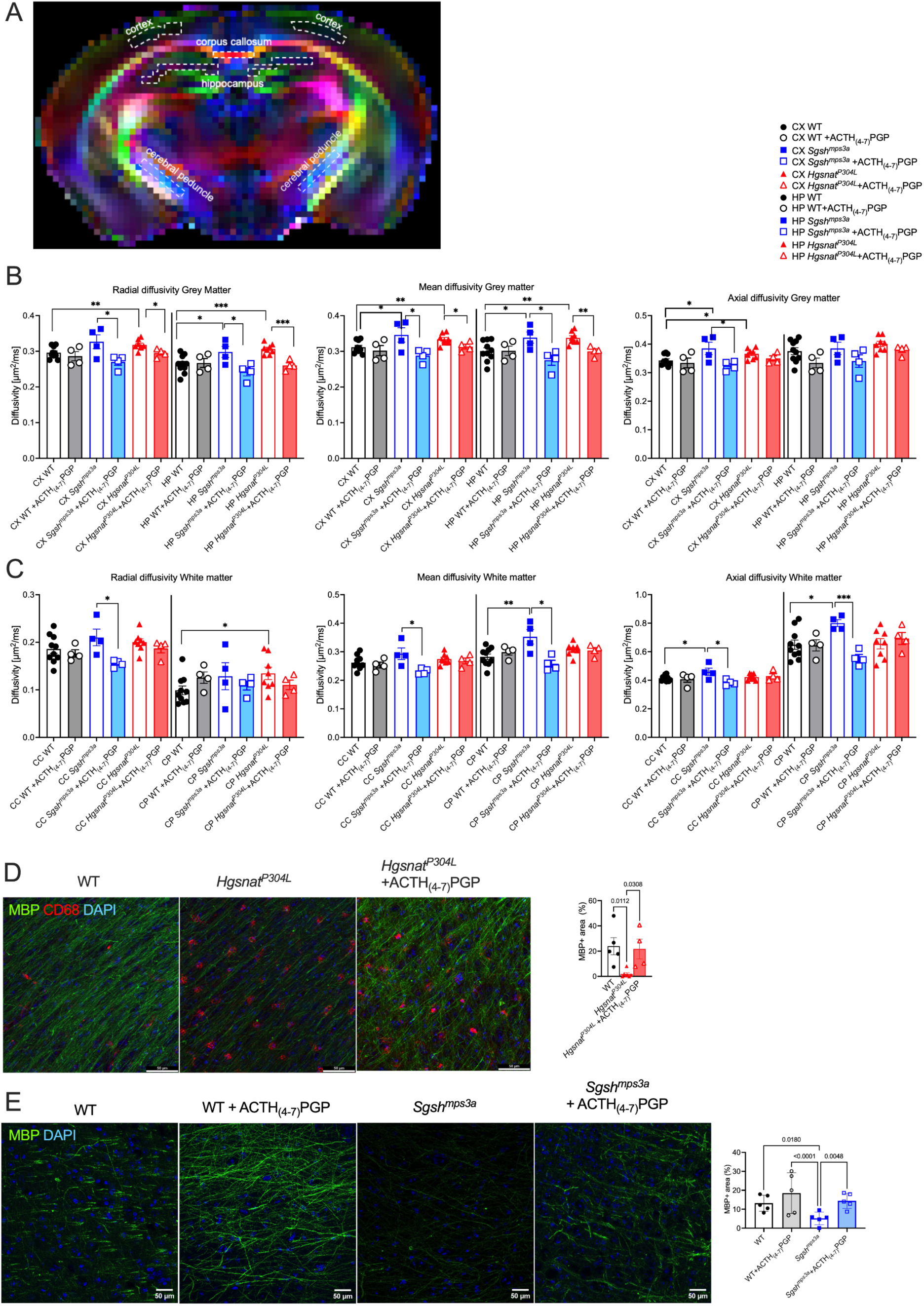
ACTH_(4-7)_PGP treatment protects MPS IIIA and MPS IIIC mice from axonal demyelination and grey/white matter injury. **(A-C)** Diffusion map **(A)** and metrics **(B-C)** from DTI for untreated or ACTH_(4-7)_PGP-treated WT, *Sgsh^mps3a^* MPS IIIA and *Hgsnat^P304L^* MPS IIIC 7-month-old mice. Bar graphs show Axial, Radial and Mean diffusion metrics from DTI of Cortex (CX), hippocampus (HP), corpus callosum (CC) and cerebral peduncle (CP). Abbreviations: AD, Axial diffusivity; RD, Radial diffusivity; MD, Mean diffusivity; Graphs show individual data, means and SEM. N=10-4 per group. P values were calculated by multiple unpaired t-test; * *P*<0.05; ** p<0.01; *** p<0.005. **(D-E)** Analysis of myelination in the brains of untreated and ACTH_(4-7)_PGP-treated WT, *Sgsh^mps3a^* MPS IIIA and *Hgsnat^P304L^* MPS IIIC 6-7-month-old mice. Panels show representative confocal microscopy images of CX areas of WT, untreated and ACTH_(4-7)_PGP-treated *Hgsnat^P304L^* mice **(C)** or untreated and ACTH_(4-7)_PGP-treated WT and *Sgsh^mps3a^* **(D)** mice labelled with antibodies against Myelin Basic Protein (MBP, green) and CD68 (red). DAPI (blue) was used as a nuclear counterstain. Scale bars equal 50 µm. Graphs show quantification of MBP areas. Individual values, means and SD are shown. N=4-3 per group. P values were calculated using nested ANOVA with Tukey post hoc test.

To confirm the results of DTI, we analyzed by immunofluorescence the levels of Myelin Basic Protein (MBP) in coronal sections of the same mouse brains after completion of DTI. Untreated 7-month-old *Sgsh^mps3a^* and *Hgsnat^P304L^* mice had substantially reduced MBP levels compared to their WT counterparts. The brains of ACTH_(4-7)_PGP-treated *Hgsnat^P304L^* and *Sgsh^mps3a^* mice showed a substantial increase in the levels of MBP compared to untreated mice suggesting that the treatment preserved myelination of axons (**Figure 9D-E**).

### 6. Single-cell RNA sequencing reveals reduction of neuroinflammatory response, and induction of neuronal synaptogenesis in treated *Hgsnat^P304L^* mice

To study the mechanism of the drug action *in vivo*, we performed a single-cell RNA sequencing (scRNA-seq) analysis of dissected hippocampi of WT and *Hgsnat^P304L^* mice treated with ACTH_(4-7)_PGP or saline for two weeks (between P27 and P42). The age and the duration of treatment were selected to observe the early effects of the drug in young mainly presymptomatic animals while also obtaining the cells with the higher viability, which was not possible when using older mice. For each condition, we analyzed 10,000 dissociated brain cells obtained from 2 male and 2 female mice. We identified 19 brain cell clusters based on the expression of 25 cell type-specific genes (**Figure 10A** and **Figure S14**). The structure of clusters was maintained across the 4 conditions (**Figure S15A**).

**Figure 10.**
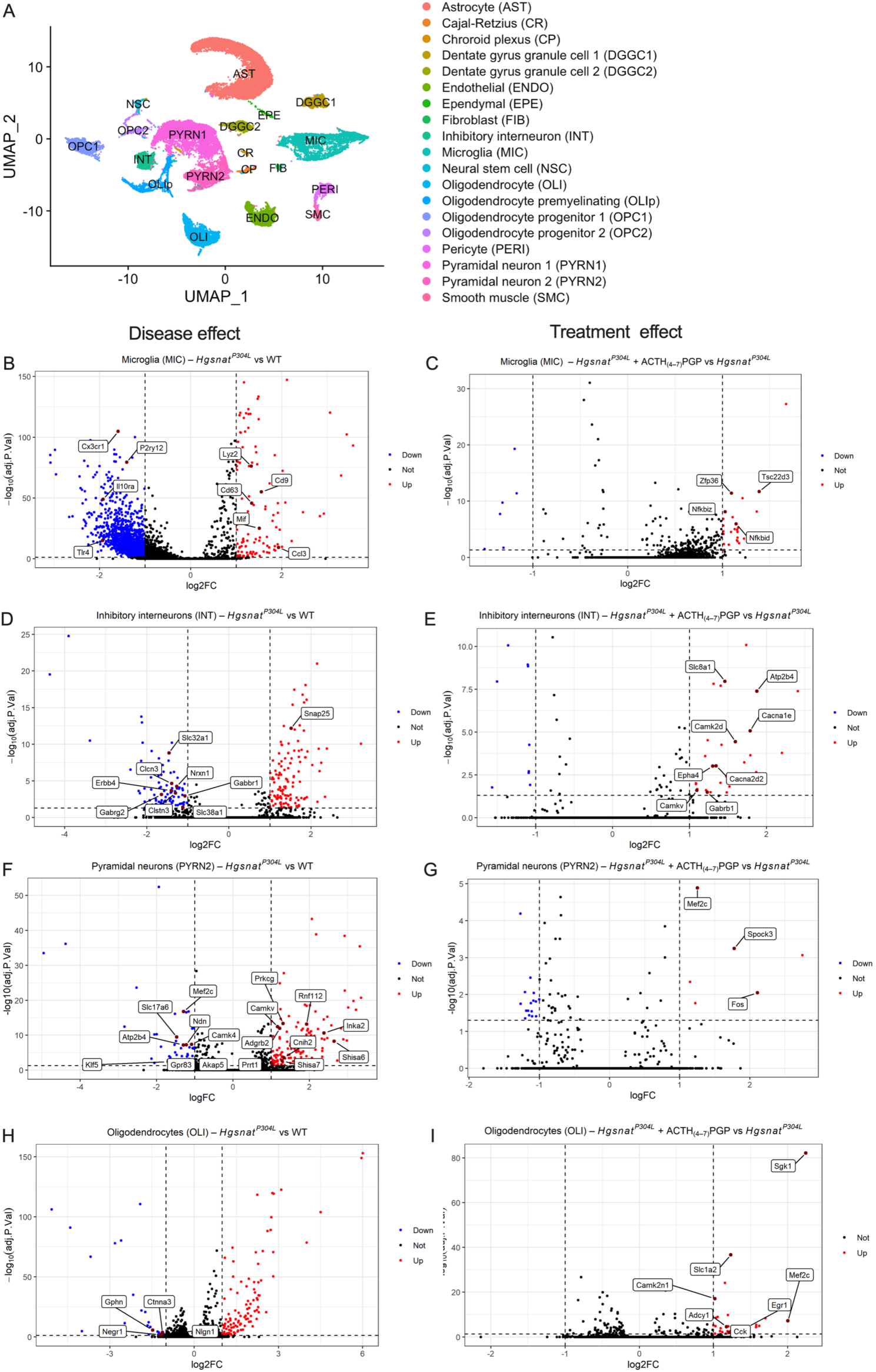
Single-cell RNA sequencing of dissociated hippocampal cells reveals that ACTH_(2-7)_PGP treatment recovers gene expression patterns altered by the disease. **(A)** Uniform manifold approximation and projection (UMAP) plot of single-cell RNA sequencing (scRNA-seq) of hippocampal cells from WT and *Hgsnat^P304L^* mice treated with saline or ACTH_(4-7)_PGP between P27 and P42. **(B-G)** Volcano plots of differentially expressed genes in the microglia (**B,C**), inhibitory interneurons (**D,E**), glutamatergic pyramidal (**F,G**) and oligodendrocytes (**H,I**) for saline-treated WT vs *Hgsnat^P304L^* mice (left) or saline-treated vs ACTH_(4-7)_PGP-treated *Hgsnat^P304L^* mice (right). Threshold of discovery (dotted lines), |log2FC| >1, padj value<0.001 were calculated by Wilcoxon Rank Sum test.

The analysis of the differentially expressed genes (DEG) in response to disease (untreated *Hgsnat^P304L^* mice vs untreated WT) or to the drug (untreated vs treated *Hgsnat^P304L^* mice) was further conducted for each cluster to identify genes with significantly changed expression. The highest number of DEG in response to both disease and the drug were found in the four clusters: microglia (MIC), inhibitory interneurons (INT), glutamatergic pyramidal neurons (PYR1/2), and oligodendrocytes (OLI) (**Figure 10B-I**). In the microglia (MIC) cluster, a comparison of untreated *Hgsnat^P304L^* vs untreated WT mice (**Figure 10B, Figure S16**) revealed a drastic decrease in the expression of genes in the pathways related to differentiation of immune cells, as well as both pro-inflammatory and anti-inflammatory response. This included the genes encoding the TLR4 receptor responsible for recognition of pathogen-associated molecular patterns, PAMPs (*Tlr4*), a subunit of an inflammatory cytokine receptor IL10 (*Il10ra*), and an adenylate cyclase-inhibiting G protein receptor (*P2ry12*), that regulates migration of parenchymal safeguarding microglia [41] (**Figure 10B**). The upregulated pathways also involved multiple genes related to the innate immune response, microglial activation and inflammatory response. These genes included *Cd9* and *Cd63,* encoding components of the CD9/CD63 integrin receptor complex, that regulates integrin-dependent migration of macrophages, particularly relevant for the inflammatory response. Notably, CD63 is also a marker of lysosomal storage in MPS[42]. Other overexpressed genes encoded established MPS IIIC biomarkers, the inflammatory monokine, Mip1α (*Ccl3*) and antibacterial Lysozyme C-2 (*Lyz2*), overexpressed and stored by neuronal cells in MPS IIIB and MPS IIIC mouse models [6,43]. We also detected a decreased expression of the CX3C chemokine receptor 1 (*Cx3cr1*) gene which was previously associated with a reduction of ramified microglia and a rise of neuroinflammatory markers, and Macrophage migration inhibitory factor (*Mif*) pro-inflammatory cytokine involved in the innate immune response. In contrast, ACTH_(4-7)_PGP treatment induced changes in the gene expression compatible with reduction of the pro-inflammatory phenotype of the microglial cells. This involved increased expression of several genes associated with inhibition of NF-κB activity such as *Nfkbiz*, *Nfkbid*, and *Tsc22d3*, which products directly inhibit the NF-κB nuclear translocation, and *Zfp36* that suppresses TNF-α production (**Figure 10C**). In addition, several genes involved in the NF-κB pathway inhibition such as *Nfkbia*, *Rassf2, Nr4a1*, and *Trim30a* showed a significant reduction although were below the |log2FC|>1 cut-off (not shown).

In the inhibitory interneurons (INT, *Slc32a*+ inhibitory neurons), the major pathways downregulated in MPS IIIC compared to WT mice, were related to synaptic organization, structure and activity, GABAergic synaptic transmission, cognition, learning and memory, as well as the pathways related to vesicular transport (**Figure S16**). These changes were, in general, consistent with the alterations we previously detected at the protein level [8]. We also detected alterations in the expression of the genes related to the ER stress and ERAD pathway, which we previously reported in *Hgsnat^P304^* mice [33]. At the individual gene level, we found a major reduction in the expression of the genes encoding GABA receptor subunits (*Gabbr1* and *Gabrg2*), receptor tyrosine-protein kinase ERBB-4, an essential cell surface receptor for neuregulin (*Erbb4*), postsynaptic adhesion molecule Calsyntenin-3 (*Clstn3*), neuroligin-binding protein Neurexin-1 (*Nrxn1*) and multiple interneuron specific transporters including a glutamate transporter *Slc38a1*, H^+^/Cl^-^ exchange transporter 3 (*Clcn3*), and Vesicular inhibitory amino acid transporter VGAT (*Slc32a1*) whose deficiency we have also detected at the protein level. The expression of the *Snap25* gene encoding the synaptosomal-associated protein 25, a biomarker of synaptic loss in neurodegeneration [44], was upregulated (**Figure 10D**).

In interneurons of treated mice, we observed an induction of multiple synaptic genes including those encoding subunits of Voltage-dependent P/Q-type and N-type calcium channels (*Cacna2d2* and *Cacna1e*, respectively) pivotal to neuronal function, Sodium/calcium exchanger 1 (*Slc8a1*), an essential component of post-synaptic densities that regulates voltage-gated channels activity, GABA receptor subunit (*Gabrb1*), and Plasma membrane calcium-transporting ATPase 4 (*Atp2b4*). We also detected increased expression of the Calcium/calmodulin-dependent protein kinase type II (*Camk2d*), CaM kinase-like vesicle-associated protein (*Camkv*), and the Ephrin type-A receptor 4 (*Epha4*) genes associated with regulation of synaptic plasticity and axonal guidance (**Figure 10E**).

In the glutamatergic pyramidal neuron cluster (PYRN2, the marker VGLUT1 glutamate transporter gene, *Slc17a7*), UMAP visualization revealed several distinct subclusters with condition-specific enrichment (**Figure S15B**). Subclusters 0, 1, and 6 were predominantly composed of cells from WT mice, while subclusters 2, 3, and 7 were enriched with cells from the *Hgsnat^P304L^* group. Based on this distribution, we merged subclusters 0, 1, and 6 into the control-associated group, and subclusters 2, 3, and 7 into the disease-associated group to identify disease-related transcriptional alterations. We, then, used the disease-associated subclusters 2, 3, and 7 to evaluate the transcriptional response to the treatment in the *Hgsnat^P304L^* mice. The analysis of transcriptional changes altered in response to the disease pointed to a profound dysregulation of synaptic and neuronal homeostasis in glutamatergic neurons in *Hgsnat^P304^* mice (**Figure S16**). The downregulation of genes encoding vesicular glutamate transporter, VGLUT2 (*Slc17a6*), a transcription activator Myocyte-specific enhancer factor 2C (*Mef2c*) that regulates formation of the excitatory synapse, the scaffold protein A-kinase anchor protein 5 (*Akap5*) that regulates plasticity at excitatory synapses as well as other key genes such as *Klf5*, *Gpr83*, *Ndn*, *Camk4*, and *Atp2b4* was consistent with a reduced glutamate release, impaired activity-dependent gene regulation, and disruption of calcium signaling pathways, all of which are essential for excitatory transmission and neuronal plasticity (**Figure 10F**).

Upregulated genes also included those involved in AMPA/NMDA receptors biogenesis, assembly trafficking and stabilization (*Shisa6*, *Shisa7*, *Cnih2*, *Prrt1*, *Prkcg*, *Camkv*) as well as other essential genes of neuronal differentiation, dendritic structure, and plasticity-related signaling cascades (*Inka2*, *Rnf112*, and *Adgrb2*) (**Figure 10F**). These changes, most probably, point to a compensatory remodeling of synaptic architecture or/and a maladaptive response leading to circuit imbalance consistent with the results of our electrophysiological experiments (**Figure 3**).

In contrast, comparison of gene expression in treated vs untreated MPS IIIC mice revealed a coordinated upregulation of genes associated with neuronal activity (*Fos*, *Mef2c*), and extracellular remodeling (*Spock3*), suggesting that treatment triggers a partial restoration of functional synaptic plasticity, and may support neuroprotection or circuit reorganization in the disease state (**Figure 10G**).

The analysis of oligodendrocyte and related cell type clusters (OLI, OLIp, OPC1, OPC2) in untreated *Hgsnat^P304L^* vs WT mice revealed a reduction in the relative number of mature oligodendrocytes (OLI) and a corresponding increase in the relative number of oligodendrocyte progenitors OPC1 and OPC2 (OPC1+OPC2 / OLI cell count ratio = 0.865 in the WT and 1.01 in the MPS IIIC brains). This scarcity of mature oligodendrocytes, also detected by immunofluorescence analysis in *Hgsnat^P304L^* mouse brain [13], can potentially be responsible for myelination defects in the mouse models and patients. In the brains of ACTH_(4-7)_PGP-treated mice the OPC1+OPC2 / OLI cell count ratio was 0.821, similar to the WT mice. When gene expression profiles for the OLI cluster were compared for WT and MPS IIIC brains, we observed a reduction in the expression of genes participating in the positive regulation of cell projection organization. This included the *Nlgn1* gene encoding for Neuroligin-1, essential protein for synaptic formation, the *Negr1* gene encoding the neuronal growth regulator 1, a mediator of permissive axon-myelin interactions, the *Gphn* gene encoding for inhibitory postsynaptic scaffold protein Gephyrin, present in the oligodendrocyte precursor cells where it contributes to myelin sheath formation by forming neuron-glial synapse [45] and the *Ctnna3* gene encoding alpha-T-catenin abundantly detected in oligodendrocytes [46] **(Figure 10H)**. Notably, catenin proteins participate in the Wnt/β-catenin signaling pathway essential for the initial myelination (reviewed in the ref. [47]). In contrast, differential gene expression (DGE) and Gene Ontology (GO) enrichment analyses revealed that, in *Hgsnat^P304L^* mice, the drug treatment resulted in the upregulation of genes associated with key neurobiological processes, including cognition (GO:0050890), learning and memory (GO:0007611), regulation of synaptic plasticity (GO:0048167), long-term potentiation (long-term synaptic potentiation, GO:0060291), and neuronal migration (neuron migration, GO:0001764). Representative genes include *Sgk1, Camk2n1, Mef2c, Adcy1, Egr1, Cck, and Slc1a2* (**Figure 10I**; **Figure S16**).

Overall, our scRNA-seq results are consistent with the suggestion that the drug mitigates the changes in expression of brain cell genes related to neuroinflammatory response and defects in neuronal synaptogenesis and myelination.

### 7. Therapeutic action of ACTH_(4-7)_PGP is mediated through melanocortin receptor 4 (MC4R) and BDNF-cAMP response element-binding protein (CREB) pathway

We hypothesized that ACTH_(4-7)_PGP, as a melanocortin derivative, induces synaptogenesis and reduces neuroinflammation by acting on the melanocortin receptors 3 and/or 4 (MC3/4R) expressed in the brain cells. To test this, cultured primary WT mouse cortical neurons or iPSC-derived human cortical MPS IIIB neurons were treated with either 10 µM ACTH_(4-7)_PGP or a 10 µM peptide with a scrambled sequence (Ac-PEMHGFP-OH, SCR) in the absence or presence of broad-spectrum melanocortin antagonist (SHU9119, acting concentration 10 µM) or a selective MC4R antagonist (HS024, active concentration 0.5 µM). By immunofluorescence analysis, we found that BDNF levels increased in neurons treated with ACTH_(4-7)_PGP but not with a scrambled peptide. In contrast, cells treated with ACTH_(4-7)_PGP in the presence of either SHU9119 or HS024 did not show an increase in BDNF levels suggesting that both MC3R/MC4R and MC4R antagonists blocked the effect of the drug (**Figure 11A**). These results were confirmed by immunoblotting in SH-SY5Y human neuroblastoma cell line differentiated into neurone-like cells by a 4-day induction with retinoic acid in a serum-free media [48]. The cells were further cultured in a complete neurobasal media for 3 days, before exposure to ACTH_(4-7)_PGP for 24 h in the absence or presence of SHU9119 and HS024 (**Figure 11B**). Cells treated with ACTH_(4-7)_PGP alone, but not those co-treated with MC3/4R antagonists, revealed increase both in BDNF levels and the levels of phosphorylated TrkB (Tropomyosin receptor kinase B), the BDNF receptor essential for synaptic gene expression, reviewed in [49,50]. We further analyzed phosphorylation of the BDNF transcription factor, cAMP response element-binding protein (CREB) as well as Extracellular signal-regulated kinase 1/2 (ERK 1/2), and protein kinase A (PKA/AKT), the kinases common for major signalling pathways downstream of MC3R and MC4R [51,52]. ACTH_(4-7)_PGP induced a rapid and long-term phosphorylation of CREB with the maximal pCREB levels observed between 30 and 90 min of treatment. We also detected rapid phosphorylation of the PI3K substrate kinase, AKT (maximal level at 15-90 min), and a steady (15-120 min) increase in the levels of pTrkB (**Figure S17**). Conversely, SH-SY5Y cells showed a high basal level of ERK1/2 phosphorylation, consistent with previous reports [53] and not altered by the drug. These results were confirmed in iPSC-derived neurons of a healthy human subject. In these cells, in addition to phosphorylation of AKT, CREB, and TrkB, ACTH_(4-7)_PGP also induced levels of pERK1/2, suggesting that the drug action involves an interplay between ERK and PI3K/AKT signaling (**Figure S18**). Based on these results, we further tested whether the phosphorylation of PI3K, AKT and CREB induced by ACTH_(4-7)_PGP in SH-SY5Y cells was blocked by pre-treatment with SHU9119, or HS024. The results of these experiments confirmed that both antagonists blocked activation of the AKT/CREB/TrkB pathway suggesting that it was mediated by MC4R receptors (**Figure 11C**).

**Figure 11.**
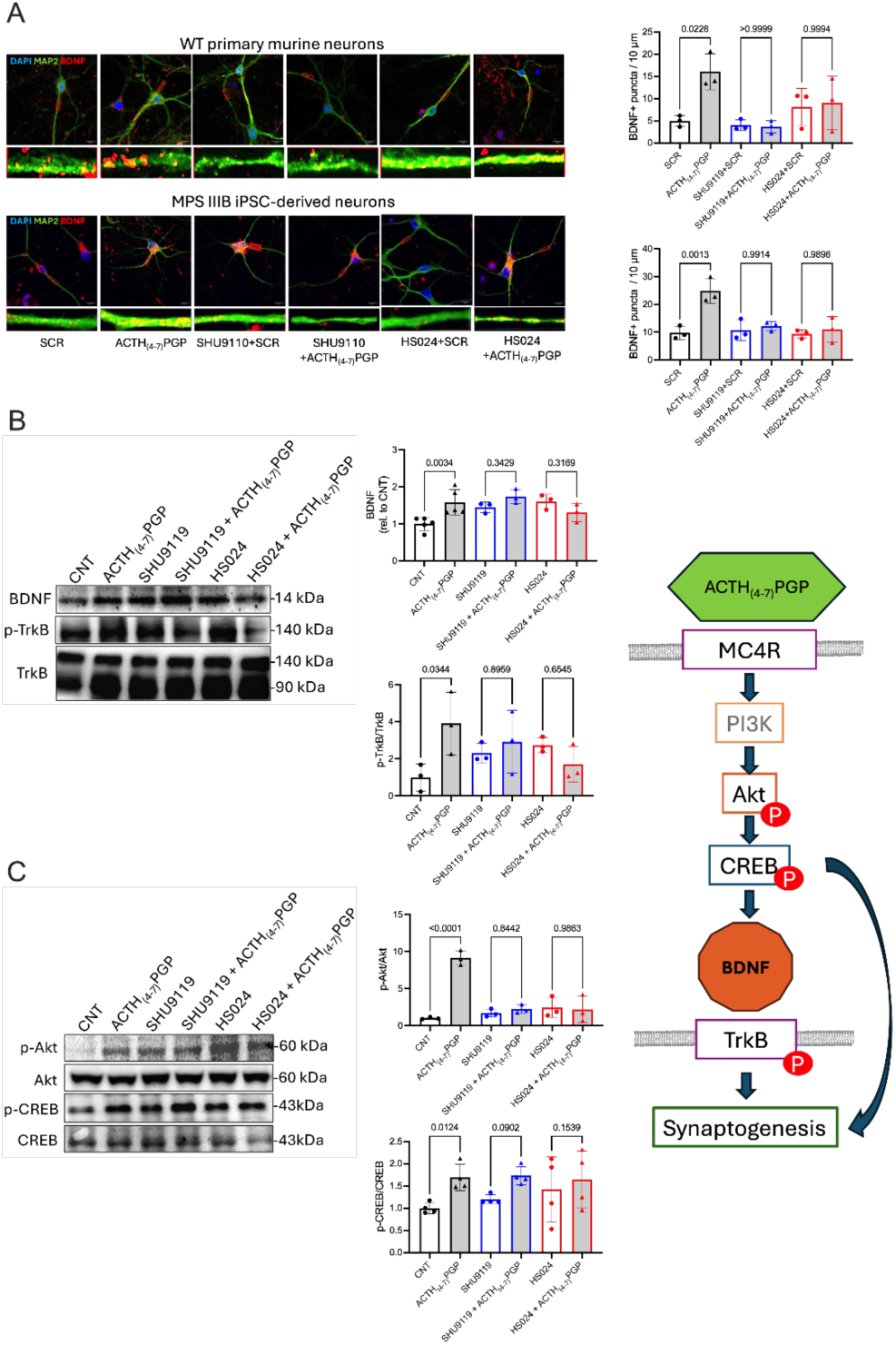
ACTH_(4-7)_PGP acts on neuronal MC3/4R receptors and activates the PI3K/AKT/CREB signaling pathway. **(A)** Representative images of cultured primary WT mouse and iPSC-derived human MPS IIIB cortical neurons treated in culture with ACTH_(4-7)_PGP and scrambled (SCR) peptides in the absence or presence of MC3/4R antagonists, SHU9119 or HS024 labeled for MAP2 (green) or BDNF (red). Nuclei were labeled with DAPI (blue). The scale bars equal 10 µm. Inserts show enlarged images of neurites selected at ≥10 µm from the soma (red rectangles). The graphs show quantification of BDNF+ puncta along 10-µm segments of neuronal projections by ImageJ software. Individual values, means and SD from 3 biological replicates (>15 cells in each experiment) are shown. *P*-values were calculated using one-way ANOVA and Tukey post hoc test. **(B-C)** Representative immunoblots showing the abundance of BDNF, p-TrkB and TrkB **(B)**, or Akt, p-AKT, CREB, and p-CREB **(C)** in differentiated SH-SY5Y neuroblastoma cells treated with ACTH_(4-7)_PGP for 90 min, with or without pre-incubation with MC3/4R antagonists, SHU9119 or HS024. Bar graphs show quantification of BDNF band intensities normalized to the total protein or ratios of p-Akt/Akt and p-CREB/CREB intensities relative to untreated control cells (CNT). Individual data (n = 3-5) and means ± SD are shown. Statistical analysis was performed using one-way ANOVA followed by Šídák’s post hoc test.

To get insight into the anti-inflammatory action of ACTH_(4-7)_PGP, we analysed whether the MC3/4R antagonists block the effect of the drug in primary mouse microglia cells. Microglia from brains of neonatal (P1-P2) WT and *Hgsnat^P304L^* mice were treated with ACTH_(4-7)_PGP or the scrambled peptide in the absence or presence of MC3/4R antagonists for 24 h followed by the analysis of the intracellular BDNF levels by immunofluorescence microscopy. ACTH_(4-7)_PGP (but not the scrambled peptide) drastically increased BDNF levels in *Hgsnat^P304L^*microglia (**Figure 12A**). Like in neurons, this effect was blocked by the MC3/4R antagonists suggesting that ACTH_(4-7)_PGP induced production of BDNF through MC4R in both neurons and microglia (**Figure 12A**). When activation of untreated or ACTH_(4-7)_PGP-treated *Hgsnat^P304L^* microglia was studied by immunofluorescence analysis, we found that, compared to WT cells, *Hgsnat^P304L^* microglia had an increased size, showed higher levels of Iba1 protein and higher affinity to ILB4 lectin characteristic of the proinflammatory M1 phenotype. A 24 h treatment with 10 µM ACTH_(4-7)_PGP (but not with the scrambled peptide), reduced Iba1 and ILB4 labeling to the WT levels (**Figure 12B**). Pre-treatment of cells with either SHU9119, or HS024, completely abolished the anti-inflammatory effect of ACTH_(4-7)_PGP confirming that MCR4 is the primary drug target also in microglia (**Figure 12B**).

**Figure 12.**
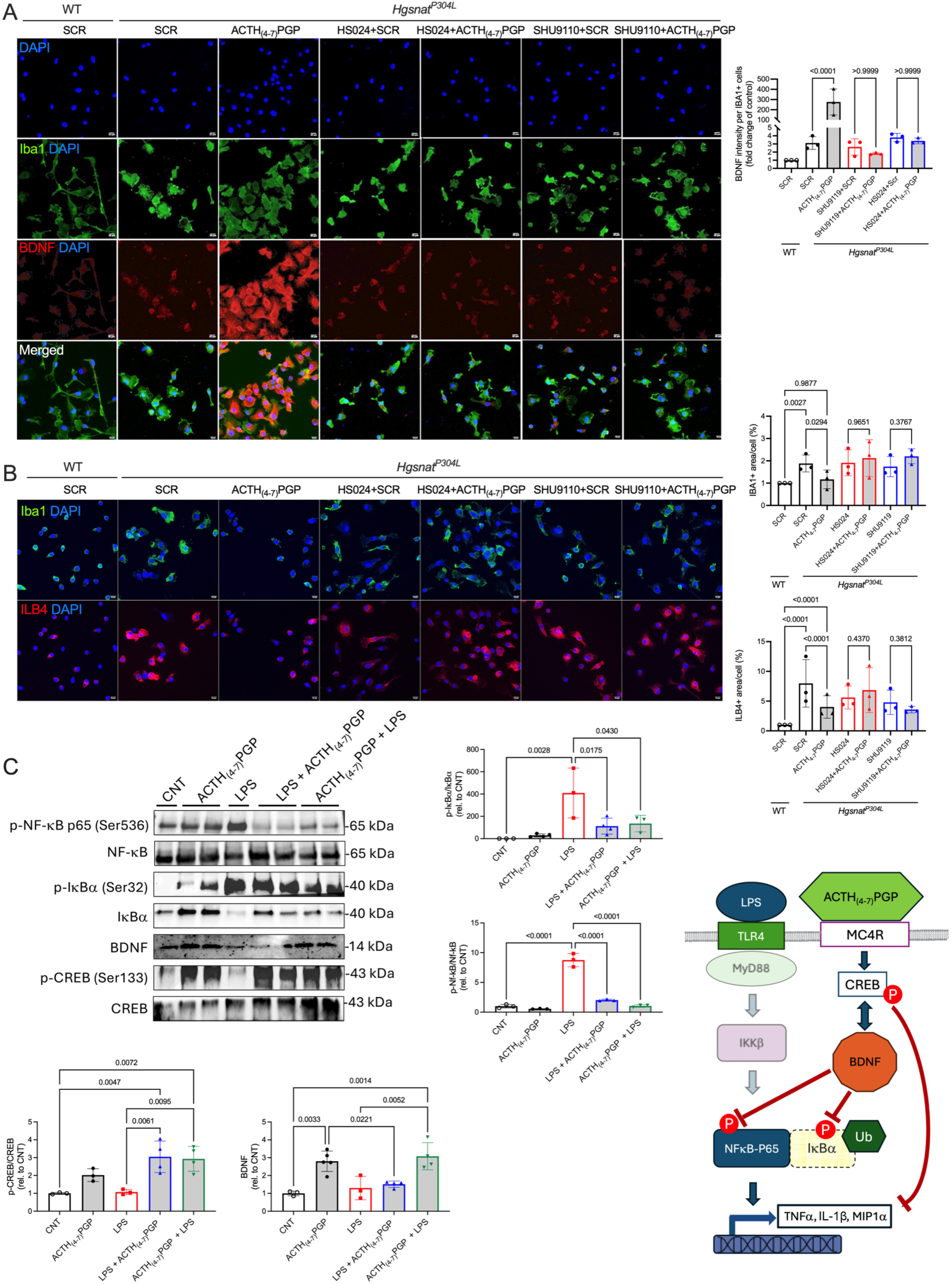
ACTH_(4-7)_PGP acts on microglia MC4R receptors, activates the PI3K/AKT/CREB signaling pathway and blocks the NF-κB phosphorylation and activity. **(A)** ACTH_(4-7)_PGP increases BDNF level in cultured mouse microglia *via* the MC4R receptors. Representative ICC images of primary WT and *Hgsnat^P304L^* murine microglia cells labeled with antibodies against BDNF (red) and the microglia marker Iba1 (green). DAPI (blue) was used as a nuclear counterstain. The cells were treated for 24 h with 10 µM ACTH_(4-7)_PGP or scrambled peptide (SCR) in the presence or absence of 10 µM SHU9119 or 0.5 µM HS024. Graph shows quantification of BDNF in Iba1+ cells using ImageJ software. Statistical significance of changes was estimated using one-way ANOVA with Tukey post hoc test. **(B)** ACTH_(4-7)_PGP attenuates inflammatory phenotype of cultured mouse *Hgsnat^P304L^* microglia *via* the MC4R receptors. Representative ICC images of primary WT and *Hgsnat^P304L^* murine microglia cells labeled with antibodies against Iba1 (green). DAPI (blue) was used as a nuclear counterstain. The cells were treated for 24 h with 10 µM ACTH_(4-7)_PGP or scrambled peptide (SCR) in the presence or absence of 10 µM SHU9119 or 0.5 µM HS024. Graph shows quantification of BDNF in Iba1+ cells using ImageJ software. Statistical significance of changes was estimated using one-way ANOVA with Tukey post hoc test. **(C)** Representative immunoblots showing the abundance of p-NF-κB, NF-κB, p-IκBα, IκBα, BDNF, p-CREB and CREB in untreated primary WT mouse microglia cells (CNT), or cells treated with ACTH_(4-7)_PGP for 60 min, 10 nM LPS for 60 min, LPS for 48 h followed by ACTH_(4-7)_PGP for 30 min, or ACTH_(4-7)_PGP for 48 h followed by LPS for 60 min. Bar graphs show quantification of BDNF band intensities normalized to the total protein or ratios of p-NF-κB/NF-κB, p-IκBα/IκBα and p-CREB/CREB intensities relative to untreated control cells (CNT). Individual data (n = 3-5) and means ± SD are shown. Statistical analysis was performed using one-way ANOVA followed by Šídák’s post hoc test.

Since previous work established that HS stimulates activation of microglia by acting on TLR receptors and triggering the NF-κB pathway [5], we further tested whether ACTH_(4-7)_PGP inhibits the NF-κB activity in microglia. We, first, established that treatment of cultured primary WT mouse microglia with 10 nM LPS for either 1 h or 48 h resulted in a drastic increase in phosphorylation of the 65-kDa NF-κB peptide (NF-κB p65) analysed by immunoblot (not shown). We further assessed if LPS-induced p65 phosphorylation was altered by 10 µM ACTH_(4-7)_PGP. In the first experimental setting designed to model the ability of the drug to protect cells from inflammatory activation, the cells were pre-treated with ACTH_(4-7)_PGP for 48 h before being exposed to LPS for 1 h. In the second experimental setting designed to study the acute ACTH_(4-7)_PGP effect on preactivated microglia, the drug was added for 30 min to the cells pre-treated with 10 nM LPS for 48 h. These experiments showed that ACTH_(4-7)_PGP, added to the cells either 48 h before or 1 h after LPS, reduced the phosphorylation level of NF-κB p65 peptide compared to LPS-treated cells (**Figure 12C**).

Since our scRNA-seq analysis revealed that a 15-day treatment with ACTH_(4-7)_PGP increases expression of IκBα in hippocampal microglia *in vivo*, we analyzed levels of this protein in the homogenates of cultured microglia treated with LPS or with a combination of ACTH_(4-7)_PGP and LPS as described above. We found that the levels of total IκBα protein were increased in the ACTH_(4-7)_PGP-treated cells, while the IκBα phosphorylation at the Ser32 residue (which precedes its sequestering to the proteosomes for degradation) was decreased (**Figure 12C**). This was consistent with the suggestion that in ACTH_(4-7)_PGP-treated cells majority of NF-κB protein was in the cytoplasmic IκBα-bound inactive form. ACTH_(4-7)_PGP-treated microglia also had substantially increased levels of p-CREB (**Figure 12C**), known to directly inhibit the transcriptional activity of NF-κB by competing with it for the association with the nuclear coactivator CBP [54,55].

## Discussion

Despite many years of extensive research, no specific treatments have been approved for Sanfilippo patients. Preclinical work and clinical trials for *in vivo* and *ex vivo* gene therapy are underway, however, as we currently know, gene therapy alone cannot reverse CNS pathology in the symptomatic patients emphasizing the need for development of additional/alternative strategies including those using small molecule drugs [56,57]. In the current work, we examined whether synaptic deficits, the hallmarks of the CNS pathology in the diseases of the Sanfilippo spectrum can be reversed or prevented by ACTH_(4-7)_PGP, a synthetic peptide analog of ACTH hormone known to induce BDNF in several experimental models.

Through its action on TrkB, BDNF supports neuronal survival, induces neurogenesis and synaptogenesis related to learning and memory [58,59] making it potentially useful for treatment of neurodegenerative disorders, like the Sanfilippo spectrum. However, the direct use of BDNF for therapy is complicated. One of the main problems is to achieve a sustained BDNF concentration in the CNS, which is challenged by short half-life of the recombinant protein and its inability to cross the BBB. To resolve these issues, different research groups either used BDNF fused with cell-penetrating peptides or expressed it in the CNF using AAV vectors [60,61]. In particular, an AAV mediated expression of BDNF in striatal neurons in the R6/2 mouse model of Huntington’s disease induced neurogenesis and increased the lifespan of animals [62]. However, the use of AAV is limited by its narrow biodistribution and a common immunogenicity to the virus. Stem cell transplants overexpressing BDNF and capable to specific migration to inflammation and apoptosis sites, showed efficacy, but their integration was short-term [63–66]. This is why, more and more attention is now given to the use of small molecule compounds capable of inducing brain BDNF levels such as the hybrid synthetic peptide ACTH_(4-7)_PGP.

The results of the current study demonstrate that acetylated at the N-terminus ACTH_(4-7)_PGP can ameliorate pathological signs in the human neuronal models of MPS IIIA, IIIB and IIIC as well as in the mouse models of MPS IIIA and IIIC. Treatment of cultured primary hippocampal neurons of *Hgsnat^P304L^* mice and iPSC-derived cortical neurons of MPS IIIA, MPS IIIB and MPS IIIC patients rescued deficient levels of synaptic markers VGLUT1, SYN1, PSD-95 and BDNF, suggesting that the drug can potentially ameliorate defects in synaptic transmission. This was further tested in MPS IIIA and MPS IIIC mice by measuring miniature and induced synaptic currents on the pyramidal CA1 neurons in acute hippocampal slices. Both MPS IIIA and MPS IIIC mice showed a clear reduction in excitatory and inhibitory miniature events at P60, indicating reduced synaptic transmission and consistent with an overall reduction in the number of functional synapses. Mice also showed drastically reduced AMPA/NMDA ratios in evoked glutamatergic responses consistent with reduced excitatory transmission and enhanced synaptic depression. These results well correlated with our scRNA-seq experiments which identified a dual pattern of synaptic downregulation and compensatory upregulation that may contribute to these synaptic dysfunctions as well as to neurodevelopmental impairments characteristic of the disease and highlighted the need to consider both loss-of-function and gain-of-function transcriptional adaptations in Sanfilippo disease. Notably, in both MPS IIIA and MPS IIIC models, a 30-day intranasal treatment with ACTH_(4-7)_PGP increased the synaptic activity to the levels of control WT mice.

Administration of ACTH_(4-7)_PGP to 4-month-old MPS IIIC mice for 10 consequent days rescued impairment of short-term memory in NOR test, while a single dose given 17 h before the test had no effect, indicating that a prolonged administration of the peptide is required to rescue memory deficits in Sanfilippo animals. Thus, we further tested the clinical efficacy of long-term administration of ACTH_(4-7)_PGP in MPS IIIA (*Sgsh^mps3a^*) and PMS IIIC (*Hgsnat^P304L^*) mice. For both strains, the treatment was started in pre-adolescent/peri-adolescent mice (P30) which corresponds to the neurodevelopmental human age of 10 years [67], the time of diagnosis for majority of Sanfilippo patients (reviewed in [2–4]). Since most patients are diagnosed post-symptomatically, this age would likely be the treatment starting point. Although *Hgsnat^P304L^* and *Sgsh^mps3a^* mice at P30 do not show behavioral alterations, according to our current results and published data [7,8,37], their CA1 pyramidal neurons already reveal synaptic deficits at the electrophysiological level, significantly reduced density of dendritic spines, primary storage of HS in microglia and a high level of micro/astrogliosis. For both models, we observed a rescue of behavioral abnormalities in the cohort of MPS III mice treated with ACTH_(4-7)_PGP. Treated 4-month-old *Hgsnat^P304L^*mice showed a rescue of hyperactivity and reduced anxiety in OF and EPM tests and deficits in spatial and short-term memory in YM and NOR tests. Treated 6-month-old *Sgsh^mps3a^* mice showed a rescue of short-term memory in NOR test, impaired socialization in Crawley’s TCS test and associative learning and long-term memory in CSF test. Notably, for the *Hgsnat^P304L^* mice, treatment also rescued reduced anxiety, hyperactivity and short-term memory deficit at the age of 8 months, the terminal point of the disease in untreated animals.

Necropsy of treated and untreated mice corroborated the results of the behavioural analysis. We found that long-term treatment with ACTH_(4-7)_PGP rescued reduced levels of BDNF and synaptic proteins, SYN1, VGLUT1 and PSD-95 in hippocampal and cortical pyramidal neurons, confirming the results obtained in the neuronal cultures. TEM analysis of *Hgsnat^P304L^* mouse brain revealed that the treatment restored reduced densities of synaptic vesicles in the terminals of excitatory synapses to the levels of WT mice and rescued reduced areas of postsynaptic densities. An increase in the levels of BDNF and synaptic proteins was also observed in the neurons of ACTH_(4-7)_PGP-treated *Sgsh^mps3a^* mice. Somewhat unexpectedly, we observed that the drug also reduced secondary pathological changes, most pronounced in the pyramidal neurons of the somatosensory cortex, including accumulation of GM2 ganglioside, LC3 puncta, SCMAS and Thioflavin-S-positive protein aggregates. It also prevented the loss of myelin, previously detected by immunochemistry and DTI in MPS III patients and MPS IIIC *Hgsnat^P304L^* mouse model [13].

DTI allows the generation of quantitative maps that reveal tissue microstructure by decomposing diffusion signals into axial diffusivity and radial diffusivity. Furthermore, ex-vivo DTI has been shown to be a reliable marker of neuroprotection efficiency [68,69]. While *in vivo* DTI yields more signal, *ex-vivo* imaging allows for high quality maps due to increased acquisition times [70]. In the grey matter, we observed an increase in diffusivity in the three DTI metrics potentially indicative of a tissue loss and increased extracellular space. Therapy was shown to restore diffusivity measures in the grey matter in both models. Diffusivity maps of the white matter tracks in MPS IIIA and MPS IIIC mice showed a significant increase in axial and mean diffusivity in the cerebral peduncle, indicative of edema and tissue loss. Therapy was shown to restore diffusivity metrics mostly in the MPS IIIA model. This approach will likely be useful for clinical translation, since it allows to monitor changes in grey and white matter of study participants.

The drug also mitigated the neuroinflammatory response, decreasing the density of activated microglia and astrocytes and reducing expression levels of pro-inflammatory cytokines, IL-1β and TNFα, and chemotactic chemokine, MIP-1α, in the brains of chronically treated MPS IIIC and MPS IIIA mice. Notably, in *Hgsnat^P304L^* mice, ACTH_(4-7)_PGP delayed the onset of lethal urinary retention by approximately 8 weeks, extending the life of animals from 41 weeks to 49 weeks, and reduced splenomegaly in both *Hgsnat^P304L^* and *Sgsh^mps3a^* mice, suggesting that the drug exerts an anti-inflammatory effect in both CNS and peripheral tissues.

Since the treatment did not change the level of HS storage, we hypothesized that it ameliorates the pathophysiological changes in the brain by normalizing gene expression profiles in the brain cells altered by the disease. To test this, we conducted a scRNA-seq analysis of dissociated hippocampi cells from one-month-old WT and MPS IIIC mice treated with ACTH_(4-7)_PGP or saline for 15 consecutive days. This analysis, to the best of our knowledge conducted for the first time for a Sanfilippo animal model, confirmed that the treatment induced changes in the gene expression opposite to those caused by the HS storage. The drug caused an increase in the expression of genes associated with inhibition of inflammatory activation in the microglia. We also detected increased expression of genes encoding synaptic proteins in pyramidal and interneurons confirming the results obtained by immunohistochemistry. In the cluster of oligodendrocytes, we found an increase in the expression of genes essential for axonal myelination. Besides, the number of immature oligodendrocytes was reduced in the brains of treated mice while the number of mature cells increased.

Our results, in general, agreed with those of bulk RNA-seq analysis of brain subcortex tissues in rats with a transient middle cerebral artery occlusion (an established Cerebral Ischemia-Reperfusion model) treated or not with unacetylated ACTH_(4-7)_PGP [71]. Similarly to our study, his report also demonstrated reduced expression of proinflammatory and increased expression of neurotransmitter genes in the treated animals.

The specific target of ACTH_(4-7)_PGP in the brain has never been identified, although it has been hypothesised that the drug may act on melanocortin receptors (see for example ref. [72]). We report here that, in cultured primary mouse microglia, iPSC-derived human cortical neurons and differentiated SH-SY5Y human neuroblastoma cells, ACTH_(4-7)_PGP induced production of BDNF and phosphorylation of its transcription factor, CREB. Both processes were inhibited by pre-treatment of cells with either a pan-specific MC3/4R antagonist or a specific MC4R antagonist. Both antagonists also blocked the outcomes of ACTH_(4-7)_PGP treatment, an increase in the levels of synaptic proteins in the iPSC-derived neurons and the inhibition of LPS-mediated activation in the primary mouse microglia. We, thus, conclude that in both neurons and microglia the therapeutic action of ACTH_(4-7)_PGP is mediated through MC4R receptors. This is also consistent with a previous studies showing that ACTH_(4-7)_PGP antagonized the action of α-MSH on the melanocortin MC4 receptor, but not on the melanocortin MC3 receptor [73].

We, further, showed that, in microglia, ACTH_(4-7)_PGP induces the levels of the NF-κB inhibitory protein, IκBα, also detected *in vivo* by scRNA-seq analysis of ACTH_(4-7)_PGP-treated MPS IIIC mice. This, together with reduction in NF-κB p65 levels and increase in levels of pCREP, a direct competitor of the NF-κB p65/nuclear coactivator CBP signalling, suggested that the drug inhibits the neuroinflammatory response by blocking the NF-κB pathway in microglia.

In neurons, we found that ACTH_(4-7)_PGP treatment causes rapid activation of two CREB effector kinases, ERK1/2 and AKT, followed by attenuated activation of the BDNF receptor kinase, TrkB. Our results, consistent with previous reports implicating these kinases as the major mediators of MCR signalling leading to activation of BDNF-CREB pathway [51,52], suggest that ACTH_(4-7)_PGP acts through CREB signaling, enhancing neurogenesis, long-term potentiation (LTP) and synaptic plasticity dysregulated by lysosomal HS storage.

One of the limitations of the current study is that we did not test ACTH_(4-7)_PGP efficacy in the mouse models of other Sanfilippo subtypes, MPS IIIB and MPS IIID. These studies are currently in progress in our laboratory. It also remains to be to studied whether a long-term administration of ACTH_(4-7)_PGP rescues other pathological changes in the CNS including mitochondrial disfunction and vesicle trafficking defects [6,8]. Finally, we did not specifically study the drug safety beyond demonstrating the absence of changes in the body weight gain and increased longevity of treated animals.

Together, our data provide a compelling evidence that ACTH_(4-7)_PGP delays neurological manifestations in MPS III by rescuing glutamatergic neurotransmission and synaptogenesis defects. They also demonstrate that the drug delays immunoinflammatory response in CNS and peripheral tissues and increases longevity. In our opinion, this justifies testing drug’s efficacy in clinical settings.

## Author contributions

T.M., P.D., G.V., P.B., E.Z., B.N., D.N.R., M.T., P.P.v.V., E.B., S.K., and I.L. conducted experiments and acquired data; T.M., P.D., G.V., P.B., E.Z., B.N., D.N.R., L.M.F., M.T., P.P.v.V., E.B., S.K., S.S., S.T., P.T., C.M., G.D.C., G.P., J.B., G.A.L. and A.V.P. analysed data; F.A., G.A., G.L., J.B., S.T., P.T., J.W. provided essential resources; A.V.P., T.M., P.T., G.L., J.B. designed the experiments and wrote the manuscript (first draft); T.M., P.D., , J.W. and A.V.P. edited the manuscript. All authors read and approved the final version of the manuscript. The order of the 2 first authors’ names was determined by joint decision of these authors.

## Acknowledgements

We thank Dr. Elke Küster-Schöck and the Plateforme d’Imagerie Microscopique (PIM – CHU Sainte Justine) for the help with confocal microscopy, Irene Londono for the help with mouse brain imaging and Dr. Mila Ashmarina for critically reading the manuscript and helpful advice.

## Materials and Methods

### Availability of data and materials

The data, analytic methods, and study materials will be made available to other researchers for purposes of reproducing the results or replicating the procedure.

### Study approval

Generation and analysis of iPSC-derived cortical neurons of MPS III patients was approved by the Ethical research board of CHU Ste-Justine (approval number 2022-3817). Approval for the use of the animals in experimentation was granted by the Animal Care and Use Committee of the CHU Ste-Justine (approval numbers 2022-3452 and 2022-3453).

### Murine models

The knockout *Hgsnat-Geo* and knock-in, *Hgsnat^P304L^*mouse models of MPS IIIC have been previously described [6,33]. The *Sgsh^mps3a^* strain (spontaneously occurring mouse model of MPS IIIA [39]) was purchased from The Jackson Laboratory (B6.Cg-*Sgsh^mps3a^*/PstJ, Stock No. 003780). Heterozygote breeding pairs were used to maintain the colony and generate the initial homozygous mutants, followed by homozygote breading to generate homozygous MPS IIIA and MPS IIIC mice. The animals were housed in the Animal facilities of CHU Ste-Justine, following the guidelines of the Canadian Council on Animal Care (CCAC). The animals were kept in an enriched environment with a 12:12 h light/dark cycle, fixed temperature, humidity and continuous access to water and a normal chow diet (5% fat, 57% carbohydrate). Equal cohorts of male and female mice were studied separately for each experiment, and statistical methods were used to test whether the progression of the disease, levels of biomarkers or response to therapy were different for male and female animals. Since differences between sexes were not detected, the data for male and female mice were pooled together.

### Pharmacokinetics

MPS IIIC 4-month-old mice were dosed intranasally with 10 μl of 50 mM ACTH_(4-7)_PGP in saline (5 μl /nostril). Ten min, 1 h, 2 h and 24 h after dosing, mice were anesthetised with sodium pentobarbital, and 500 μl of blood collected by cardiac puncture. Mice were then sacrificed by cranial dislodgement and their brain and visceral organs extracted. The brain was dissected into 4 segments (frontal to dorsal). Tissues and blood plasma were homogenized in acetonitrile (1:4, tissue (plasma)/solvent ratio). The extracts were spiked with heavy isotope-labelled (Phe U-^13^C_9_; U-^15^N) ACTH_(4-7)_PGP peptide as an internal standard, and analyzed by targeted LC-MS/MS, using parallel reaction monitoring on Orbitrap Exploris 480 instrument.

### Enzyme activity assays

Mouse tissues or cultured cell pellets were snap-frozen in liquid nitrogen before storage at - 80°C. Fifty mg samples were homogenized in 250 µl of H_2_O using a sonic homogenizer (Artek Systems Corporation). For HGSNAT assays, 5 µl aliquots of the homogenates were combined with 5 µl of McIlvain Buffer (pH 5.5), 5 µl of 3 mM 4-methylumbelliferyl-β-D-glucosaminide (Moscerdam), 5 µl of 5 mM acetyl-coenzyme A and 5 µl of H_2_O. The reaction was incubated for 3 h at 37°C, stopped with 975 µl of 0.4 M glycine buffer (pH 10.4), and fluorescence was measured using a ClarioStar plate reader (BMG Labtech). Blank samples were incubated without the homogenates which were added after the glycine buffer. The SGSH activity was measured as previously described by Karpova et al. [74] using the synthetic fluorogenic substrate, 4-methylumbelliferyl-2-Deoxy-2-sulfamino-α-D-glucopyranoside Na salt (4MU-α-GLcNs) in a 0.1 M Tris buffer, pH 6.5. The activity of NAGLU was measured by combining 10 µl of homogenate with 5 µl of 0.5 M sodium acetate buffer (pH 4.3), with 2 µl of BSA (5 mg/mL), and 10 µl of 2 mM 4-Methylumbelliferyl-N-acetyl-α-D-glucosaminide (Sigma-Aldrich) followed by incubation for 2 hr at 37°C. The reaction was stopped with 0.4 M glycine buffer (pH 10.4) and fluorescence was measured as above. The activity of β-hexosaminidase was measured by combining 2.5 µl of 10x diluted homogenate (∼ 2.5 ng of protein) with 15 µl of 0.1 M sodium acetate buffer (pH 4.2), and 12.5 µl of 3 mM 4-methylumbelliferyl N-acetyl-β-D-glucosaminide (Sigma-Aldrich) followed by incubation for 30 min at 37°C. The reaction was stopped 0.4 M glycine buffer (pH 10.4) and fluorescence was measured as above. The activity of acidic β-galactosidase was measured by adding 12.5 µl of 0.4 M sodium acetate, 0.2 M NaCl (pH 4.2) and 12.5 µl of 1.5 mM 4-methylumbelliferyl β-D-galactoside (Sigma-Aldrich) to 10 µl of 10x diluted homogenate (∼1 ng of protein). After 15-min incubation at 37°C, the reaction was stopped with 0.4 M glycine buffer (pH 10.4) and fluorescence was measured as above.

### Behavioral analysis

Y-maze test was used to measure the spontaneous alternation behavior, spatial working memory and exploratory activity of mice as previously described [75]. The maze consisted of three identical white Plexiglas arms (40 × 10 × 20 cm, 120° apart) under dim lighting conditions. Each mouse was placed at the end of one arm, facing the center, and allowed to explore the maze for 8 min. All experiments were performed at the same time of the day and by the same investigator to avoid circadian and handling bias. Sessions were video-recorded and arm entries were scored by a trained observer, unaware of the mouse genotype or treatment. Successful alternation was defined as consecutive entries into a new arm before returning to the two previously visited arms. Alternation was calculated as: [number of alternations/total number of arm entries − 2] × 100.

Novel object recognition test was used for assessing short-term recognition memory [76,77]. Mice were placed individually in a 44 ×33 × 40 cm (length x width x height) testing chamber with light grey Plexiglas walls for 20 min habituation period and returned to their home cage. The next day, mice were placed in the testing chamber for 10 min with two identical objects (blue plastic towers, 3 x 1.5 x 4.5 cm), returned to the home cages, and 1 h later, placed back into the testing chamber in the presence of one of the original (familiar) objects and one novel object (a red plastic base, 4.5 x 4.5 x 2 cm). After each mouse, the test arena as well as the plastic objects were cleaned with 70% ethanol to avoid olfactory cue bias. The discrimination index (DI) was calculated as the difference between the novel object exploration time and the familiar object exploration time, divided by the total exploration time. A preference for the novel object was defined as a DI significantly higher than 0.25 [78]. Mice who showed a side preference (DI ± 0.20) during familiarization period, a total exploration times lower than 3 seconds, or a 100% preference for either the new or familiar object were excluded from the analysis.

The open-field test was performed as previously described [79]. Briefly, mice were habituated in the experimental room for 30 mins before the commencement of the test. Each mouse was then gently placed in the center of the open-field arena (40 x 40 x 20 cm) and allowed to explore for 20 min. The mouse was removed and transferred to its home cage, and the arena was cleaned with 70% ethanol before the next test. Analysis of the behavioral activity was done using the Smart video tracking software (v3.0, Panlab Harvard Apparatus), and total distance traveled and percent of time spent in the center zone were measured for hyperactivity and anxiety assessment, respectively.

The elevated plus-maze test was performed as described by Amegandjin, *et al.* [79]. Each mouse was placed in the center of the elevated plus maze and allowed to freely explore undisturbed for 10 min. After each testing, the mouse was returned to the home cage and the arena was cleaned with 70% ethanol before the commencement of the next test. Analysis of the behavioral activity (percentage of time spent in the center zone, closed arms, and open arms; as well as the number of open arm entries) was done by the Smart v3.0 software.

Contextual fear conditioning used to analyze the ability of mice to associate an environment (context) with a fear-inducing stimulus was performed as previously described [80]. Mice were transported to an isolated behaviour room 30 min before testing. Conditioning was performed in the 30.5 x 24 x 21 cm chamber with all-white paneling, and a clear Plexiglas door. Mice were habituated in the experimental chamber for 3 min on day 0 (D0) preceding the training session, during which no electric shocks were administered. On the training day (D1), during 5 min, mice were exposed to three electric foot shocks (0.6 mA, 2 s duration) administered at 120 s, 200 s, and 280 s. Mice were tested on D2 and D7, for 5 min each time in the same chamber without receiving any electric shocks. Mouse freezing behavior was analyzed using the automated Freeze Frame 5 software (Harvard Apparatus), which scores the percentage of total freezing time and calculates the mean duration of freezing episodes. The minimal movement time threshold was set to 0.75 s.

Social interaction behavior was studied using Crawley’s sociability and preference for social novelty test as previously described [81]. Briefly mice were allowed to freely explore a rectangular, three-chamber box with each chamber measuring 19 x 45 cm and the dividing walls made from clear Plexiglas, with an open middle section and a free access to each chamber. Side chambers contained two identical, wire cup containers. During the first session, one chamber was empty and another, holding a naïve/unfamiliar mouse. During the second session, one chamber was holding a familiar mouse from the first session and another, holding a naïve/unfamiliar mouse. All chambers were cleaned with 70% ethanol after each trial. Mice were habituated for 5 min, after which the walls between the compartments were removed to allow free access for the subject mouse to explore each of the three chambers for the duration of 10 min. The time spend in each chamber and the time spend interacting with wire containers were calculated based on the video records.

### Transmission electron microscopy

At 3 and 6 months, 3 mice from each group were anesthetized with sodium pentobarbital (50 mg/kg BW) and perfused with PBS, followed by 2.5% glutaraldehyde in 0.2 M phosphate buffer (pH 7.2). The brains were extracted and post-fixed in the same fixative for 24 h at 4 °C. The hippocampi were dissected, mounted on glass slides, stained with toluidine blue and examined on a Leica DMS light microscope to select the CA1 region of the hippocampus for electron microscopy. The blocks were further embedded in Epon, and 100 nm ultrathin sections were cut with an Ultracut E ultramicrotome, mounted on 200-mesh copper grids, stained with uranyl acetate (Electron Microscopy Sciences) and lead citrate, and examined on a FEI Tecnai 12 transmission electron microscope. For quantification, the micrographs were taken with 13,000 x and 30,000 x magnification.

### Mouse primary cell cultures

Primary hippocampal neurons were cultured from the brains of embryo at gestational day 16 (E16). The hippocampi were isolated and treated with 2.5% trypsin solution (Sigma-Aldrich, T4674) for 15 min at 37°C. The cells were washed 3 times with Hank’s Balanced Salt Solution (HBSS, Gibco, 14025-092) and mechanically dissociated by pipetting, using glass Pasteur pipettes with 3 different opening sizes (3, 2 and 1 mm). Then, they were counted with the viability dye trypan blue (ThermoFisher Scientific, 15250061), using a hemocytometer, and resuspended in Neurobasal media (Gibco, 21103-049) containing L-glutamine, B27, N2, penicillin and streptomycin. The cells were plated at a density of 60,000 cells per well, respectively, in a 12-well plate on coverslips previously coated with Poly-L-Lysine (Sigma Aldrich, P9155). Cells were cultured for 21 days in a humidified incubator at 37°C with 5% CO₂, and 50% of media was changed every three days. Primary cortical neuron cultures were established from tissues of postnatal WT or MPS IIIC pups (P0–P3) using the same technique. Primary glial cell cultures were established from the brains of WT or MPS IIIC pups (P0–P3) as described [82]. Briefly, brain hemispheres were treated with a 2.5% trypsin solution, mechanically dissociated as described for the neuronal cultures, resuspended in DMEM (Gibco, 21103-049) supplemented with 10% fetal bovine serum (FBS), penicillin, and streptomycin and plated in poly-D-lysine (PDL)-precoated T75 cell culture flasks. After 14 days in culture (D14), microglial cells were isolated by shaking the flasks on a rotary shaker at 180 rpm for 3 h. The medium containing the floating cells was then collected, centrifuged, and resuspended in DMEM supplemented with 10% FBS. Microglia cells were plated at a density of 50,000 cells per well in 24-well plates containing coverslips pre-coated with Poly-D-Lysine (Sigma-Aldrich, P9155). The cells were further cultured for 21 days, with 50% of media changed every three days and treated with 10 nM LPS (Lipopolysaccharides from *Escherichia coli* O111:B4, Sigma, L4391) and/or ACTH_(4-7)_PGP as described in the Results section.

### SH-SY5Y cell culture

Human neuroblastoma SH-SY5Y cells (ATCC® CRL-2266™, Manassas, VA, USA) were seeded in T25 cell culture flasks at a density of 600,000 cells/flask and grown in DMEM/F-12 medium supplemented with 10% FBS, 100 U/mL penicillin, and 100 µg/mL streptomycin. Twenty-four h after seeding, the medium was replaced with FBS free DMEM/F-12, supplemented with 1 µM retinoic acid (RA), to induce neuronal differentiation as described [48]. Cells were maintained in the differentiation medium for 5 days, with 50% of media changed at D3. At D5, the medium was switched to Neurobasal medium supplemented with L-glutamine, N2, and B27 and the cells were cultured for an additional 5 days in the absence or presence of AVP6 and MC3/4R antagonists.

### iPSC-derived neuronal cultures

*Generation of iPSC lines* MPS III patient fibroblast lines were obtained from the Coriell Institute for Medical Research (NJ, USA), or from the hospitals were the patients were diagnosed/followed with the informed consent of patient’s families (see Table 1 for details). The fibroblasts were propagated in Dulbecco’s Modified Eagle Medium (DMEM, ThermoFisher) with 10% fetal bovine serum (FBS) and 1% Antibiotic-Antimycotic (15240062, ThermoFisher) and tested for mycoplasma. The cells were further reprogrammed into iPSCs at the CHUSJ iPSC Platform using a non-integrating CytoTune-Sendai viral reprograming kit (A16517, Thermo Fisher Scientific, MA, USA) according to the manufacture’s protocol. Two colonies for each iPSC line were used for further proliferation. iPSCs were expanded and maintained on six-well plates coated with Matrigel^®^ mTeSR™ Plus medium at 37◦C, in 5% CO_2_/5% O_2_ atmosphere following the medium manufacturer’s protocol [83]. At 60-80% confluency the cells were passaged using Gentile Cell Dissociation reagent (GCDR, StemCell Technologies^TM^) and plated in mTeSR™ Plus medium containing 10 µM RI (Y27632 ROCK inhibitor, Selleckchem). The following day, the medium was replaced by fresh mTeSR™ Plus medium without RI.

**Table 1:**
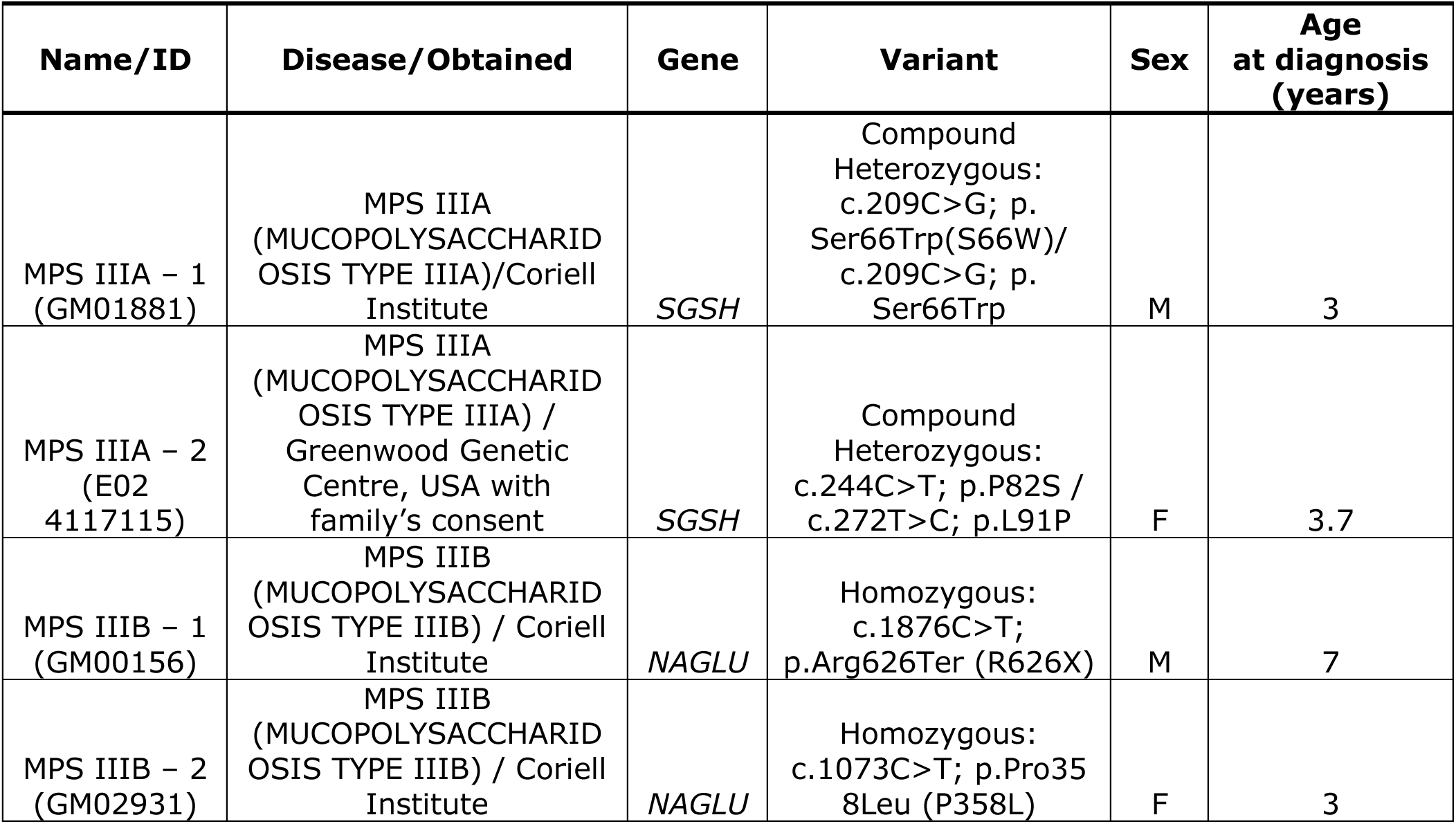
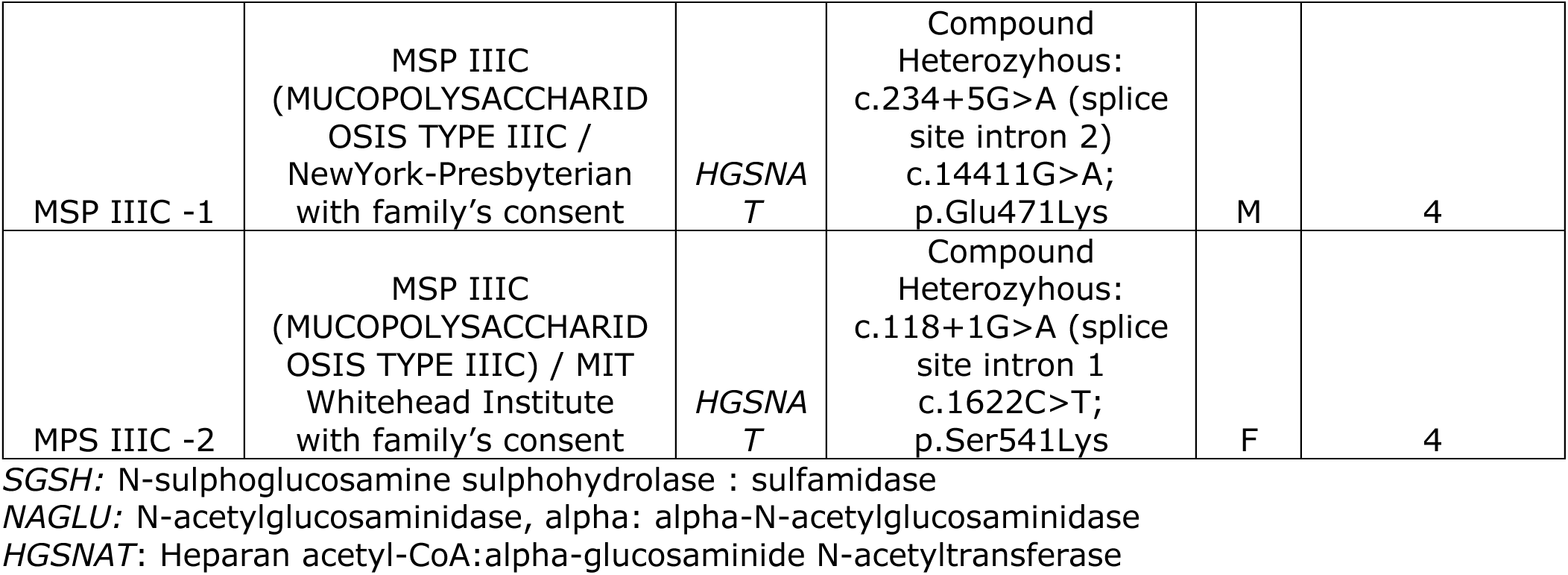
Fibroblast cell lines of MPS III patients used for generation of iPSCs.

| Name/ID | Disease/Obtained | Gene | Variant | Sex | Age at diagnosis (years) |
| --- | --- | --- | --- | --- | --- |
| MPS IIIA – 1 (GM01881) | MPS IIIA (MUCOPOLYSACCHARID OSIS TYPE IIIA)/Coriell Institute | *SGSH* | Compound Heterozygous: c.209C>G; p. Ser66Trp(S66W)/ c.209C>G; p. Ser66Trp | M | 3 |
| MPS IIIA – 2 (E02 4117115) | MPS IIIA (MUCOPOLYSACCHARID OSIS TYPE IIIA) / Greenwood Genetic Centre, USA with family's consent | *SGSH* | Compound Heterozygous: c.244C>T; p.P82S / c.272T>C; p.L91P | F | 3.7 |
| MPS IIIB – 1 (GM00156) | MPS IIIB (MUCOPOLYSACCHARID OSIS TYPE IIIB) / Coriell Institute | *NAGLU* | Homozygous: c.1876C>T; p.Arg626Ter (R626X) | M | 7 |
| MPS IIIB – 2 (GM02931) | MPS IIIB (MUCOPOLYSACCHARID OSIS TYPE IIIB) / Coriell Institute | *NAGLU* | Homozygous: c.1073C>T; p.Pro358Leu (P358L) | F | 3 |
| MSP IIIC -1 | MSP IIIC<br>(MUCOPOLYSACCHARID<br>OSIS TYPE IIIC /<br>NewYork-Presbyterian<br>with family's consent | HGSNA<br>T | Compound<br>Heterozyhous:<br>c.234+5G>A (splice<br>site intron 2)<br>c.14411G>A;<br>p.Glu471Lys | M | 4 |
| MPS IIIC -2 | MSP IIIC<br>(MUCOPOLYSACCHARID<br>OSIS TYPE IIIC) / MIT<br>Whitehead Institute<br>with family's consent | HGSNA<br>T | Compound<br>Heterozyhous:<br>c.118+1G>A (splice<br>site intron 1)<br>c.1622C>T;<br>p.Ser541Lys | F | 4 |

*SGSH*: N-sulphoglucosamine sulphohydrolase : sulfamidase
*NAGLU*: N-acetylglucosaminidase, alpha: alpha-N-acetylglucosaminidase
*HGSNAT*: Heparan acetyl-CoA:alpha-glucosaminide N-acetyltransferase

*Induction of cortical NPC and cortical neurons* iPSCs were differentiated into cortical forebrain committed neural precursor cells (NPCs) by dual SMAD inhibition, as described [35] by passaging iPSCs to Matrigel^®^ coated dishes with subsequent passaging to poly-L-ornithine (PO)/laminin coated dishes. NPC induction was performed in a monolayer with the cortical neuronal induction media essentially as described [35] but with FGF-8 used instead of FGFb-2. Eighty percent of media was changed every 2 days. After induction for 3 weeks the cells were analysed by immunofluorescence microscopy for the presence of neuronal markers PAX6, NeuN, and TUBB3, confirmation of disease-specific enzymatic deficiencies and lysosomal storage phenotype (increased size of LAMP2+ puncta).

*Neuronal Differentiation* NPCs were differentiated into cortical neurons as previously described [36]. First, NPCs were passage into PO/laminin coated plates in a 1/1 mixture of DMEMF-12/neurobasal (NB) media containing B27, N2, NEAA, BDNF, GDNF, Laminin, dbCAMP, Compound E and TGF-B3 containing 2 µM RI. The following day, media was changed for a 100% NB media with containing the above components. Neurons were then cultured for up to 4 weeks (D28) until fully differentiated and matured, in the presence or absence of 10 µM ACTH_(4-7)_PGP.

### Acute brain slice preparation and electrophysiological recordings

Mice at P60–P64 were deeply anesthetized with a ketamine-xylazine mixture (ketamine: 100 mg/kg, xylazine: 10 mg/kg) and transcardially perfused with 25 ml of ice-cold brain slice cutting solution consisting of (in mM): 250 sucrose, 2 KCl, 1.25 NaH_2_PO_4_, 26 NaHCO_3_, 7 MgSO_4_, 0.5 CaCl_2_ and 10 glucose (pH 7.4, 330–340 mOsm/l). Afterward, animals were immediately decapitated, and their brains were transferred into the Vibratome (VT1000A; Leica Microsystems) specimen tray and immersed in ice-cold brain slice cutting solution to obtain 300 μm thick coronal brain slices. The brain slices were then allowed to recover for at least 45 min in a 34 °C recovery solution composed of (in mM): 118 NaCl, 2.5 KCl, 1 NaH_2_PO_4_, 26 NaHCO_3_, 3 MgSO_4_, 1 CaCl_2,_ and 10 glucose (pH 7.4, 300 mOsm/l), continuously bubbled with carbogen (95% O_2_ and 5% CO_2_). Whole-cell patch clamp recordings of CA1 neurons were made by transferring acute slices from the recovery solution to a recording chamber perfused with artificial cerebrospinal fluid (ACSF) containing 2.5 mM KCl, 118 mM NaCl, 1 mM NaH_2_PO_4_, 26 mM NaHCO_3,_ and 10 mM glucose, continuously bubbled with carbogen (5% CO_2_ and 95% O_2_). Before recording, 2 mM CaCl_2_ and 2 mM MgCl_2_ were added to the ACSF. All recordings were made at 30 °C. Patch pipettes (3–5 MΩ) were filled with internal solution: 115 mM cesium methanesulfonate, 20 mM CsCl, 10 mM HEPES, 2.5 mM MgCl_2_, 0.6 mM EGTA, 4 mM Na_2_ATP, 0.4 mM Na_3_GTP and 10 mM sodium phosphocreatine at pH 7.2 and 290 mOsm. Miniature events were recorded in the presence of 1 μM TTX. Excitatory miniature events were recorded at −60 mV holding potential and inhibitory miniature events at 0 mV holding potential. The first 100 miniature events in each recording were analyzed for amplitude and frequency (Clampfit). AMPA-R and NMDA-R responses in CA1 neurons were evoked by two cluster bipolar stimulation electrodes (FHC #30207) placed above the Schaffer collaterals (150 to 250 mm away from the soma). AMPA-R and NMDA-R currents were evoked at 0.2 Hz in the presence of 100 µM picrotoxin in the ACSF perfusion and recorded while neurons were maintained at − 60 mV and + 40 mV, respectively. For the ratio of AMPA-R vs. NMDA-R responses, we used the AMPA-R peak amplitude at −60 mV from each recording and divided it by its corresponding NMDA-R peak amplitude at +40 mV recorded 80 ms after the stimulation artifact.

### Real-time qPCR

RNA was isolated from snap-frozen brain, kidney and liver tissues using the TRIzol reagent (Invitrogen) and reverse-transcribed using the iScript™ Reverse Transcription Supermix (Bio RAD #1708840) according to the manufacturer’s protocol. qPCR was performed using a LightCycler® 96 Instrument (Roche) and SsoFast™ EvaGreen® Supermix with Low ROX (Bio RAD #1725211) according to the manufacturer’s protocol. RLP32 mRNA was used as a reference control. The forward (F) and reverse (R) primers used in the experiments ale shown in the Table 2.

**Table 2:**
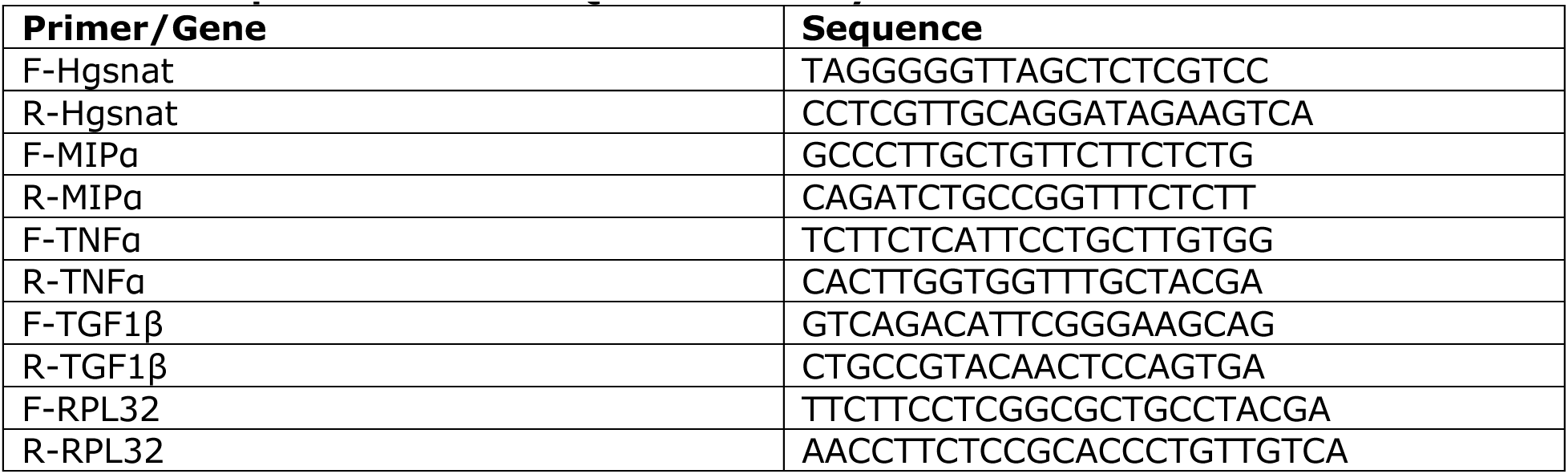
PCR primers used for Q-RT-PCR analysis.

### Immunofluorescence microscopy

Mouse brains were collected from animals perfused with 4% PFA in PBS and post-fixed in 4% PFA in PBS overnight. Brains were cryopreserved in 30% sucrose for 2 days at 4°C, embedded in Tissue-Tek® OCT Compound and stored at −80°C. Brains were cut in 40 µm-thick sections and stored in cryopreservation buffer (0.05 M sodium phosphate buffer pH 7.4, 15% sucrose, 40% ethylene glycol) at −20°C pending immunohistochemistry. Mouse brain sections were washed 3 times with PBS and permeabilized/blocked by incubating in 5% bovine serum albumin (BSA), 0.3% Triton X-100 in PBS for 1 h at room temperature. Incubation with primary antibodies, diluted in 1% BSA, 0.3% Triton X-100 in PBS, was performed overnight at 4°C. The antibodies used in the study and their working concentrations are shown in Table 3.

**Table 3:** Antibodies used in the study and their working concentration.

| Antigen | Host/Target species | Dilution | Manufacturer |
| --- | --- | --- | --- |
| Synapsin-1 | Rabbit anti-mouse | 1:200 (tissues) | Abcam (ab64581) |
| GFAP | Rabbit anti-mouse | 1:300 (tissues) | DSHB (8-1E7-s) |
| Lysosomal-associated membrane protein 2 (LAMP2) | Rat anti-mouse | 1:200 (tissues) | DSHB (ABL-93-s) |
| Lysosomal-associated membrane protein 2 (LAMP2) | Mouse anti-human | 1:100 (cells) | DSHB (H4B4-s) |
| Heparan sulfate (10E4 epitope) | Mouse anti-mouse | 1:200 (tissues) | AMSBIO (F58-10E4) |
| NeuN | Rabbit anti-mouse | 1:400 (cells)<br>1:250 (tissues) | Millipore Sigma (MABN140) |
| CD68 | Rabbit polyclonal to CD68 | 1:200 (tissues) | Abcam (ab125212) |
| GM2 | Mouse humanized | 1:400 (tissues) | KM966 |
| LC3B | Rabbit anti-mouse | 1:200 (tissues) | GeneTex (GTX82986) |
| Brain-derived neurotrophic Factor (BDNF) | Mouse anti-Mouse | 1:2000 (cells)<br>1:200 (tissues) | DSHB (#9-S) |
| Vesicular glutamate transporter1 (VGLUT1) | Rabbit anti-mouse | 1:1000 (cells) and<br>1:200 (tissues) | Abcam (ab104898) |
| Postsynaptic density protein 95 (PSD-95) | Mouse anti-mouse | 1:1000 (cells) and<br>1:200 (tissues) | Abcam (ab99009) |
| Ubiquitin | Mouse | 1:200 (tissues) | Abcam (ab7254) |
| β-Amyloid (D54D2) | Rabbit anti-mouse | 1:200 (tissues) | Cell Signaling (8243S) |
| Recombinant Anti-ATP synthase C antibody (SCMAS) | Rabbit anti-mouse | 1:200 (tissues) | Abcam (ab181243) |
| Anti-O-GlcNAc | Mouse anti-mouse | 1:400 (tissues) | Kerafast (EJH004) |
| Microtubule-associated protein 2 (MAP2) | Chicken anti-mouse | 1:2000 (cells and tissues) | Abcam (ab5392) |
| Myelin Binding Protein (MBP) | Rabbit anti-mouse | 1:500 | Abcam (ab218011) |
| Vesicular gamma aminobutyric acid (GABA) transporter VGAT | Rabbit anti-mouse | 1:1000 | Synaptic Systems (131003) |
| Gephyrin | mouse anti-mouse | 1:1000 | Synaptic Systems (147021) |
| Calreticulin | Rabbit anti-human | 1:200 | Sigma Aldrich (208910) |
| IgG | Goat anti-rabbit, anti-mouse, anti-rat or anti-chicken Alexa Fluor 488-, Alexa Fluor 555- or Alexa Fluor 633-conjugated | 1:400 | Thermo Fisher Scientific |

The mouse brain sections were washed 3 times with PBS and counterstained with Alexa Fluor-labeled secondary antibodies (dilution 1:400) for 2 h at room temperature. After washing 3 times with PBS, the mouse brain sections were treated with TrueBlack® Lipofuscin Autofluorescence Quencher (Biotium, 23007, dilution 1:10) for 1 min, and then again washed 3 times with PBS. To conduct Thioflavin-S staining, brain sections stained with Draq5, primary antibodies against β-amyloid and Alexa Fluor-labeled secondary antibodies were washed 3 times with PBS and incubated in a 0.05% Thioflavin-S (Sigma, T1892) solution in 50% ethanol/water for 10 min protected from light. Then the sections were washed 2 times with 50% ethanol followed by two washes with ddH_2_O. The slides were mounted with Prolong Gold Antifade mounting reagent with DAPI (Invitrogen, P36935) and analyzed using Leica DM 5500 Q upright confocal microscope (10x, 40x, and 63x oil objective, N.A. 1.4). Images were processed and quantified using ImageJ 1.50i software (National Institutes of Health, Bethesda, MD, USA) in a blinded fashion. Panels were assembled with Adobe Photoshop.

### Immunocytochemistry

Cultured mouse neurons at day in vitro (DIV) 21 or iPSC-derived neurons at DIV28 were fixed in 4% paraformaldehyde and 4% sucrose solution in PBS, pH 7.4, for 20 min. The cells were permeabilized with 0.1% Triton-X100 in PBS for 5 min, and non-specific binding sites were blocked with 5% BSA (Wisent) in PBS for 2 h and then, incubated overnight at 4°C with primary antibodies in 1% BSA in PBS (see Table 1 for the source of antibodies and their dilutions). On the following day, neurons were washed 3 times with PBS and labeled with Alexa Fluor 488-, Alexa Fluor-555- or Alexa Fluor 633-conjugated appropriate anti-IgG antibodies (1:1000, all from Thermo Fisher Scientific) for one hour in 1% BSA in PBS at room temperature. Coverslips were washed 3 times again in PBS and mounted on slides using ProLong Gold mounting medium, containing 4′,6-diamidino-2-phenylindole (DAPI; Invitrogen, Cat # P36935), and analyzed by a Leica SP8-DSL or Leica TCS SPE confocal microscopes (× 63 glycerol immersion objectives, N.A. 1.4). Images were processed with Leica Application Suite X (LAS-X) software or Photoshop 2021 (Adobe) and quantified using Fiji-ImageJ 1.50i software (National Institutes of Health, Bethesda, MD, USA). Analysis of images was performed with summation of 9-10 z-stacks separated by 0.5 µm. Soma or axon areas were defined by TUBB3, NEUN, or NF-M staining and, within this area, the appropriate markers were measured establishing a threshold. To obtain LAMP2+ area per neuron, NEUN was used as reference area of the neuron and the image was measured for LAMP2+ puncta while removing background threshold. Synaptic puncta were counted along 10-30 µm segments of neuronal projections at a <10 µm distance from the soma. Quantification was blinded and performed in at least 3 independent experiments.

### Immunoblotting

The cerebral cortical tissues or pellets of cultured cells were homogenized in five volumes of RIPA lysis buffer (50 mM Tris-HCl pH 7.4, 150 mM NaCl, 1% NP-40, 0.25% sodium deoxycholate, 0.1% SDS, 2 mM EDTA, 1 mM PMSF), containing protease and phosphate inhibitor cocktails (Roche, Basel, Switzerland, cat# 4693132001 and 4906837001), using a Dounce homogenizer. The homogenates were kept on ice for 30 min and centrifuged at 13,000 g at 4°C for 15 min. The protein concentration in the lysates was measured using the bicinchoninic acid (BCA) assay. For each sample, 30 µg of protein was loaded and separated on SDS-PAGE polyacrylamide gels either 4–20% gradient precast (Bio-Rad) or prepared. Western blot analyses were performed according to standard protocols using antibodies and conditions listed in the Table 4. Equal protein loading was confirmed by Ponceau S staining. Band intensities were measured using the Image Lab 3.0 software program (Bio-Rad), and normalized to Ponceau S staining.

**Table 4:**
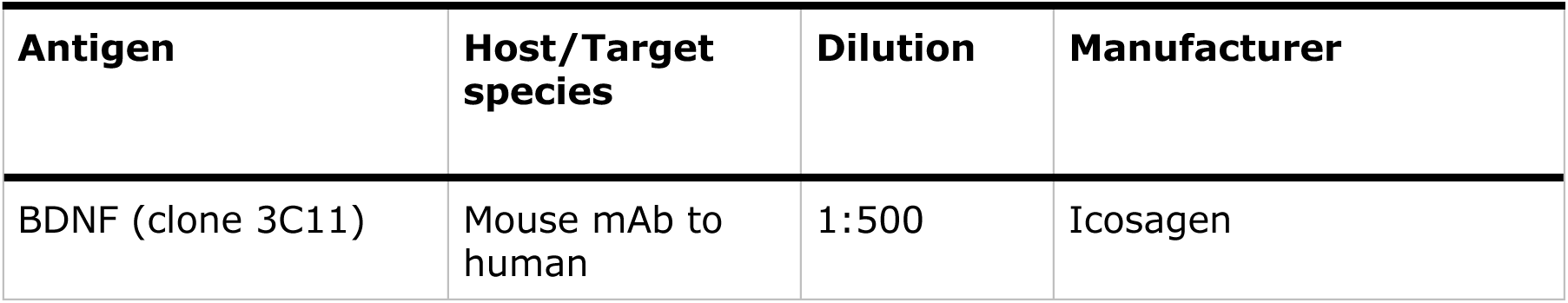
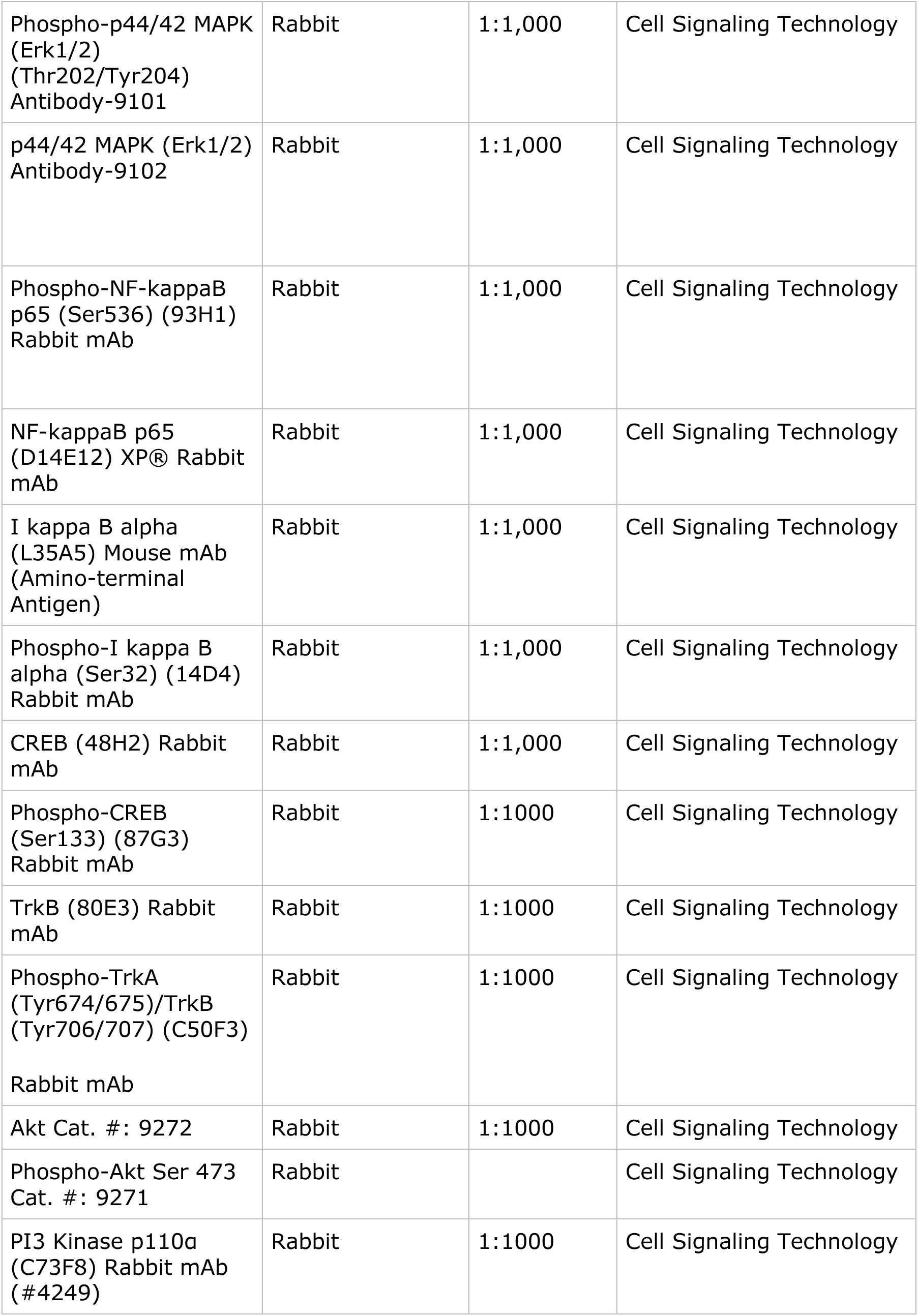
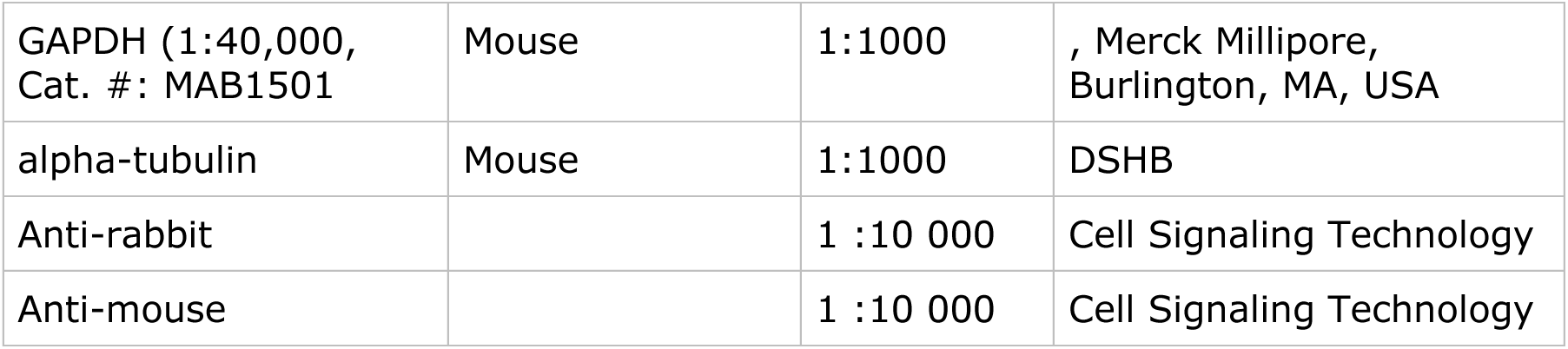
Antibodies used for immunoblots and their working concentrations.

### Analysis of glycosaminoglycans by LC-MS/MS

Analysis of brain glycans was conducted as previously described by Viana *et al*. [10]. Briefly, 30-50 mg of mouse brain tissues were homogenized in ice-cold acetone and centrifuged at 12,000× g for 30 min at 4°C. The pellets were dried, resuspended in 0.5 N NaOH and incubated for 2 h at 50°C. Then the pH of the samples was neutralized with 1 N HCl, and NaCl was added to the reaction mix in a final concentration of 3 M. After centrifugation at 10,000× g for 5 min at room temperature, the supernatants were collected and acidified using 1 N HCl. Following another centrifugation at 10,000× g for 5 min at room temperature, the supernatants were collected and neutralized with 1 N NaOH to a pH of 7.0. The samples were diluted at a ratio of 1:2 with 1.3% potassium acetate in absolute ethanol and centrifuged at 12,000× g and 4°C for 30 min. The pellets were washed with cold 80% ethanol, dried at room temperature, and dissolved in 50 mM Tris-HCl buffer. The samples were further filtered using AcroPrep^TM^ Advance 96-Well Filter Plates with Ultrafiltration Omega 10 K membrane filters (PALL Corporation, USA) and digested with chondroitinase B, heparitinase, and keratanase II, overnight at 37°C. The samples were analysed by mass spectrometry using a 6460 Triple Quad instrument (Agilent technologies) using Hypercarb columns, as described [10].

### ACTH_(4-7)_PGP treatment

Starting from 4 weeks of age, WT C57Bl6 and homozygous *Sgsh^mps3a^* or *Hgsnat^P304L^* male and female mice were randomly divided in treatment and control groups (6-13 mice/sex/genotype/treatment; see Supplementary Table I). The control groups were daily administered with saline (5 µL to each nostril), while for the treatment group, saline was supplemented with 125 μg of ACTH_(4-7)_PGP/mL, which resulted in a dose of ∼50 µg/kg BW/day. The peptide formulation was prepared once, aliquoted and kept frozen at −80°C until use. *Sgsh^mps3a^* mice were studied by TCS, NOR, and CFC behavioural tests and sacrificed. Their blood plasma was collected and their tissues were either snap-frozen or fixed and cryopreserved to analyze CNS pathology as described above [84]. *Hgsnat^P304L^* mice were studied by EPM, OF, YM and NOR behavioral tests at 5 months. Administration of the drug or saline was continued through the days on which the assays were conducted. Then approximately at 6 months 4-5 mice in each group were sacrificed. For the remaining mice, treatment was continued and their behaviour was studied again at the age of 7 months using OF, NOR and YM tests. Starting from the age of 8 months, *Hgsnat^P304L^* treated and untreated mice were daily studied for the signs of urinary retention. When such signs were detected, the mice were sacrificed within 1-2 days. Treated and untreated WT mice were sacrificed at the end of the study, approximately at 11 months of age.

### Preclinical high-field MRI

Six groups of mice were imaged using a preclinical MRI scanner at the Cerebral Imaging Centre of the Douglas Mental Health University Institute (Montreal, QC, Canada). Under terminal anesthesia, animals were perfused with phosphate-buffered saline (PBS) followed by 4% paraformaldehyde (PFA) in PBS. The brains were carefully extracted and immersed in 4% PFA in PBS for 5 hours. Subsequently, the brains were mounted in syringes containing Fomblin oil to facilitate ex vivo MRI scanning. A custom-built solenoid coil designed to fit the syringe was used for imaging. MRI data were acquired on a 7T Bruker scanner using 2D spin-echo sequences with pulsed gradients for diffusion-weighted imaging. The protocol included one b0 and 25 diffusion-weighted images with b-values ranging from 0 to 3000 s/mm^2^ and different diffusion directions for each b-value. Imaging parameters included a field of view of 12 mm × 12 mm, spatial resolution of 0.15 mm × 0.15 mm × 0.4 mm, and TR/TE values of 3300 ms/32 ms. Data reconstruction was performed using the Python-based Diffusion Imaging in Python (DIPY) library, specifically employing the non-linear algorithm for diffusion tensor imaging (DTI) as described by Garyfallidis et al. [85]. On two consecutive slices, regions of interest (ROIs) were manually delineated on the corpus callosum using color-coded fractional anisotropy (FA) maps generated from DTI data.

### Single-cell RNA sequencing

Cell suspensions were prepared as previously described [80,86]. In brief, male and female mice treated with ACTH_(4-7)_PGP or saline for 15 consecutive days between P27 and P42, were anesthetized with ketamine/xylazine (4:1), transcardially perfused with ice-cold aCSF solution containing (in mM): 62.5 NaCl, 70 sucrose, 2.5 KCl, 25 NaHCO_3_, 1.25 NaH_2_PO_4_, 7 MgCl_2_, 1 CaCl_2_, 20 HEPES and 20 glucose. Brains were then quickly removed, transferred to ice-cold aCSF solution, where the hippocampus was removed, and micro dissected. Dissected tissue was then dissociated into a single cell suspension using the Papain dissociation system (Worthington) as per manufacturer’s instructions, with the addition of trehalose as described in [87] Post trituration, cells were then counted using iNCYTO C-Chip hemocytometers, and resuspended at 1000 cells/µl in HBSS Buffer.

The cDNA and the libraries were then generated as described [88] using Chromium Next GEM v3.1 according to the manufacturer guideline (1000268, 10X Genomics; manual: CG000315 RevB). Libraries were sequenced on an Illumina NovaSeq with a S1 flow cell and the 10x Genomics Cell Ranger software was used to process sequencing data. QC filtering was performed in Seurat v4 [89] on each sample to remove low quality cells and potential doublets prior to merging. The merged data was normalized and scaled using Seurat’s SCTransform function [90], followed by principal component analysis (PCA). The data was then integrated using the Harmony package[91] to eliminate a potential batch effect. Dimension reduction using Uniform Manifold Approximation and Projection (UMAP) [92] and clustering was subsequently performed. The resulting clusters were annotated based on known cell type-specific gene expression (**Figure S14**). Clusters corresponding to cell types of interest were then subset and re-clustered, and differential gene expression analysis performed with Seurat’s FindMarkers function using a Wilcoxon Rank test.

### Statistical analysis

Statistical analyses were performed using Prism GraphPad 9.3.0. software (GraphPad Software San Diego, CA). The normality for all data was verified using the D’Agostino & Pearson omnibus normality test. Significance of the difference was determined using t-test (normal distribution) or Mann-Whitney test, when comparing two groups. One-way ANOVA or Nested ANOVA tests, followed by Tukey or Dunnett multiple comparison tests (normal distribution), or Kruskal-Wallis test, followed by Dunn multiple comparisons test, were used when comparing more than two groups. Two-way ANOVA followed by Tukey post hoc test was used for two-factor analysis. A *P*-value of 0.05 or less was considered significant.

---

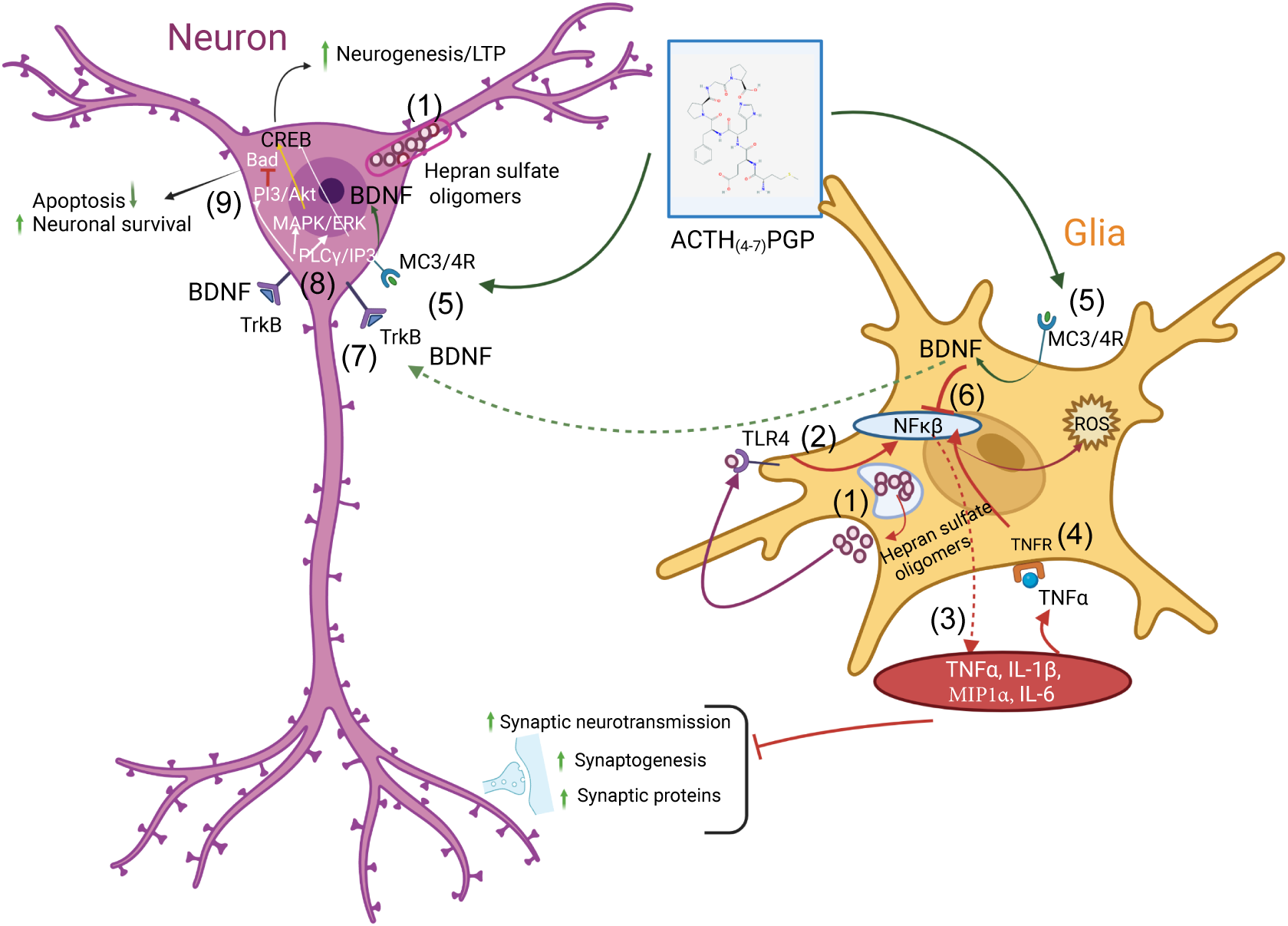
Proposed mechanism underlying ACTH_(4-7)_PGP anti-inflammatory and neuroprotective action in the MPS III brain. HS and HS-derived oligosaccharides accumulate in glial cells and neurons, where they bloc axonal transport to the synaptic terminals **(1)**. These materials are released by exocytosis, act on TLR4/TNFR receptors and trigger the TNFR/NF-kB pathway **(2)** causing activation of glia and the release of cytokines TNFα, IL-1β, IL-6, and MIP-1α **(3)**. These proinflammatory cytokines induce neuronal damage, synaptic defects and reduce neurogenesis. The glial-secreted TNFα binds to TNFR, creating a positive feedback loop that potentiates the initial neuroinflammation response **(4)**. ACTH_(4-7)_PGP acts of MC3/4R receptors in neurons and glia and induces BDNF expression **(5)**. In glia, BDNF inhibits NF-kB activity **(6)** thus blocking the neuroinflammatory response. BDNF produced by neurons and/or BDNF secreted by glial cells act in neurons through the TrkB receptor **(7)**, activating MAPK/ERK kinases and PLCg/IP3 signaling pathways **(8)** which, in turn, activates CREB, leading to increased neurogenesis and enhanced long-term potentiation (LTP). Additionally, BDNF/TrkB signaling activates PI3K/Akt pathway which inhibits the pro-apoptotic protein Bad, increasing neuronal survival **(9)**.

