## Supplemental figures and tables for "Synthetic analogue of adrenocorticotropic hormone, ACTH_(4-7)_PGP delays neurological manifestations in diseases of mucopolysaccharidosis III spectrum by reducing neuroinflammation and rescuing neurotransmission, synaptogenesis, and axonal demyelination"

### Supplementary materials

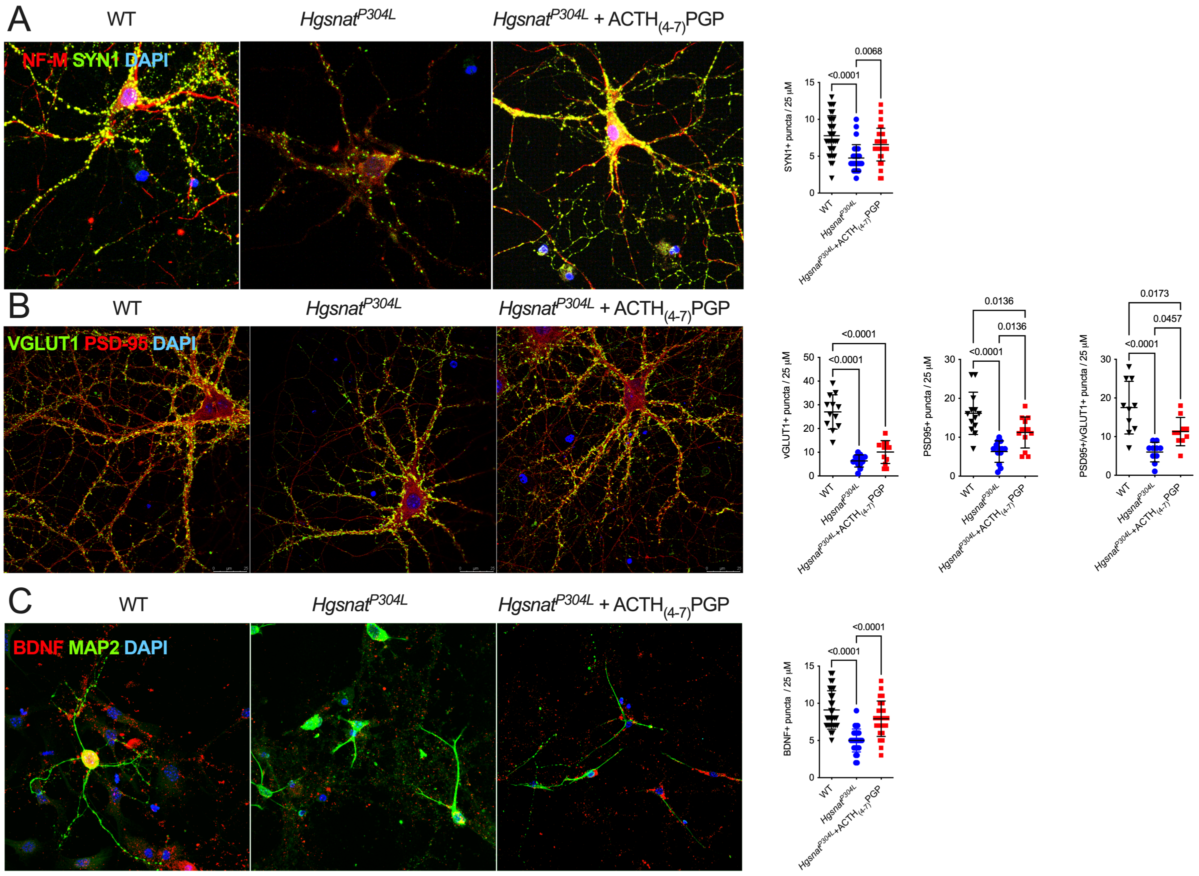


**Figure S1**. **Untreated hippocampal neurons of *Hgsnat*$^{P304L}$ mice show drastic reduction of SYN1+ puncta, VGLUT1+ puncta in juxtaposition with PSD-95+ puncta and BDNF+ puncta, compared to WT cells; the *Hgsnat*$^{P304L}$ neurons treated with ACTH$_{(4-7)}$PGP do not display these deficits.**

Panels show representative images of DIV24 hippocampal primary neurons from WT mice and from *Hgsnat*$^{P304L}$ mice treated or untreated with ACTH$_{(4-7)}$PGP, and labeled with antibodies against **(A)** the neuronal axon marker, NF-M (red) and SYN1 (green), **(B)** VGLUT1 (green) and PSD-95 (red) or **(C)** dendritic marker, MAP2 (green) and BDNF (red). DAPI (blue) was used to label nuclei. The scale bars equal 25 µm. The graphs show quantification of SYN1+, BDNF+, VGLUT1+, PSD95+ puncta and VGLUT1+/PSD95+ puncta in juxtaposition. of neuronal projections by ImageJ software. Individual values (puncta/25 µm), means and SD from 3 biological replicates (10-15 cells) are shown. P-values were calculated using nested one-way ANOVA and Tukey post hoc test.

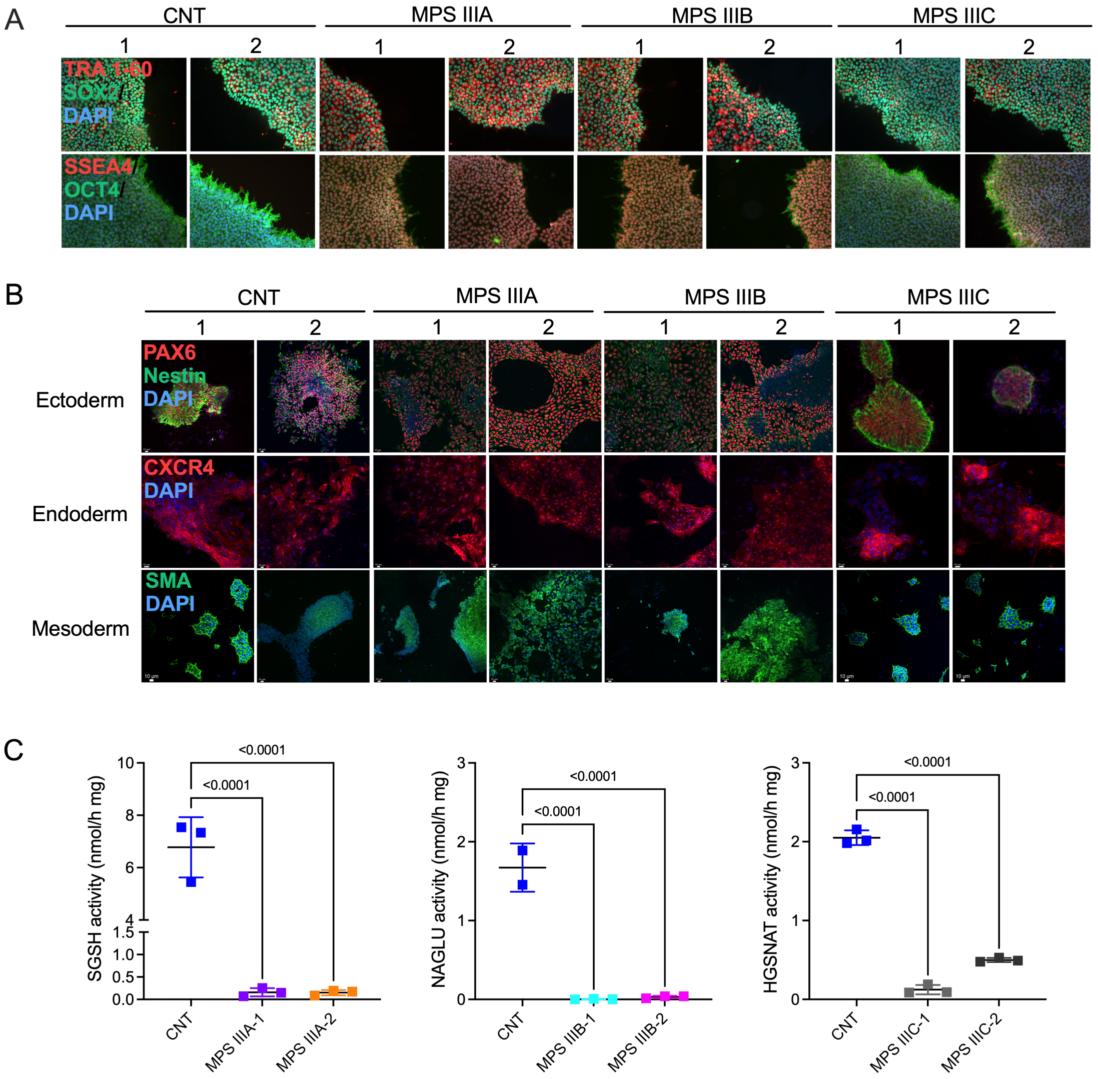


**Figure S2. Generation and characterization of control and MPS III iPSC lines.** **(A)** Representative confocal fluorescent microscopic images for iPSC colonies of healthy control individuals (CNT-1 and CNT-2), MPS IIIA (MPS IIIA-1 and MPS IIIA-2), MPS IIIB (MPS IIIB-1 and MPS IIIB-2), and MPS IIIC (MPS IIIC-1 and MPS IIIC-2) patients labeled for pluripotency and fidelity markers TRA 1-60 (red) and SOX2 (green) or SSEA4 (red) and OCT4 (green). The scale bar represents 10 μm. **(B)** Representative images of CNT, MPS IIIA, MPS IIIB, and MPS IIIC iPSC labeled for trilineage differentiation markers specific for the three primary germ layers, Ectoderm (Nestin, green and PAX6, red), Endoderm (CXCR4, red), and Mesoderm (SMA, green) after culturing with the respective STEMdiff™ media**.** Scale bars equal 10 μm. In all panels, DAPI (blue) was used to label nuclei. **(C)** iPSCs lines of MPS III patients show characteristic primarily enzymatic deficits of SGSH activity for MPS IIIA lines, NAGLU activity for MPS IIIB lines, and HGSNAT activity for MPS IIIC lines. Graphs show individual results, means and SD. N=2-3 independent cultures for MPS IIIA-C and CNT-1/CNT-2 lines; duplicate measurements were performed for each experiment. P values were calculated with one-way ANOVA with Tukey post hoc test.

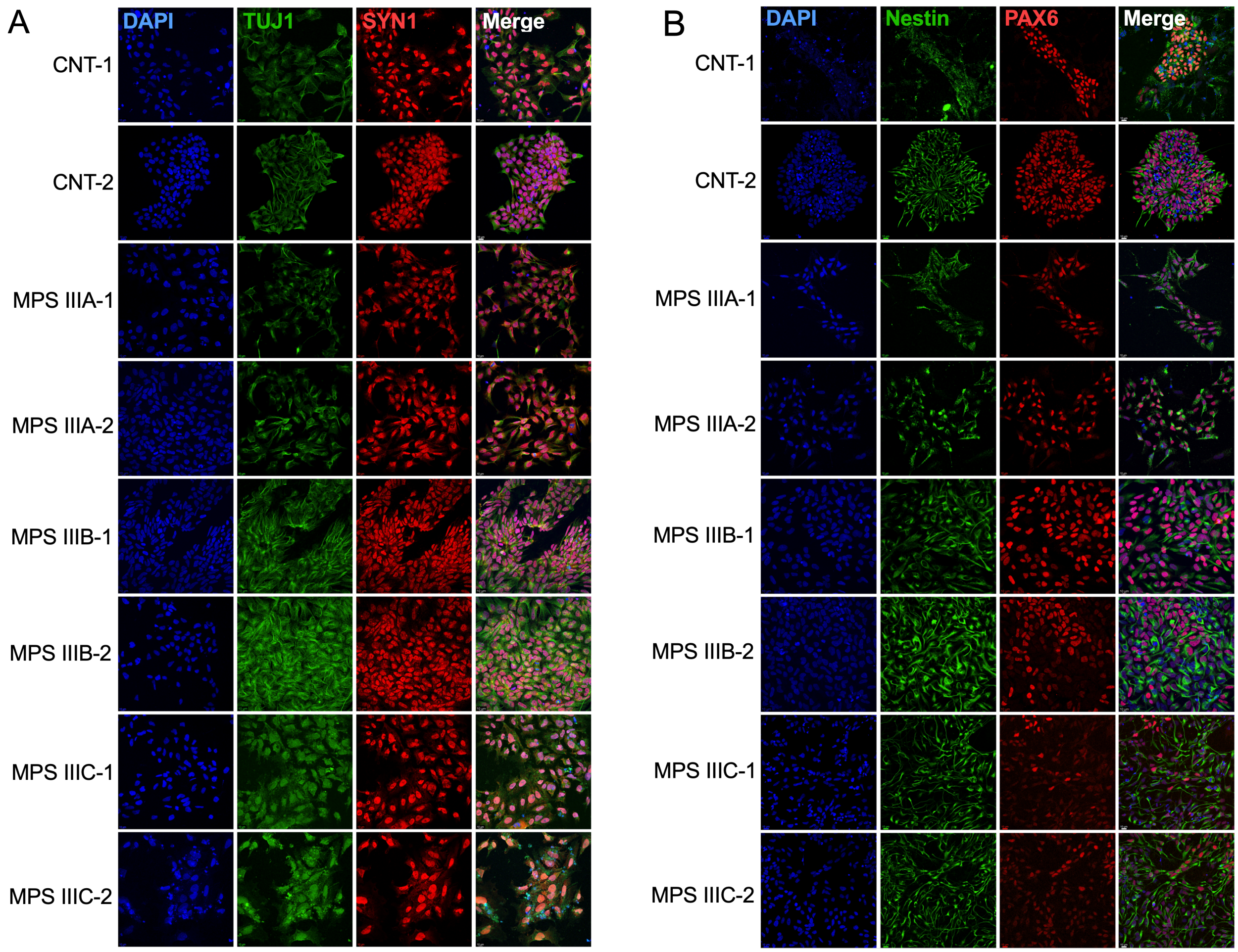


**Figure S3. iPSC-derived NPCs from MPS IIIA, IIIB, and IIIC patients show expression of synaptic markers and neuronal fidelity markers.**
**(A)** Representative confocal fluorescent microscopic images of NPCs from healthy controls (CNT-1 and CNT-2) and MPS III patients after 3 weeks of cortical induction labeled for the neuronal synaptic markers, Tuj1 (green) and Syn1 (red). **(B)** Representative images of NPCs labeled for the NPC fidelity markers, Nestin (green) and Pax6 (red). DAPI (blue) was used to detect nuclei. Scale bars equal 10 μm.

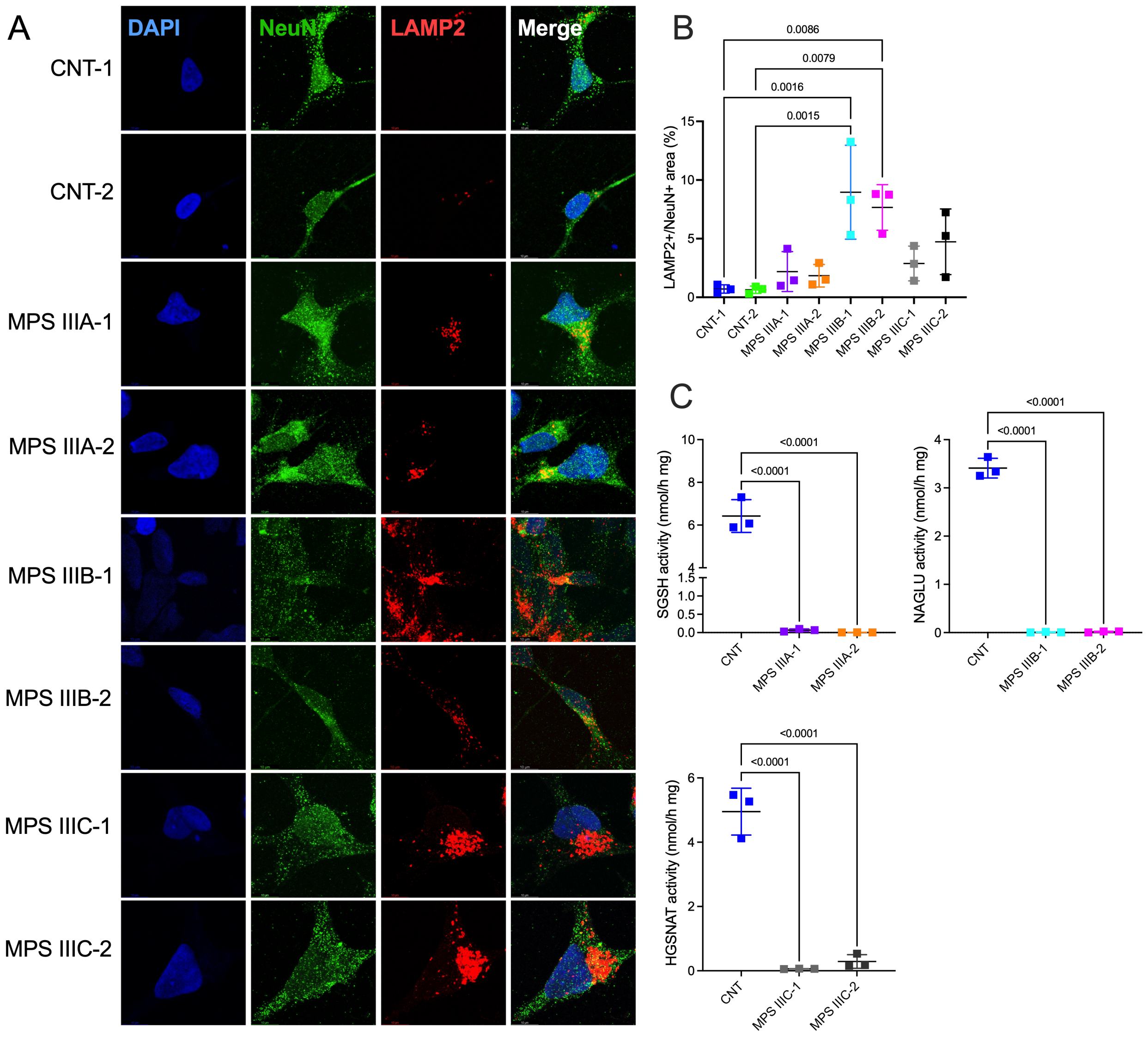


**Figure S4. NPCs of MPS IIIA, IIIB and IIIC patients retain their primary enzymatic deficits and show increased size/abundance of lysosomal LAMP2+ puncta. (A)** Representative images of NPCs from healthy controls (CNT-1 and CNT-2), MPS IIIA, IIIB, and IIIC patients labeled with antibodies against the neuronal soma marker NeuN (green) and the lysosomal marker LAMP2 (red). DAPI (blue) was used to detect nuclei. Scale bars equal 10 µm. **(B)** The graph shows quantification of LAMP2+ area (normalized for NeuN+ area) using ImageJ software. Data show individual results, means and s.e.m. from three independent cultures (5 images were analysed for each culture). P-values were calculated using nested one-way ANOVA and Tukey post hoc test. **(C)** NPCs cells from MPS III patients show characteristic enzymatic deficits of SGSH activity for MPS IIIA, NAGLU activity for MPS IIIB, and HGSNAT activity for MPS IIIC lines. Graphs show individual results, means and SD. N=2-3 independent cultures for MPS IIIA-C and CNT-1/CNT-2 lines; duplicate measurements were performed for each experiment. P values were calculated with one-way ANOVA with Tukey post hoc test.

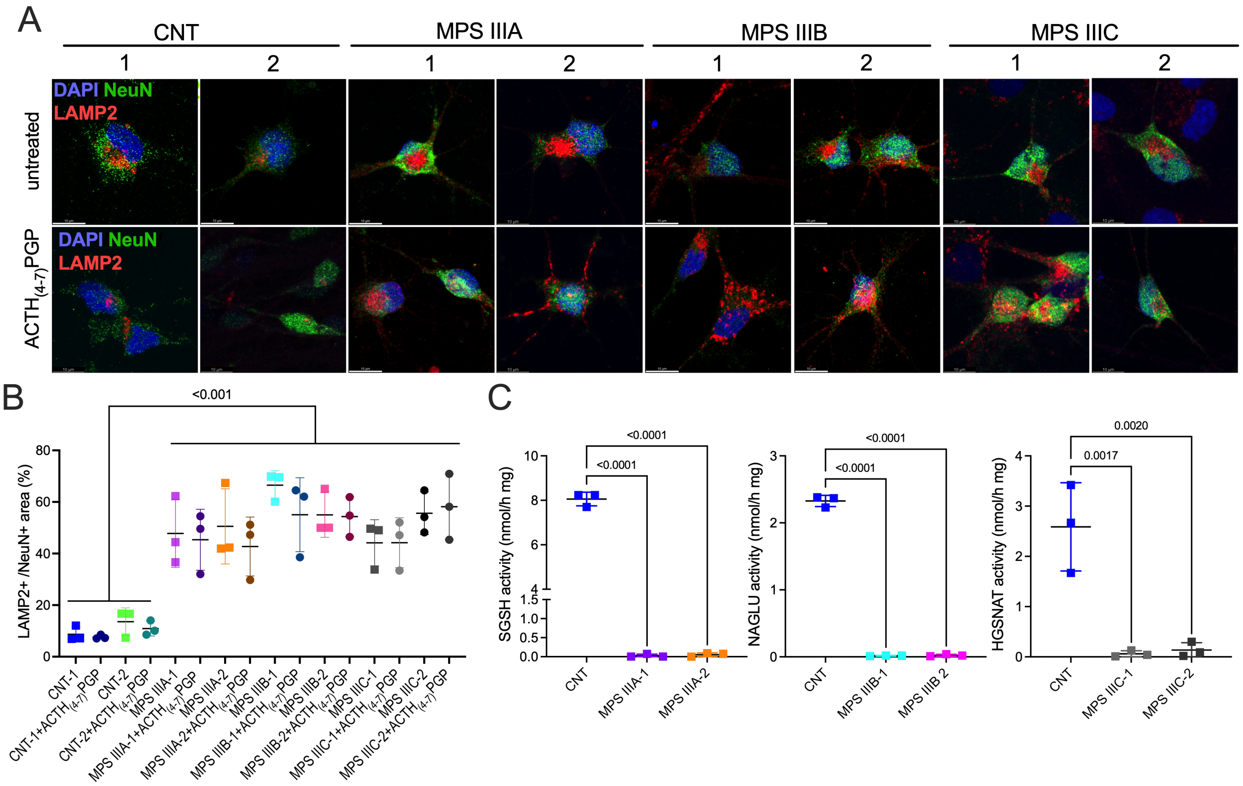


**Figure S5. iPSC-derived cortical neurons of MPS IIIA, MPS IIIB and MPS IIIC patients retain their primary enzymatic deficits and show increased size/abundance of lysosomal LAMP2+ puncta unaffected by ACTH$_{(4-7)}$PGP treatment.**

**(A)** iPSC-derived cortical neurons from MPS III patients show characteristic primarily enzymatic deficits (SGSH activity for MPS IIIA lines, NAGLU activity for MPS IIIB lines, and HGSNAT activity for MPS IIIC lines). Data show individual results, means and s.e.m. from three independent cultures (duplicate measurements were performed for each experiment). P-values were calculated using one-way ANOVA and Tukey post hoc test. **(B)** Representative images of iPSC-derived cortical neurons from healthy controls (CNT-1 and CNT-2) and MPS IIIA, IIIB, and IIIC patients, treated or untreated with ACTH$_{(4-7)}$PGP and labeled with antibodies against the neuronal soma marker NeuN (green) and LAMP2 (red). DAPI (blue) was used to detect nuclei. Scale bars equal 10 µm. **(C)** Quantification of LAMP2+ area from ACTH$_{(4-7)}$PGP-treated and non-treated neurons using ImageJ software. Graph shows data means and s.e.m. from three independent experiments (5 images were analysed in each experiment). P-values were calculated using one-way ANOVA and Tukey post hoc test.

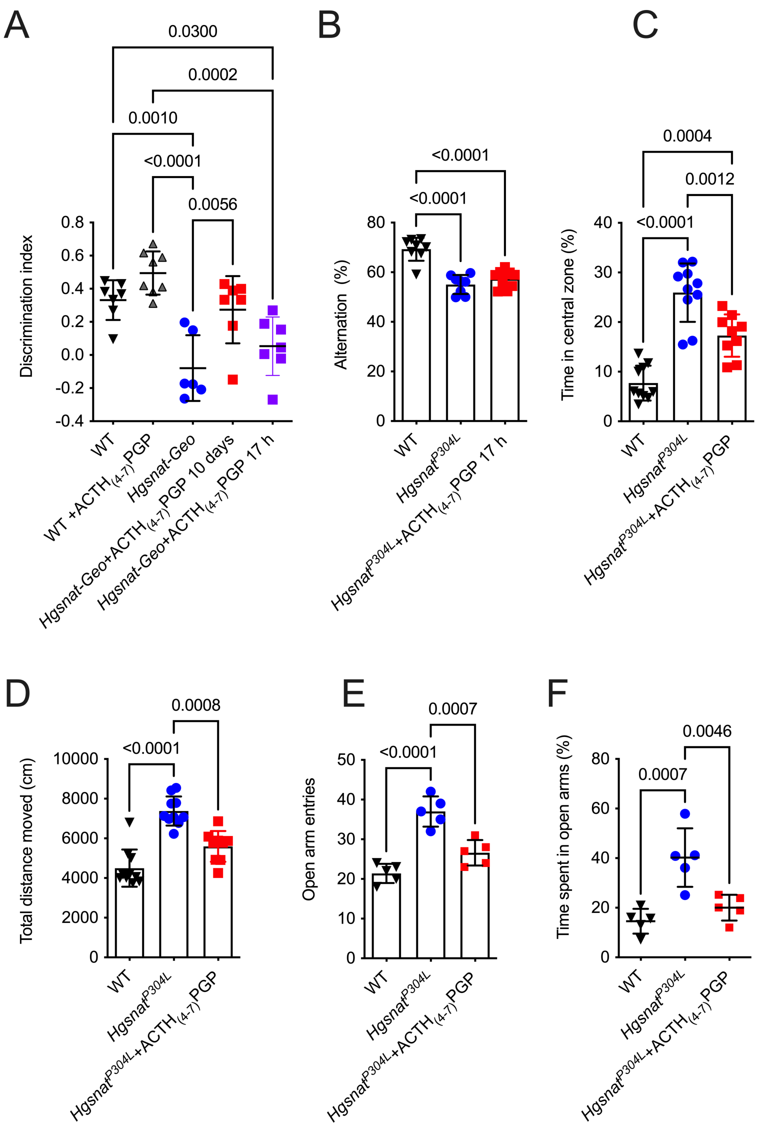


**Figure S6. Short-term treatment with ACTH$_{(4-7)}$PGP partially rescues neurobehavior manifestations in symptomatic MPS IIIC mice.**

**(A)** A significant decrease in discrimination index in the Novel Object Recognition test is observed in 4-month-old *Hgsnat-Geo* mice compared to age-matched WT controls indicating deficits of short-term memory. These deficits are rescued in *Hgsnat-Geo* mice daily treated by intranasal administration of ACTH$_{(4-7)}$PGP at a dose of 50 μg/kg BW for 10 consecutive days preceding the analysis, but not by a single dose of the drug administered 17 h before the test. **(B)** Alternation in the YM test is not increased in *Hgsnat$^{P304L}$* mice which received a single dose of ACTH$_{(4-7)}$PGP 17 h prior to the analysis. **(C-D)** *Hgsnat$^{P304L}$* mice at the age of 4 months show a significant increase in the time spent in the central zone **(C)** and the total distance traveled **(D)** in the Open Field test compared to age-matched WT controls. Both deficits are rescued in the mice, treated with a single 50 μg/kg BW dose of ACTH$_{(4-7)}$PGP 17 h prior to the analysis. **(E-F)** Four-month-old *Hgsnat$^{P304L}$* mice show significant increase in the percent of time spent in open arms and in the number of open arm entries in the Elevated Plus Maze test, compared to age-matched WT controls. These deficits are rescued in mice, treated with a single 50 μg/kg BW dose of ACTH$_{(4-7)}$PGP 17 h prior to the analysis. All graphs show individual data, means and SD (N=5-10 mice/genotype/treatment). P-values were calculated using ANOVA with Tukey post hoc test.

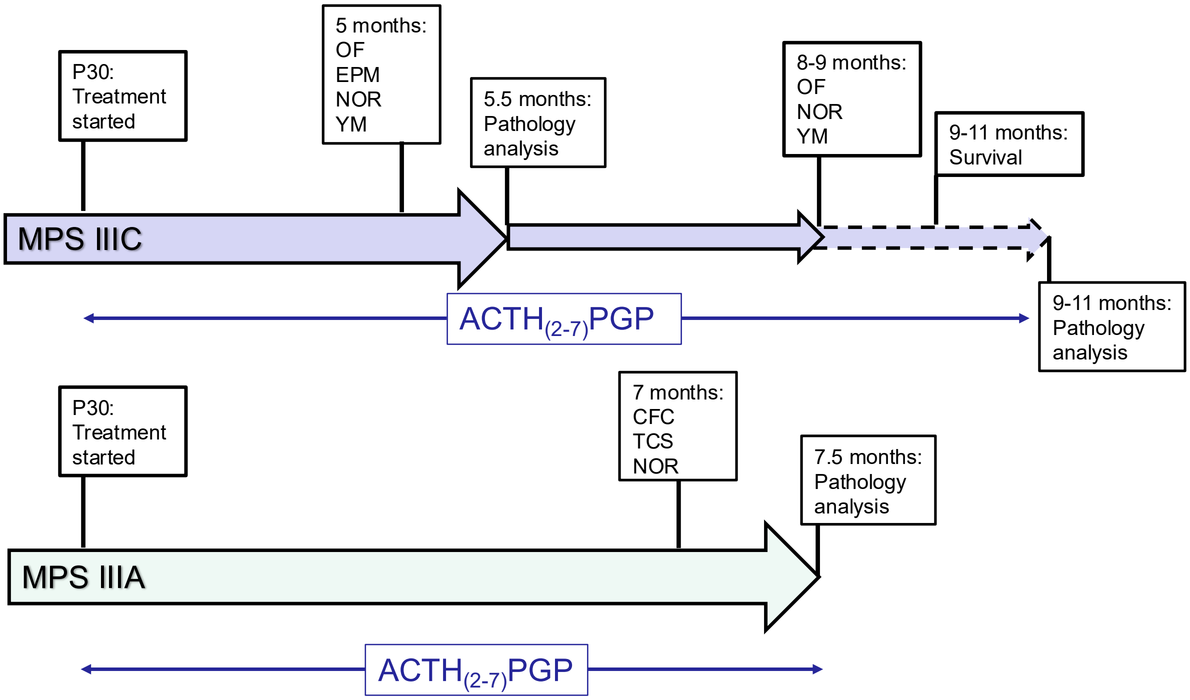


**Figure S7. ACTH$_{(4-7)}$PGP preclinical efficacy study design in MPS IIIA and MPS IIIC mice.** For both MPS IIIC (*Hgsnat*$^{P304L}$) and MPS IIIA (*Sgsh*$^{mps3a}$) models, treatment was started at P30. *Hgsnat*$^{P304L}$ mice were treated between P30 and 5 months of age, when they were assessed by OF (anxiety, fear and hyperactivity), EPM (anxiety, fear), NOR and Y-Maze (short-term memory) tests. While majority of *Hgsnat*$^{P304L}$ and WT mice were sacrificed at the age of ~5.5 months for pathological examination, the treatment was extended for 3-5 mice per sex/treatment/ genotype to test whether the peptide ameliorates behavioural abnormalities at the age of 8 months and increases survival. *Sgsh*$^{mps3a}$ mice were treated between P30 and 7 months of age, studied by NOR, Contextual Fear Conditioning (CFC) and Three-Chamber Sociability (TCS) tests and sacrificed at the age of ~7.5 months for pathological examination.

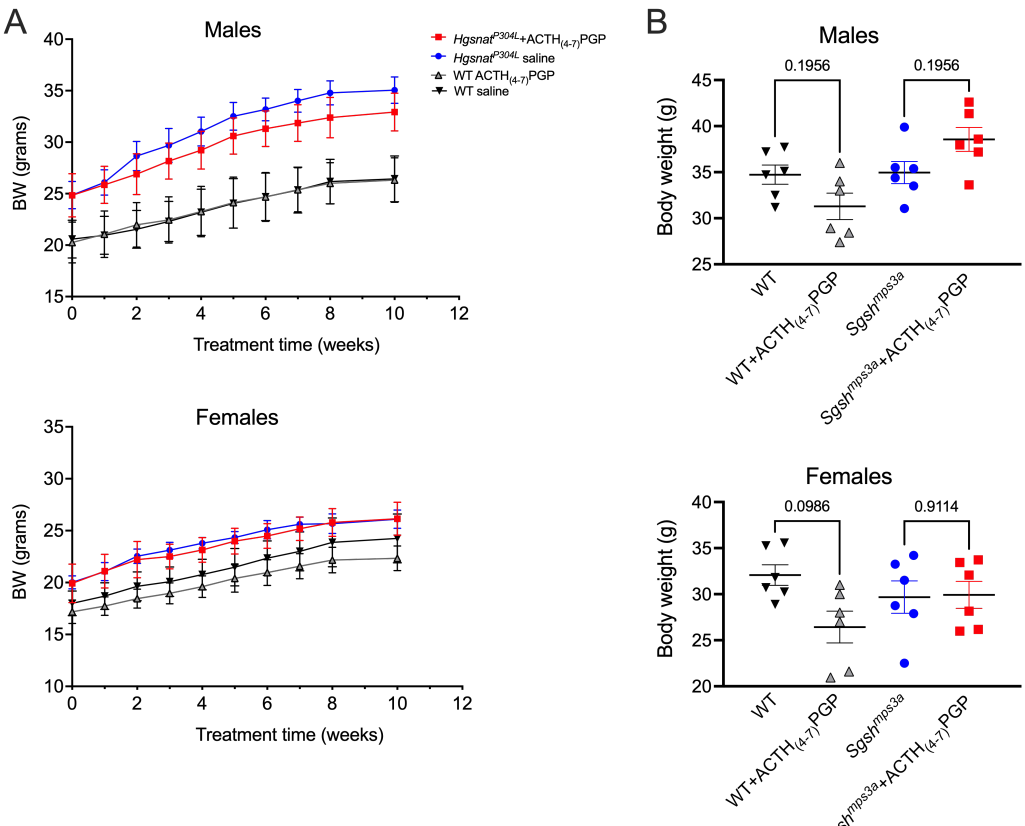


**Figure S8. Mice treated with saline and ACTH$_{(4\text{-}7)}$PGP show similar growth and body weight. (A)** Body weight of MPS IIIC (*Hgsnat*$^{P304L}$) mice was recorded every week starting from the first week of treatment. No difference was observed in the body weight or body weight gain between treated and untreated mice of the same sex and genotype. N=6-10 per sex/ genotype/treatment. Statistical analysis was performed by Repeated measurements two-way ANOVA with Tukey post hoc test. **(B)** Body weight of MPS IIIA (*Sgsh*$^{mps3a}$) mice was measured at 7 month of age before the sacrifice. No difference was observed in the body weight between treated and untreated mice of the same sex and genotype. N=6 per sex/genotype/treatment. Statistical analysis was performed by ANOVA with Holm-Sidak post hoc test.

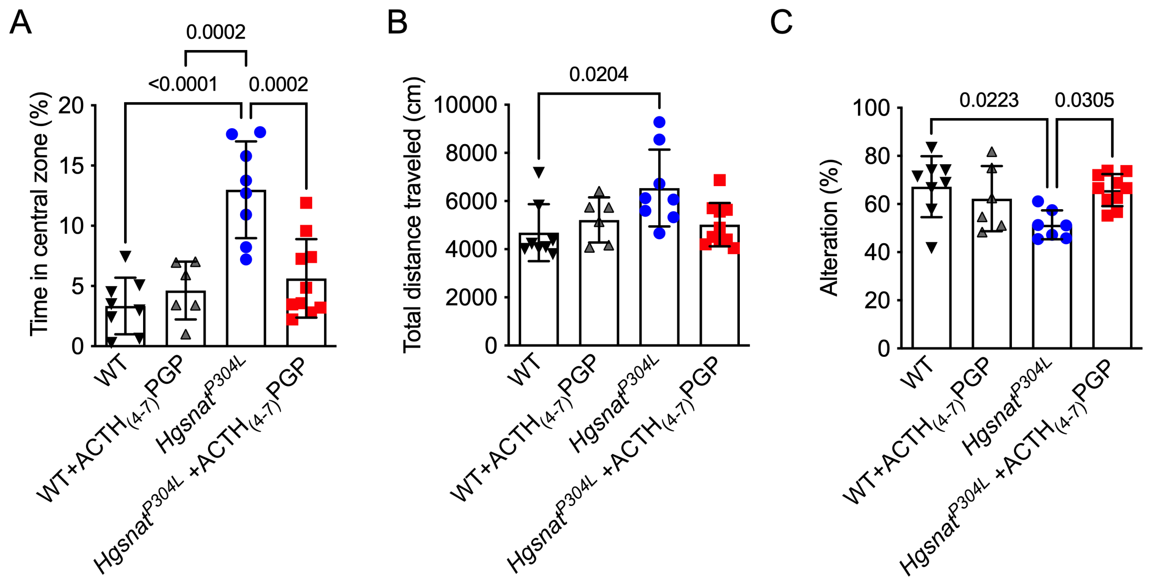


**Figure S9. Chronic daily treatment with ACTH$_{(4\text{-}7)}$PGP ameliorates neurobehavioral abnormalities at the age of 9 months in *Hgsnat$^{P304L}$* mice.**
**(A-C)** *Hgsnat$^{P304L}$* mice treated with saline at the age of 9 months show **(A)** increased time in the central zone in the OF test, **(B)** increased total travel distance in the OF test, and reduced alterations between arms in the Y-Maze test **(C)**. *Hgsnat$^{P304L}$* mice, treated daily with ACTH$_{(4\text{-}7)}$PGP show rescue of all above deficits. Individual results, means and SD for 6-10 mice per group are shown. P values were calculated using one-way ANOVA with Tukey post-hoc test.

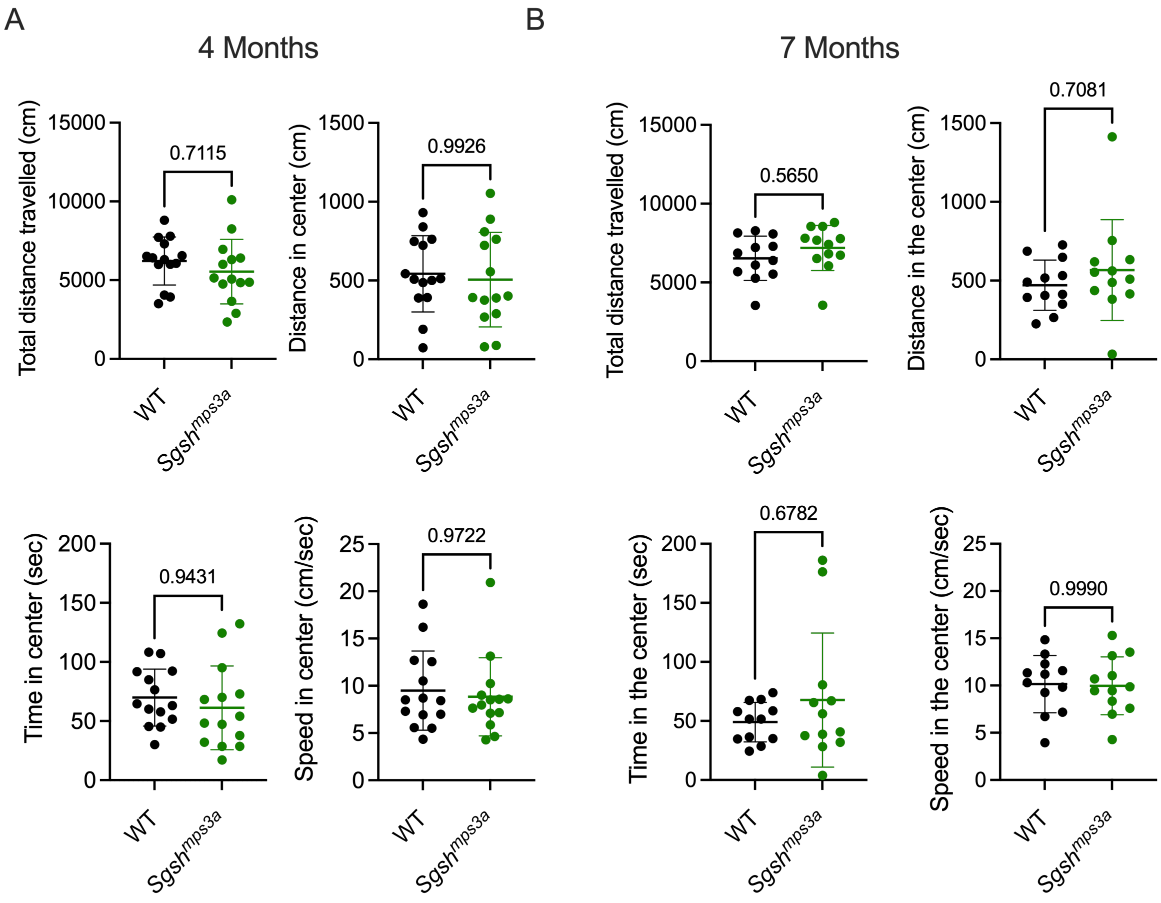


**Figure S10. MPS IIIA *Sgsh$^{mps3a}$* mice do not develop hyperactivity and increased impulsivity at the age of 4 and 7 months**

In the OF test *Sgsh$^{mps3a}$* mice at the age of 4 months **(A)** and 7 months **(B)** show total traveled distance, distance in the central zone, time in the central zone, and speed in the central zone similar to those of the WT mice. Individual results, means and SD for 6-10 male and female mice per group are shown. P values were calculated using two-tailed t test.

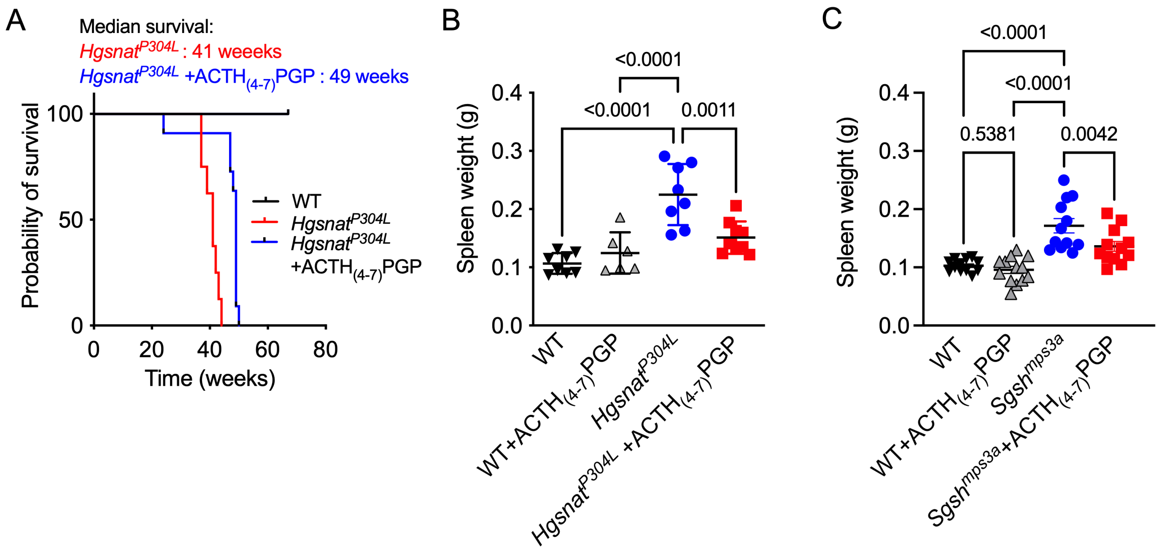


**Figure S11. Chronic daily treatment with $ACTH_{(4\text{-}7)}$PGP increases survival in *Hgsnat$^{P304L}$* mice and reduces splenomegaly in both *Hgsnat$^{P304L}$* and *Sgsh$^{mps3a}$* mice.**

**(A)** Kaplan-Meier plot showing survival of saline-treated (n=8) and $ACTH_{(4\text{-}7)}$PGP-treated *Hgsnat$^{P304L}$* male and female mice (n=10), and their saline-treated (n=8) WT counterparts. Survival rates of treated and untreated *Hgsnat$^{P304L}$* mice are significantly different ($P<0.001$ determined by the Mantel-Cox test). By the age of 43 weeks, all saline-treated *Hgsnat$^{P304L}$* mice had to be sacrificed due to urinary retention, while $ACTH_{(4\text{-}7)}$PGP-treated *Hgsnat$^{P304L}$* mice survived to the average age of 49 weeks. **(B)** Spleen weight of treated and untreated *Hgsnat$^{P304L}$* and WT mice at sacrifice (9.5-11 months of age). Spleen weights are increased in saline-treated *Hgsnat$^{P304L}$* mice compared to WT controls and $ACTH_{(4\text{-}7)}$PGP-treated *Hgsnat$^{P304L}$* mice. **(C)** Spleen weights of treated and untreated *Sgsh$^{mps3a}$* and WT mice at sacrifice (7.5 months of age). Spleen weights are increased in saline-treated *Sgsh$^{mps3a}$* mice compared to WT controls and $ACTH_{(4\text{-}7)}$PGP-treated *Sgsh$^{mps3a}$* mice. Graphs shows individual data, means and SD for 6-12 animals per group. P values were calculated using ANOVA with Tukey post-hoc test.

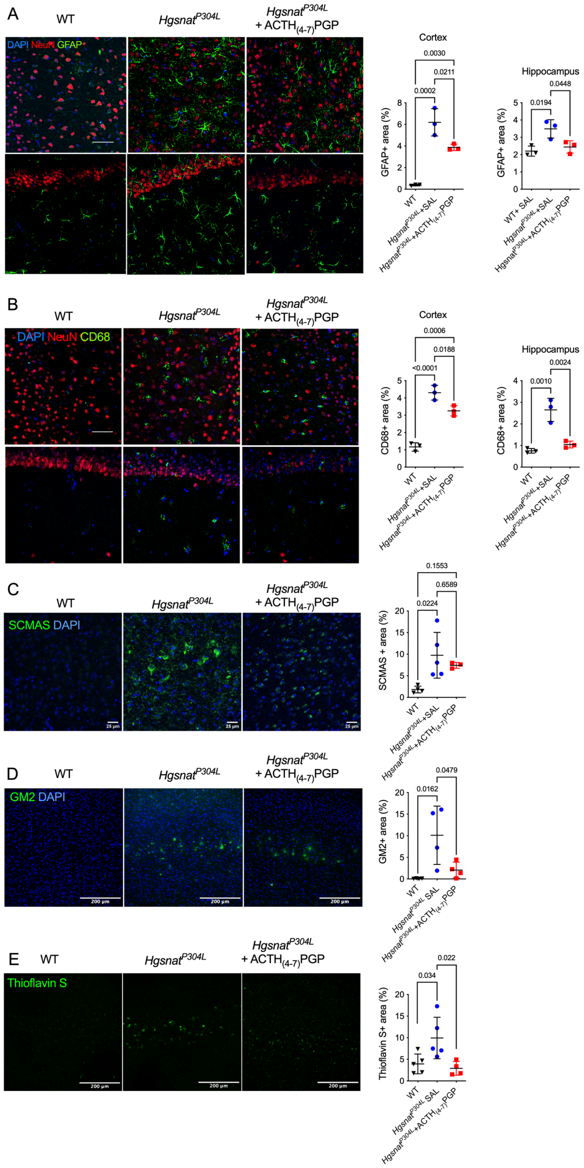

**Figure S12. *Hgsnat$^{P304L}$* mice treated with ACTH$_{(4-7)}$PGP reveal partial rescue of CNS pathology at the age of 10-11 months.**

Levels of activated GFAP+ astrocytes **(A)**, CD68+ microglia **(B),** SCMAS aggregates, **(C)** GM2 ganglioside **(D)** and Tioflavin-S+ materials **(E)** in the brain of 10-11 month-old *Hgsnat$^{P304L}$* mice chronically treated with ACTH$_{(4-7)}$PGP are reduced or show a trend towards reduction as compared to saline-treated *Hgsnat$^{P304L}$* mice. Panels show representative images of brain cortex (layers 4-5) or CA1 areas of hippocampus of WT mice and *Hgsnat$^{P304L}$* mice, treated with saline or ACTH$_{(4-7)}$PGP. The tissues are stained with antibodies against GFAP (green) and NeuN (red) **(A)**, CD68 (green) and NeuN (red) **(B),** SCMAS **(C)** and GM2 ganglioside **(D)** or Thioflavin-S **(E)**. In all panels, DAPI (blue) was used as a nuclear counterstain. Scale bar equals 25 µm **(A-C)** or 200 µm **(D)**. The graphs show quantification of fluorescence with ImageJ software. Individual results, means and SD from experiments performed with 3-4 mice per genotype (3 areas/mouse), per treatment are shown. P values were calculated using nested ANOVA with Tukey post hoc test.

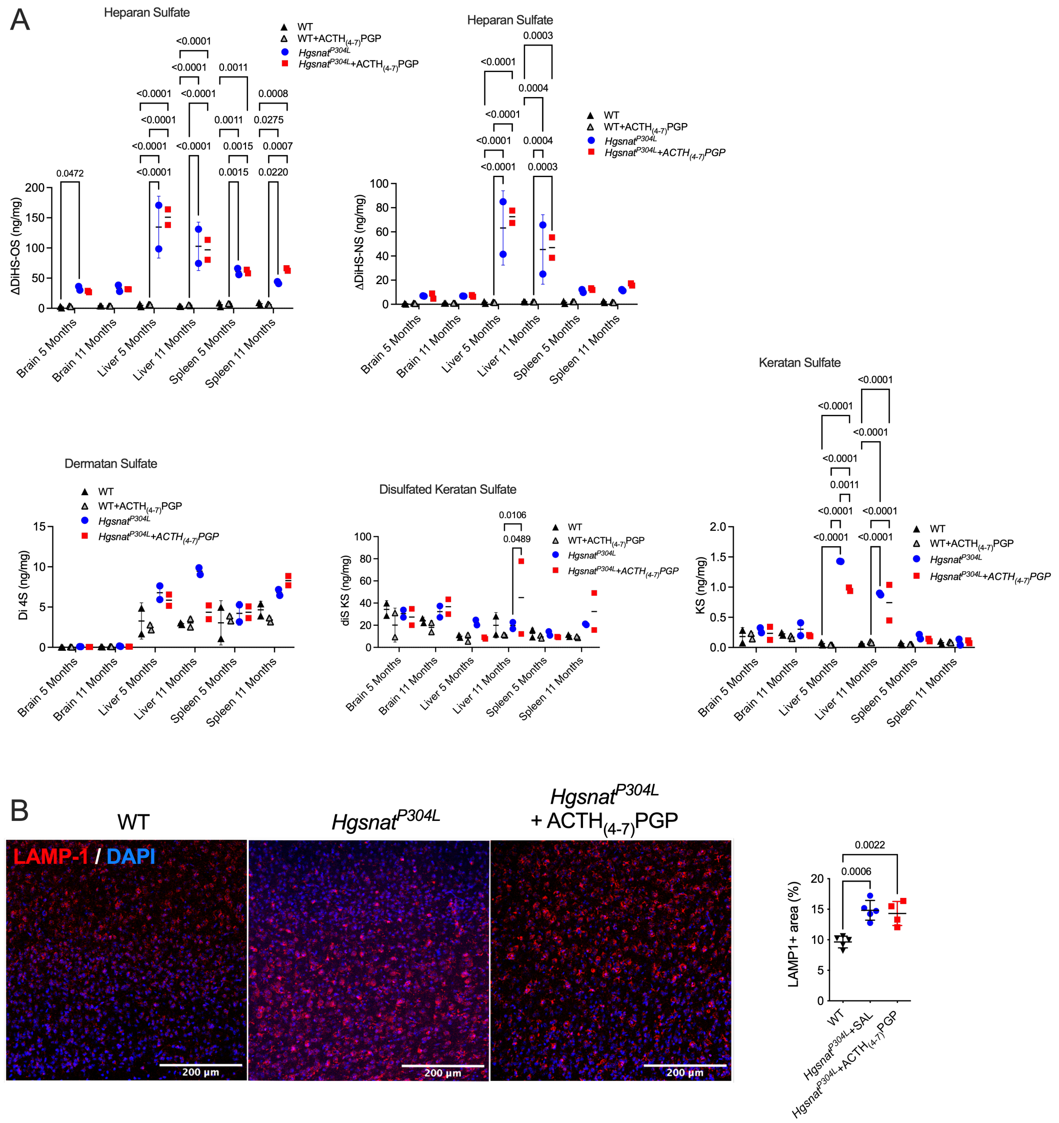


**Figure S13. Levels of HS and LAMP-1 are similar in *Hgsnat$^{P304L}$* mice treated with ACTH$_{(4-7)}$PGP and saline.**

**(A)** Levels of disaccharides produced by enzymatic digestion of HS (ΔDiHS-0S and ΔDiHS-NS), dermatan sulfate (Di 4S), and mono- and disulfated KS were measured by tandem mass spectrometry in dissected brain regions of 5 and 11-month-old WT, and *Hgsnat$^{P304L}$* mice treated with saline or ACTH$_{(4-7)}$PGP. Individual data, and means (±SD) from 2 mice are shown. P values were determined by two-way ANOVA with Dunnett post hoc test. **(B)** Panels show representative images of brain cortex (layers 4-5) from 11-month-old WT mice and *Hgsnat$^{P304L}$* mice, treated with saline or ACTH$_{(4-7)}$PGP. The tissues are stained with antibodies against LAMP-1 (red). DAPI (blue) was used as a nuclear counterstain. Scale bar equals 200 µm. The graph shows quantification of fluorescence with ImageJ software. Individual results, means and SD from experiments performed with 5-4 mice per genotype (3 areas/mouse), per treatment are shown. P values were calculated using nested ANOVA with Tukey post hoc test.

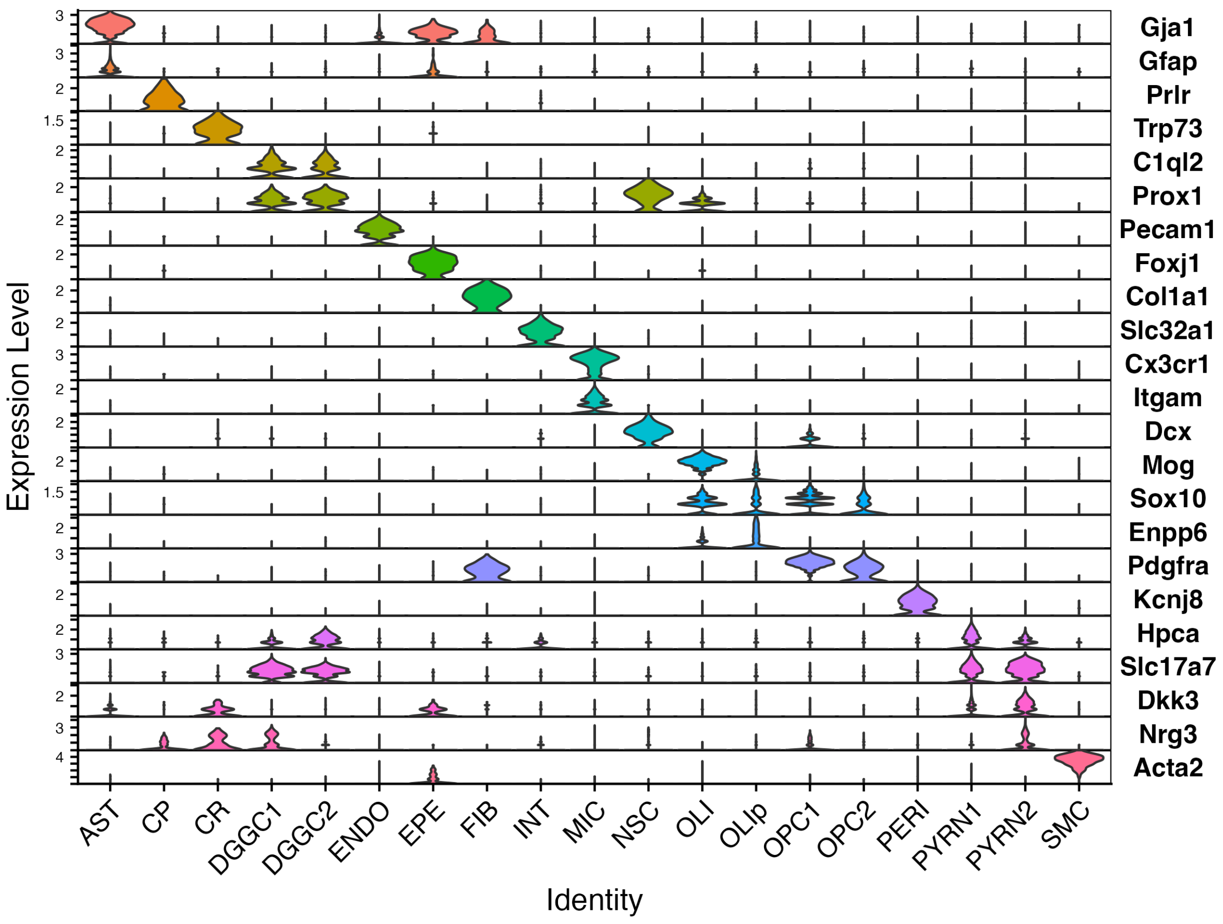


**Figure S14. Violin plots of representative genes used to identify the cell types of each cluster in UMAP from Figure 10A.**

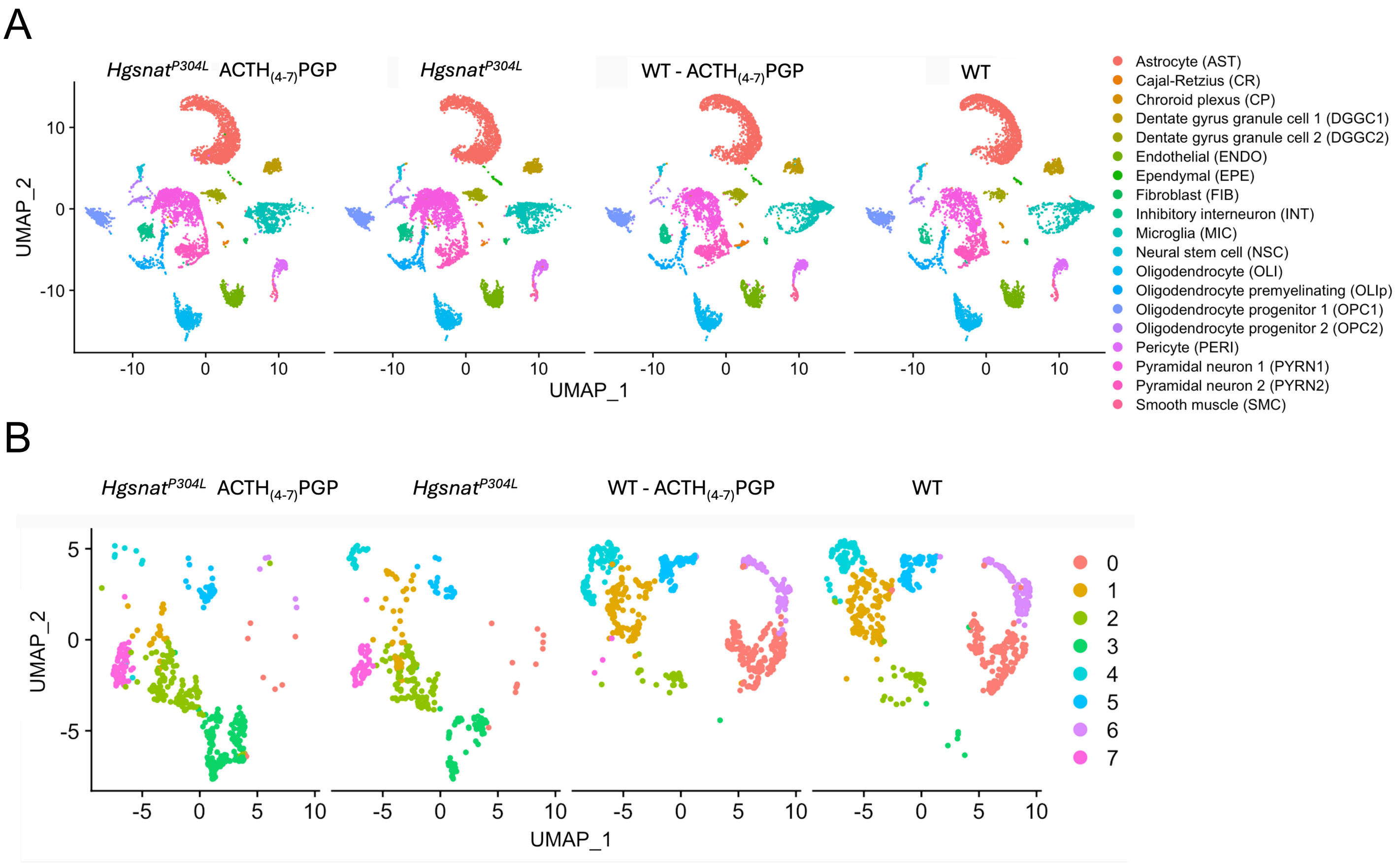


**Figure S15. Uniform manifold approximation and projection (UMAP) plot of single-cell RNA sequencing (scRNA-seq) of WT and *Hgsnat*$^{P304L}$ mice treated with saline or ACTH$_{(4-7)}$PGP.** **(A)** UMAP reveals mostly the same cluster structure for all conditions with minor alterations between the WT and HgsnatP304L mice for pyramidal (PYRN1/PYRN2) and inhibitory (INT) neurons. **(B)** UMAP visualization of PYRN2 cells revealed several subclusters with condition-specific enrichment. Subclusters 0, 1, and 6 were predominantly composed of cells from WT mice, while subclusters 2, 3, and 7 were enriched in cells from the *Hgsnat*$^{P304L}$ group.

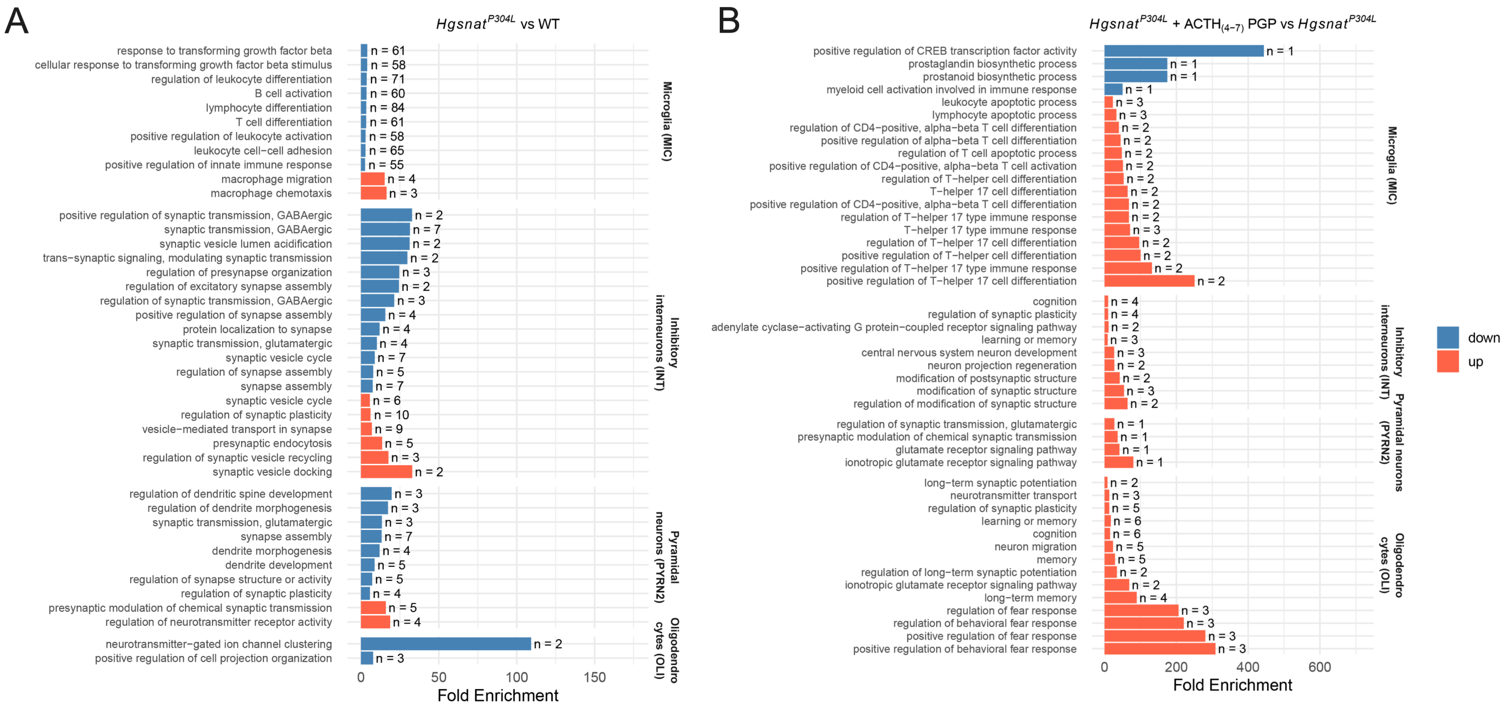


**Figure S16. Selected GO biological process (BP) annotations of the pathways upregulated and downregulated in microglia (MIC), inhibitory interneurons (INT), pyramidal neurons (PYRN2) and oligodendrocytes (OLI) by the disease (A) or treatment (B) for each dataset.** GO terms were selected based on their biological relevance to each cell type and comparison, and enrichment analysis was performed using the GOenrich tool. Bar plots show significantly enriched GO biological processes for two comparisons: *Hgsnat*$^{P304L}$ vs. WT **(A)**, and ACTH$_{(4\text{-}7)}$PGP-treated *Hgsnat*$^{P304L}$ vs. *Hgsnat*$^{P304L}$ **(B)**. The x-axis represents fold enrichment, and the y-axis lists the selected GO terms. The number of differentially expressed genes associated with each term is indicated at the end of each bar. Upregulated terms are shown in red, and downregulated terms in blue. Enrichment analysis was performed on differentially (adjusted $P$<0.05) expressed genes identified within each cluster.

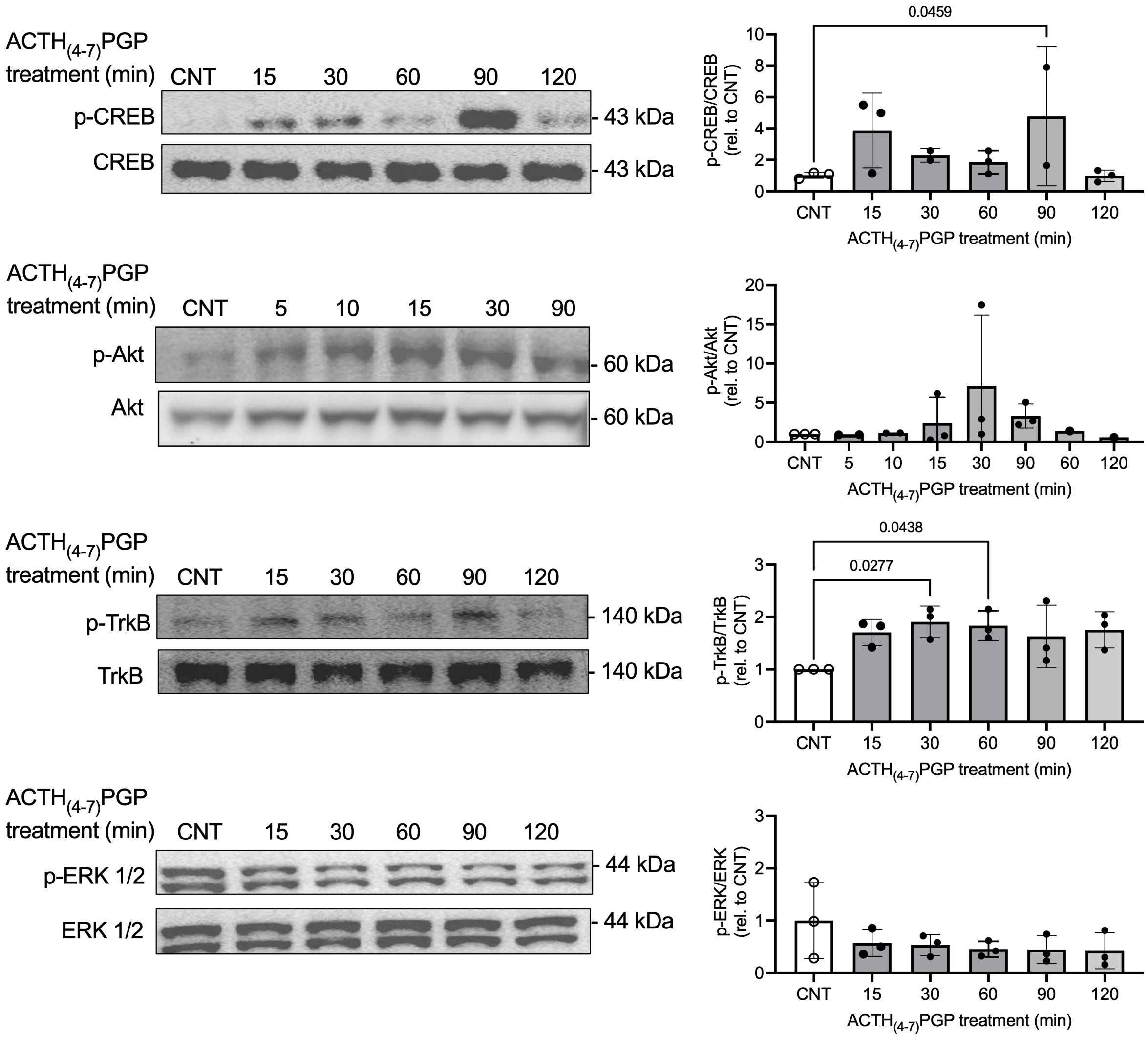


**Figure S17. ACTH$_{(4\text{-}7)}$PGP activates the AKT/CREB/TrkB signaling pathway in differentiated SH-SY5Y neuroblastoma cells.**

Representative immunoblots showing the abundance of CREB and p-CREB, Akt and p-AKT, p-TrkB and TrkB or p-ERK 1/2 and ERK 1/2 in differentiated SH-SY5Y neuroblastoma cells treated with ACTH$_{(4\text{-}7)}$PGP for 15-120 min. Bar graphs show quantification of band intensities (ratio of intensities with phospho-specific and pan-specific antibodies) relative to untreated control cells (CNT). Individual data (n = 2-3) and means ± SD are shown. Statistical analysis was performed using one-way ANOVA followed by Šídák's post hoc test. Drug treatment results in sustained activation of AKT, CREB and TrkB but not ERK 1/2, that shows high basal activation level.

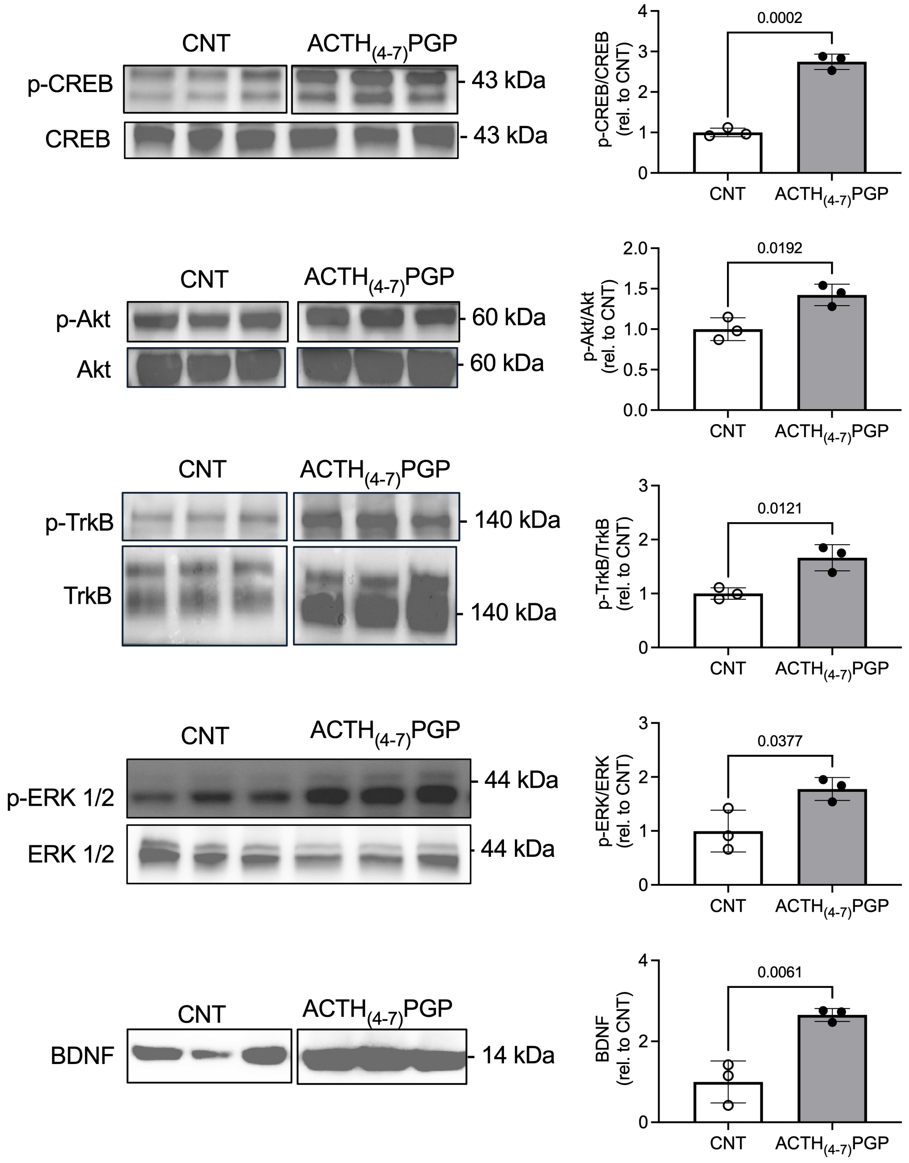


**Figure S18. ACTH$_{(4\text{-}7)}$PGP activates the ERK 1/2/AKT/CREB/TrkB signaling pathway in healthy control iPSC-derived human cortical neurons.**

Representative immunoblots showing the abundance of CREB and p-CREB, Akt and p-AKT, p-TrkB and TrkB, p-ERK 1/2 and ERK 1/2 or mature BDNF in iPSC-derived normal control human cortical neurons cultured in the absence or presence of 10 μM ACTH$_{(4\text{-}7)}$PGP for 28 days. Bar graphs show quantification of band intensities (ratio of intensities with phospho-specific and pan-specific antibodies) relative to untreated control cells (CNT). BDNF band intensities were normalised for total protein detected by Ponceau. Individual data (n = 3) and means ± SD are shown. Statistical analysis was performed using t test. Drug treatment results in sustained activation of AKT, CREB, TrkB and ERK 1/2, as well as in induced BDNF protein levels.
